# GoM DE: interpreting structure in sequence count data with differential expression analysis allowing for grades of membership

**DOI:** 10.1101/2023.03.03.531029

**Authors:** Peter Carbonetto, Kaixuan Luo, Abhishek Sarkar, Anthony Hung, Karl Tayeb, Sebastian Pott, Matthew Stephens

## Abstract

Parts-based representations, such as non-negative matrix factorization and topic modeling, have been used to identify structure from single-cell sequencing data sets, in particular structure that is not as well captured by clustering or other dimensionality reduction methods. However, interpreting the individual parts remains a challenge. To address this challenge, we extend methods for differential expression analysis by allowing cells to have partial membership to multiple groups. We call this grade of membership differential expression (GoM DE). We illustrate the benefits of GoM DE for annotating topics identified in several single-cell RNA-seq and ATAC-seq data sets.

## Background

A key methodological aim in single-cell genomics is to learn structure from single-cell sequencing data in a systematic, data-driven way [1–3]. Clustering [4–7] and dimensionality reduction techniques such as such as PCA [8–10], *t*-SNE [11] or UMAP [12] are commonly used for this aim. Despite the fact that many of these techniques have been applied “out-of-the-box” (with some caveats [13–18]), they have been remarkably successful in revealing and visualizing biologically interesting substructures from single-cell data [7, 19– 29].

Another class of dimensionality reduction approaches that have been used to identify structure from singlecell data are what are sometimes called *parts-based representations*—these approaches include non-negative matrix factorization (NMF) [30–44] and topic modeling [45–56], which also have formal connections [48, 57, 58]. Parts-based representations share some of the features of both a clustering and a dimensionality reduction: on the one hand, they learn a lower dimensional representation of the cells; on the other hand, the individual dimensions (the “parts”) of the reduced representation can identify discrete clusters or discrete subpopulations [59, 60]. However, parts-based representations are more flexible than clustering—the dimensions can also capture other features such as continuously varying cell states.

In this paper, we investigate the question of how to interpret the individual dimensions of a parts-based representation learned by fitting a topic model. (In the topic model, the dimensions are also called “topics”.) For topics that assign observations to discrete clusters, one could apply a standard method for differential expression analysis [61, 62] to compare expression between topics, then annotate these topics by the genes that are differentially expressed. The question, therefore, is what to do with topics that do not assign observations to discrete clusters. To tackle this question, we extend models that compare expression between groups by allowing observations to have *partial membership in multiple groups*. This more flexible differential expression analysis is implemented by taking an existing model and modifying it to allow for partial memberships to groups or topics. This modified model is a “grade of membership” model [63], so we call our new method *grade of membership differential expression* (GoM DE). The idea is that, by generalizing existing methods, we can continue to take advantage of existing elements of differential expression analysis, but now apply them to learn about different types of cell features beyond discrete cell populations.

We describe the GoM DE approach more formally in the next section. Then we evaluate the GoM DE approach in simulations, showing, in particular, that it recovers the same results as existing differential expression analysis methods when the cells can be grouped into discrete clusters. In case studies, we demonstrate how the GoM DE analysis analysis can be used to uncover and interpret a variety of cell features from single-cell RNA-seq and ATAC-seq data sets.

## Results

### Methods overview and illustration

We begin by giving a brief overview of the topic model, then we describe the new methods for annotating topics. To illustrate key concepts, we analyze a singlecell RNA-seq (scRNA-seq) data set obtained from peripheral blood mononuclear cells (PBMCs) [29] that has been used in several benchmarking studies [e.g., 4, 7, 8, 64, 65]. We refer to these data as the “PBMC data.”

#### Learning expression topics from single-cell RNA-seq data

The original aim of the topic model was to discover patterns from collections of text documents, in which text documents were represented as word counts [45, 50, 66–68]. By substituting genes for words and cells for documents, topic models can also be used to learn a reduced representation of cells by their membership in multiple “topics” [47].

When applied to scRNA-seq data generated using UMIs, the topic model assumes a multinomial distribution of the RNA molecule counts in a cell,

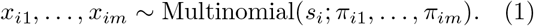

where *s*_*i*_ = *x*_*i*1_ + *· · ·* + *x*_*im*_, and *m* is the number of genes. That is, the number of RNA molecules *x*_*ij*_ observed for gene *j* in cell *i* is a noisy observation of an underlying true expression level, *π*_*ij*_ [8, 69].

For *n* cells, the topic model is a *reduced representation* of the underlying expression,

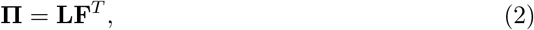

where **Π, L, F** are *n × m, n × K, m × K* matrices, respectively, with entries *π*_*ij*_, *l*_*ik*_, *f*_*jk*_. Each cell *i* is rep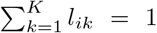, and each “expression topic” is represented by its “grade of membership” in *K* topics, a vector of proportions *l*_*i*1_, …, *l*_*iK*_, such that *l*_*ik*_ *≥* 0, resented by a vector of (relative) expression levels *f*_1*k*_, …, *f*_*mk*_, *f*_*jk*_ *≥* 0. (These are also constrained to sum to 1, which ensures that the *π*_*ij*_’s are multinomial probabilities.) To efficiently fit the topic model to large single-cell data sets, we exploit the fact that the topic model is closely related to the Poisson NMF model [48].

The matrix **L** in (2), which contains the membership proportions for all cells and topics, can be visualized using a “Structure plot”. Structure plots have been used to visualize the results of population genetics analyses [e.g., 70–72], and, more recently, to visualize the topics learned from bulk and single-cell RNAseq data [47].

A Structure plot visualizing the topic model fit to the PBMC data, with *K* = 6 topics, is given in Figure 1. In this data set, the cells have been “sorted” into different cell types which provides a cell labeling to compare against. From the Structure plot, it is apparent that a subset of topics—topics 1, 2 and 3—correspond closely to the sorted subpopulations (B cells, CD14+ monocytes, CD34+ cells). (Indeed, distinctive genes and enriched gene sets identified by the methods described below suggest these same subpopulations; Figure 1C.) Topics 4 and 5, on the other hand, are not confined to a single sorted cell type, and instead appear to capture biological processes common to T cells and natural killer (NK) cells. CD8+ cytotoxic T cells have characteristics of both NK cells and T cells—these are T cells that sometimes become “NK-like” [73]—and this is captured in the topic model by assigning membership to both topics. Topic 6 also captures continuous structure, but, unlike topics 4 and 5, it is present in almost all cells, and therefore its biological interpretation is not at all clear from the cell labeling. More generally, the topics, whether they capture largely discrete structure (topics 1–3) or more continuous structure (topics 4–6), can be thought of as a “soft” clustering [47].

**Figure 1.**
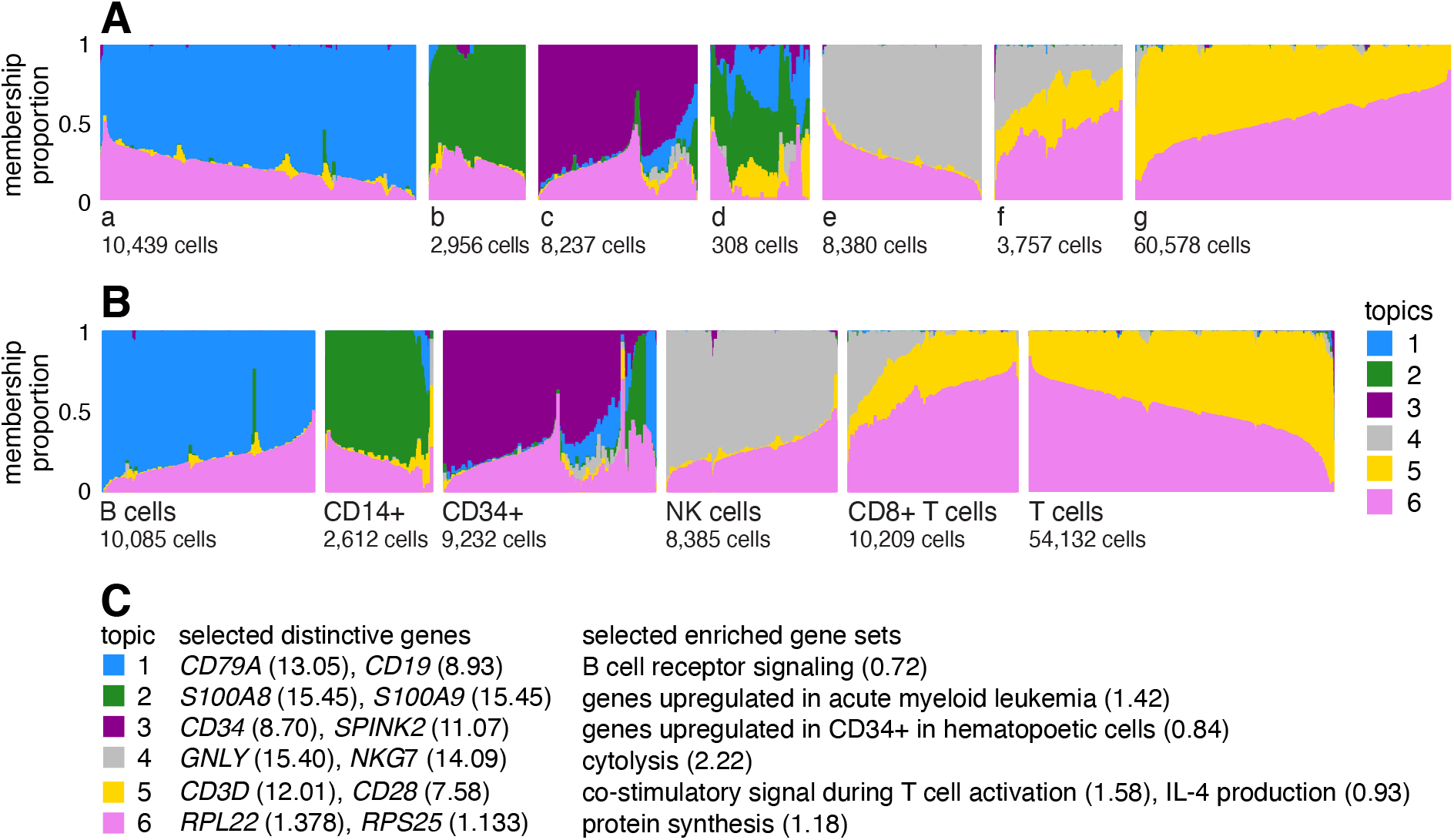
A and B give two views of the topic model fit to the PBMC data [29] (*n* = 94,655 cells, *K* = 6 topics) using Structure plots [70, 71]. Cells are arranged horizontally; bar heights correspond to cell membership proportions. In A, the cells are arranged using the estimated membership proportions only. In B, the cells are grouped by the FACS labels. (The “T cells” label combines all sorted T cell populations other than CD8+ cytotoxic T cells.) In C, the topics are annotated by distinctive genes from the GoM DE analysis (Figure 3) and by enriched gene sets. Numbers in parentheses next to genes give posterior mean l.e. LFCs, and for gene sets they are enrichment coefficients. An enrichment coefficient is an estimate of the expected increase in the LFC for genes that belong to the gene set relative to genes that do not belong to the gene set. Note the groupings a–g in A are intended only to aid visualization. See also Additional file 1: Figure S1 for an alternative visualization.

#### Learning chromatin accessibility topics from single-cell ATAC-seq data

For single-cell ATAC-seq data, the observations *x*_*ij*_ denote the number of reads mapping to region *j* in cell *i*. However, it is common to “binarize” the read counts such that *x*_*ij*_ = 1 when at least one fragment in cell *i* maps to region *j* and *x*_*ij*_ = 0 otherwise.

Using the topic model to analyze (binarized) singlecell ATAC-seq data was first suggested by [49]. Therefore, they implicitly assumed a multinomial model (1) in which the *x*_*ij*_’s are binarized accessibility values instead of UMI counts. A binomial model for binarized accessibility data was proposed in [74]. As we explain in Methods, we view both models as approximations, and under reasonable assumptions the models are similar.

#### Differential expression analysis allowing for grades of membership

Having learned the topics, our aim now is to identify genes that are distinctive to each topic. In the simplest case, the topic is a distinct or nearly distinct cluster of cells, such as topic 1 or 2 in Figure 1.

In the following, we describe methods for analyzing *differences in expression*, but they can also be understood as methods for analyzing *differences in chromatin accessibility*. Therefore, “expression,” “expressed” and “gene” in the descriptions below may be substituted with “accessibility”, “accessible” and “peak” (or “region”).

Consider a single gene, *j*. Provided unmodeled sources of variation are negligible relative to measurement error, a simple Poisson model of expression should suffice:

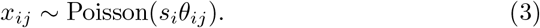

In this model, *θ*_*ij*_ for gene *j* in cell *i* is controlled by the cell’s membership in the cluster: when cell *i* belongs to the cluster, *θ*_*ij*_ = *p*_*j*1_; otherwise, *θ*_*ij*_ = *p*_*j*2_. Under this model, differential expression (DE) analysis proceeds by estimating the log-fold change (LFC) in expression for each gene *j*,

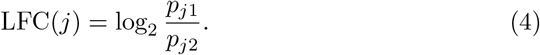

Although simple, this Poisson model forms the basis for many DE analysis methods [75–80].

We now modify the Poisson model (3) in a simple way to analyze differential expression among topics. In a clustering, each cell belongs to a single cluster, whereas in the topic model, cells have *grades of membership* to the clusters [63] in which *l*_*ik*_ is the membership proportion for cluster or topic *k*. Therefore, we extend the model to allow for partial membership in the *K* topics:

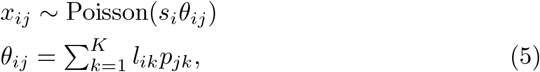

in which the membership proportions *l*_*ik*_ are treated as known, and the unknowns *p*_*j*1_, …, *p*_*jK*_ represent relative expression levels. (A related model is used in C-SIDE [80] to model cell-type mixtures in DE analysis of spatial transcriptomics data.) Note that *p*_*jk*_ will be similar to, but not the same as, *f*_*jk*_ in the topic model because the DE analysis is a gene-by-gene analysis, whereas the topic model considers all genes at once. The standard Poisson model (3) is recovered as a special case of (5) when *K* = 2 and all membership proportions *l*_*ik*_ are 0 or 1.

Recall, our aim is to identify genes that are *distinctive* to each topic. To this end, we estimate the *least extreme LFC* (l.e. LFC), which we define as

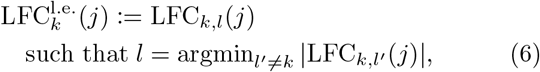

in which LFC_*k,l*_(*j*) is the *pairwise LFC*,

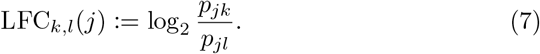

In words, the l.e. LFC for topic *k* is the LFC comparing topics *k* and *l*, in which *l* is chosen to be topic that results in the smallest (“least extreme”) change. By this definition, a “distinctive gene” is one in which its expression is significantly different from its expression in *all other topics*. (Note the l.e. LFC reduces to the standard LFC (4) when *K* = 2.) We then annotate topics by the distinctive genes. The estimation of l.e. LFCs and computation of related posterior statistics is described in Methods.

To illustrate what the least extreme LFC does and does not do, consider the following toy example with *K* = 10 topics (Figure 2). Gene 1 has high expression in topic 1 and low expression in the other topics. Therefore, all the pairwise LFCs for topic 1 are large, LFC_1,*k*_(1) = log2(100), *k* = 2, …, 10, and this results in an l.e. LFC for topic 1 of log_2_(100) *≈* 6.6. So gene 1 is a distinctive gene for topic 1. Next consider gene 2, which has high expression in topics 1 and 2 and low expression in the other topics. For gene 2, the pairwise LFCs for topic 1 are mostly large, LFC_1,*k*_(2) = log2(100), *k* = 3, …, 10, except for LFC_1,2_(2) = 0. So the l.e. LFC for topic 1 is zero and, as a result, gene 2, although potentially helpful for interpreting topic 1, is not a distinctive gene for topic 1.

**Figure 2.**
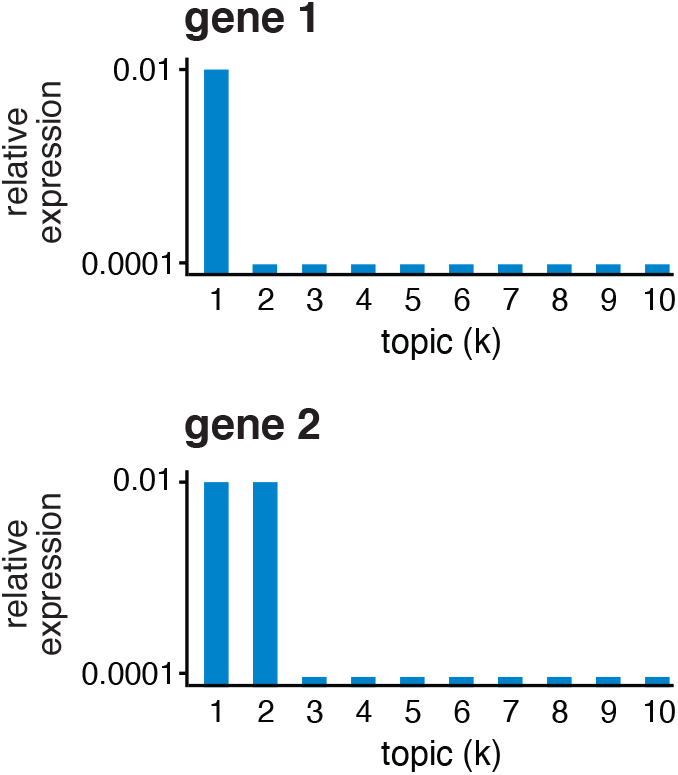
Toy example illustrating the least extreme LFC. Gene 1 has high expression in topic 1 and low expression in the other topics; *p*_11_ = 0.01, *p*_1*k*_ = 0.0001, k = 2, …, 10. Gene 2 has high expression in topics 1 and 2 and low expression in the other topics; *p*_21_ = *p*_22_ = 0.01, *p*_2*k*_ = 0.0001, k = 3, …, 10.

#### Illustration of GoM DE analysis in PBMC data set

To illustrate, we applied the GoM DE analysis to the topic model shown in Figure 1, and visualized the results in “volcano plots” (Figure 3). We then used the GoM DE results (Additional file 2: Table S1) to perform gene set enrichment analysis (Additional file 3: Tables S3, S4).

**Figure 3.**
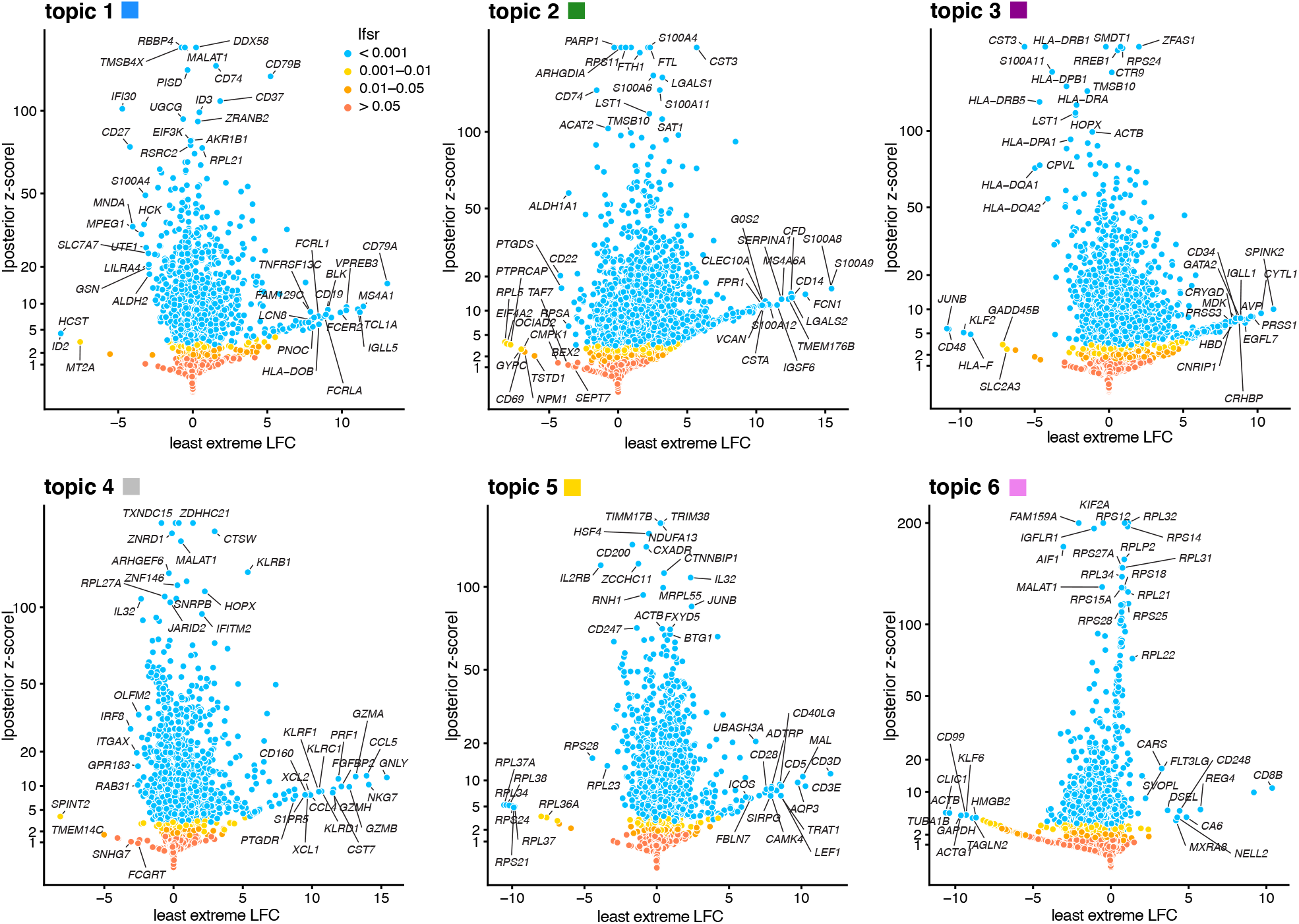
GoM DE analysis of the PBMC data using the topic model shown in Figure 1. The volcano plots show posterior mean estimates of the l.e. LFC vs. posterior *z* -scores for 17,055 genes. The posterior *z* -score is defined as the posterior mean l.e. LFC divided by the posterior standard error. Genes are colored according to the local false sign rate (*lfsr* ) [81]. A few genes with extreme posterior *z* -scores are shown with smaller posterior *z* -scores so that they fit within the y-axis range. See also the detailed GoM DE results (Additional file 2: Table S1), detailed GSEA results (Additional file 3: Table S3, S4) and the interactive volcano plots (Additional file 4).

For the topics that closely correspond to cell types, the GoM DE analysis, as expected, identified genes and gene sets reflecting these cell types. For example, topic 1 corresponds to FACS B cells, and is characterized by overexpression of *CD79A* (posterior mean l.e. LFC = 13.05) and enrichment of B cell receptor signaling genes (enrichment coefficient = 0.72). Topic 2 corresponds to myeloid cells and is characterized by overexpression of *S100A9* (l.e. LFC = 15.45) and enrichment of genes down-regulated in hematopoietic stem cells (enrichment coefficient = 0.90).

The close correspondence between topics 1 and 2 and FACS cell types (B cells, myeloid cells) provides an opportunity to contrast the GoM DE analysis with a standard DE analysis of the FACS cell types (Figure 4). This is not a perfect comparison because the topics and FACS cell populations are not exactly the same, but the LFC estimates correlate well (Figure 4A, B). This comparison illustrates to two key differences:

**Figure 4.**
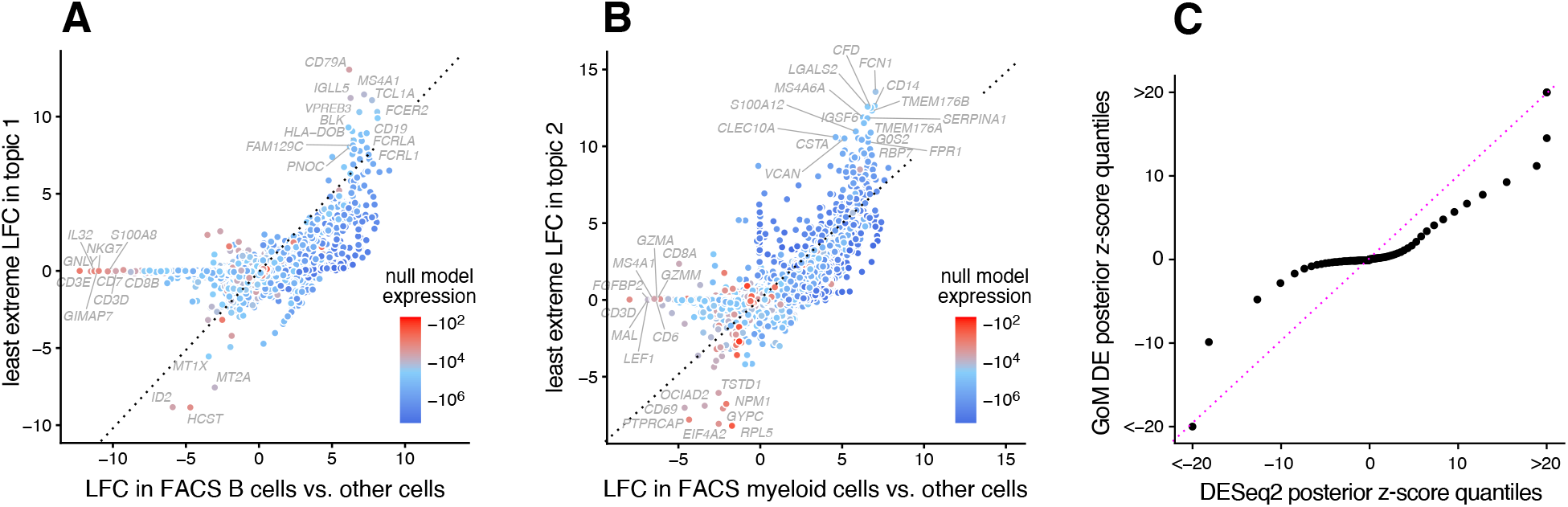
GoM DE analysis vs. DESeq2 analysis in PBMC data. Panels A and B compare differential expression in topics 1 and 2 (Figure 1) with their closely corresponding FACS cell populations. Genes are only shown if the posterior *z* -score was greater than 2 in magnitude in at least one of the DE analyses. Genes are colored by the “null model” expression rate. The Q-Q plot (Panel C) compares the overall distribution of posterior *z* -scores for B cells and myeloid cells (x-axis) and for topics 1 and 2 (y-axis). For better visualization of quantiles near zero, posterior *z* -scores larger than 20 in magnitude are shown as 20 or -20. Analysis of differential expression among the 6 FACS cell populations was performed using DESeq2 [79, 83].

1. Many more l.e. LFCs are driven toward zero in the GoM DE analysis (Figure 4C), so the l.e. LFCs more effectively draw attention to the “distinctive genes” (Figure 4A, B). This includes genes that are *distinctively underexpressed* such as *ID2* in B cells [82].
2. The GoM DE analysis yields much larger LFC estimates of the cell-type-specific genes. This is because the topic model isolates the biological processes (topics 1 and 2) related to cell type while removing background biological processes (topic 6) that do not relate to cell type.

Other topics capture more continuous structure, such as topics 4 and 5 (Figure 1). Although the GoM DE analysis of these topics is not comparable to a standard DE analysis, many of the the distinctive genes and gene sets suggest NK and T cells, which are precisely the FACS-labeled cells with greatest membership to these topics: for example, for topic 4, overexpression of *NKG7* (posterior mean l.e. LFC = 14.09), enrichment of cytolysis genes (enrichment coefficient = 2.22); for topic 5, overexpression of *CD3D* (l.e. LFC = 12.01), enrichment co-stimulatory signaling during T-cell activation (enrichment coefficient = 1.58).

Topic 6 captures continuous structure and is present in almost all cells, so knowledge of the FACS cell types is not helpful for understanding this topic. Still, the GoM DE results for topic 6 show a striking enrichment of ribosome-associated genes (Figure 3, Additional file 3: Tables S3, S4). (These ribosomal protein genes also account for a large fraction of the total expression in the cells [5].) This ability to annotate distinctly non-discrete structure is a distinguishing feature of the grade-of-membership approach, and below we will show more examples where this feature contributes to understanding of the cell populations.

### Evaluation of DE analysis methods using simulated data

Having illustrated the features of this approach, we now evaluate the methods more systematically in simulated expression data sets. We began our evaluation by first considering the case of two groups in which there is no partial membership to these groups; that is, when the cells can be separated into two cell types. The GoM DE analysis should accommodate this special case, and should compare well with existing DE analysis methods. We compared with DESeq2 [79] and MAST [84], both popular methods that have been shown to be competitive in benchmarking studies [61, 62, 85] (and are included in Seurat [25]).

To compare the ability of these methods to discover differentially expressed genes, we simulated RNA molecule count data for 10,000 genes and 200 cells in which 98% of cells were attributed to a single topic, with roughly the same number of cells assigned to each of the two topics (with membership proportions of 99% or greater). Note that although half the simulated genes had different expression levels in the two topics, most of these expression differences were small, and therefore the methods were not expected to identify most expression differences. This mimics the typical situation in gene expression studies whereby most expression differences are small. Molecule counts were simulated using a Poisson measurement model so that variation in expression across cells was due to either measurement error or true differences in expression levels between the two groups. For all DE analyses, we took group/topic assignments to be known so that incorrect assignment of cells to topics was not a source of error. Other aspects of the simulations were chosen to emulate molecule count data from scRNA-seq studies (see Methods). We repeated the simulations 20 times, and summarized the results of the DE analyses in Figure 5 (also Additional file 1: Figures S2, S3).

**Figure 5.**
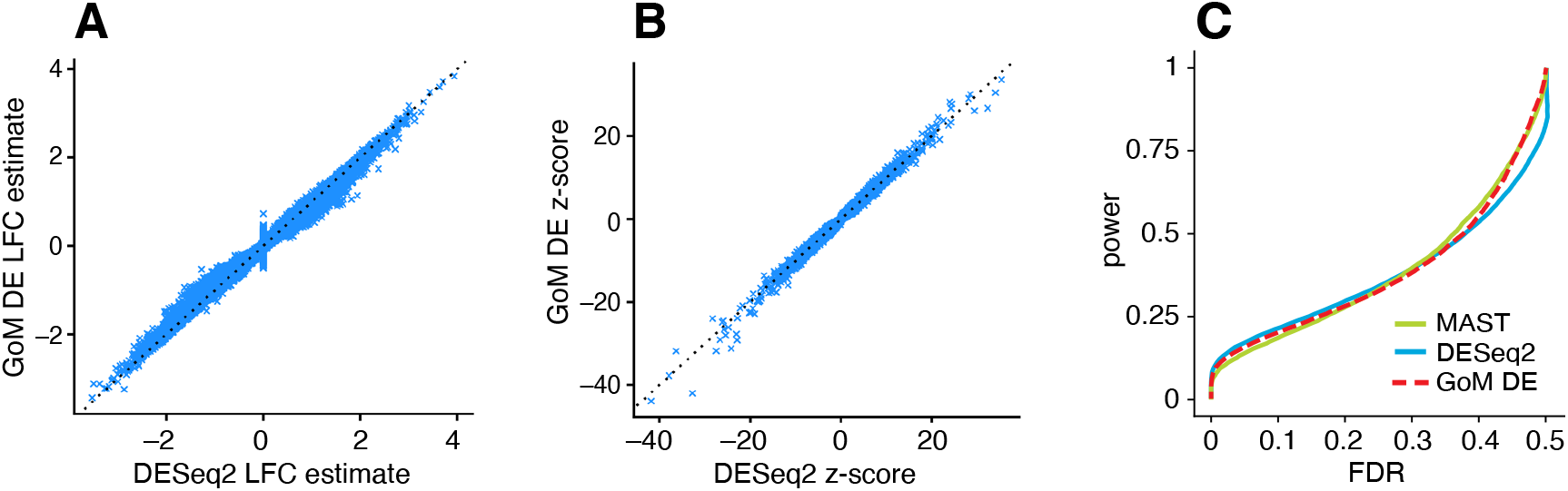
Evaluation of DE analysis methods in single-cell expression data sets in which cells were simulated from two groups without partial membership to these groups. Panels A and B compare posterior mean LFC estimates and posterior *z* -scores returned by performance in identifying differentially expressed genes in all simulated data sets; it plots power and false discovery rates (FDR) for the three methods compared as the *p*-value (MAST [84]), *s*-value (DESeq2) or *lfsr* threshold (GoM DE) is varied from 0 to 1. Power and FDR are calculated from the number of true positives (TP), false positives (FP), true negatives (TN) and false negatives (FN) as FDR = FP/(TP + FP) and power = TP/(TP + FN). See also Additional file 1: Figures S2, S3.

DESeq2 and the GoM DE analysis have several features in common: both are based on a Poisson model, and both use adaptive shrinkage [81, 83] to improve accuracy of the LFC estimates and test statistics. Therefore, we expected the GoM DE results to closely resemble DESeq2 in these simulations. Indeed, both methods produced nearly identical posterior mean LFC estimates, posterior *z*-scores (Figure 5A, B) and *s-*values (Additional file 1: Figure S3), and achieved very similar performance (Figure 5C). Although DESeq2 additionally estimates an overdispersion level for each gene, in these simulations DESeq2 correctly determined that the level of overdispersion was small for genes with large expression differences, which explains the strong similarity of the LFC estimates and posterior *z*-scores. MAST, owing to an approach that is very different from DESeq2 and the GoM DE analysis, yielded estimates that were less similar (Additional file 1, Figure S3), yet achieved comparable performance (Figure 5C).

Next we evaluated the GoM DE analysis methods in data sets in which the cells had varying degrees of membership to multiple topics. Since existing DE methods cannot handle the situation in which there are partial memberships to groups, we mainly sought to verify that the method behaves as expected in the ideal setting when data sets are simulated from the topic model (2). To provide some baseline for comparison, we also applied the method of Dey et al [47], which is not strictly a DE analysis method, but does provide a ranking of genes by their “distinctiveness” in each topic. This ranking is based on a simple KullbackLeibler (K-L) divergence measure; large K-L divergences should signal large differences in expression, as well as high overall levels of expression, so large K-L divergences should correspond to small DE *p*-values. Since the K-L divergence is not a signed measure, we omitted tests for negative expression differences from the evaluations, which was roughly half of the total number of possible tests for differential expression.

We performed 20 simulations with *K* = 2 topics and *n* = 200 cells, and another 20 simulations with *K* = 6 topics and *n* = 1,000 cells. To simplify evaluation, all genes either had the same rate of expression in all topics, or the rate was different in exactly one topic. As a result, the total number of expression differences in each data set was roughly the same regardless of the number of simulated topics. Other aspects of the simulations were kept the same as the first set of simulations (see Methods). Similar to before, we took the membership proportions to be known so that misestimation of the membership proportions would not be source of error in the GoM DE analysis and in calculation of the K-L divergence scores.

The largest K-L divergence scores in the simulated data sets reliably recovered true expression differences (Figure 6A, E). Therefore, the K-L divergence scores achieved good *true positives rates* (*i*.*e*., good power) at low *false positive rates*, FPR = FP*/*(TN + FP) (see Figure 5 for notation). However, for DE analysis a more relevant performance measure is the *false discovery rate*, FDR = FP*/*(TP + FP). Because the K-L divergence score does not fully account for uncertainty in the unknown gene expression differences, many genes with no expression differences among topics were also highly ranked, leading to poor FDR control (Figure 6D, H). By contrast, the GoM DE analysis better accounted for uncertainty in the unknown expression levels. The GoM DE analysis also more accurately recovered true expression differences at small *p-*values or *s-*values (Figure 6B, C, F, G), and therefore obtained much lower false discovery rates at corresponding levels of power (Figure 6D, H). Comparing the GoM DE analysis with and without adaptive shrinkage, the adaptive shrinkage did not necessarily lead to better performance (Figure 6D, H), but did provide more directly interpretable measures of significance (*s-*values or local false sign rates) by shrinking the LFC estimates, and adapting the rate of shrinkage to the data; for example, the expression differences were shrunk more strongly in the *K* = 6 data sets, correctly reflecting the much smaller proportion of true expression differences (compare Figure 6C and G).

**Figure 6.**
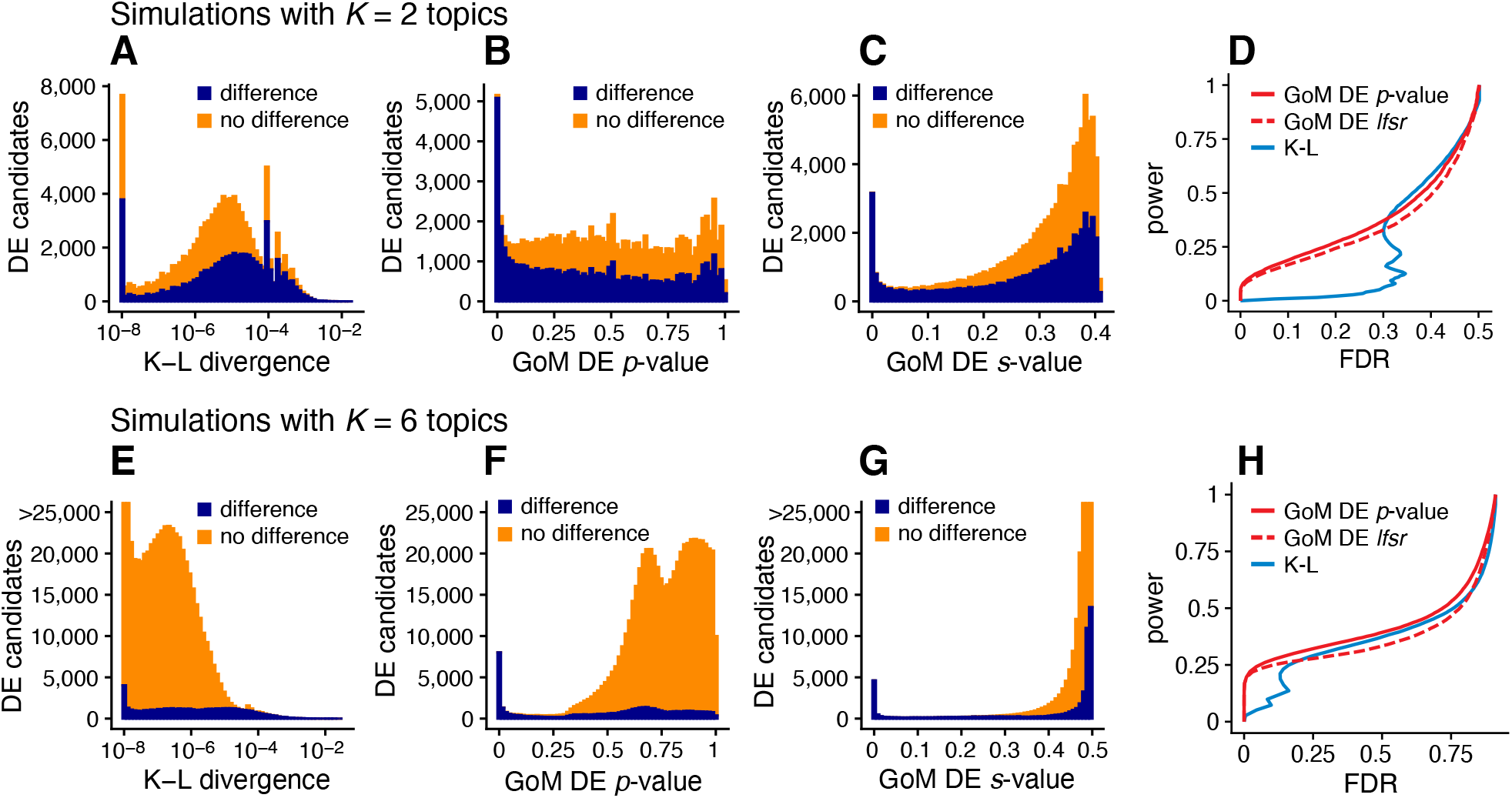
Evaluation of methods for identifying expression differences in single-cell expression data sets in which cells were simulated with partial membership to 2 topics (A–D) or 6 topics (E–H). Methods compared are the Kullback-Leibler (K-L) divergence score of [47] and GoM DE with adaptive shrinkage (*s-*values, *lfsr* ) and without adaptive shrinkage (*p*-values). The left-most panels (A, E) show the distribution of K-L divergence scores for all candidate expression differences (approximately half of 10,000 genes *×* 2 or 6 topics *×* 20 simulated data sets), shown separately for true expression differences (dark blue) and non-differences (orange). K-L divergence scores smaller than 10^*−*8^ are plotted as 10^*−*8^. Similarly, Panels B, C, F, G show the distribution of GoM DE *p-*values or *s-*values with or without adaptive shrinkage, separately among differences and non-differences. Panels D, H summarize performance in identifying expression differences; it shows power and FDR as the GoM DE *p-*value or *lfsr* are varied from 0 to 1, or as the K-L divergence score is varied from large to small. Note that in E and G some bar heights are actually larger than 25,000 but are cut off at 25,000 for better visualization.

Case study: scRNA-seq epithelial airway data from Montoro et al, 2018

We reanalyzed scRNA-seq data for *n* = 7,193 single cells sampled from the tracheal epithelium in wild-type mice [86]. The original analysis [86] used a combination of methods, including *t*-SNE, community detection [87], diffusion maps [88], and partitioning around medoids (PAM) to identify 7 epithelial cell types: abundant basal and secretory (club) cells; rare, specialized epithelial cell types, including ciliated, neuroendocrine and tuft cells; a novel subpopulation of “ionocytes”; and a novel basal-to-club transitional cell type, “hillock” cells. Although not large in comparison to other modern single-cell data sets, this data set is challenging to analyze, with complex structure, and a mixture of abundant and rare cell types. In contrast to the PBMC data set, there are no existing cell annotations to interpret the topics, so we must rely on inferences made from the expression data alone to make sense of the results.

The topic model fit to the UMI counts with *K* = 7 topics is shown in Figure 7A, and the results of the GoM DE analysis and subsequent GSEA are summarized in Figure 7. Although we do not have cell labels to compare with, distinctive genes emerging from the GoM DE analysis help connect some of the topics to known cell types. For example, the most abundant topics correspond well with predominant epithelial cell types in the lung: topic 1 shows strong overexpression of basal cell marker gene *Krt5* [89] (posterior mean l.e. LFC = 4.62); and distinctive genes in topics 2 and 3 include key secretory genes in club cells such as *Bpifa1*/*Splunc1* [90] (l.e. LFC = 4.93) and *Scgb1a1* [91] (l.e. LFC = 5.90).

**Figure 7.**
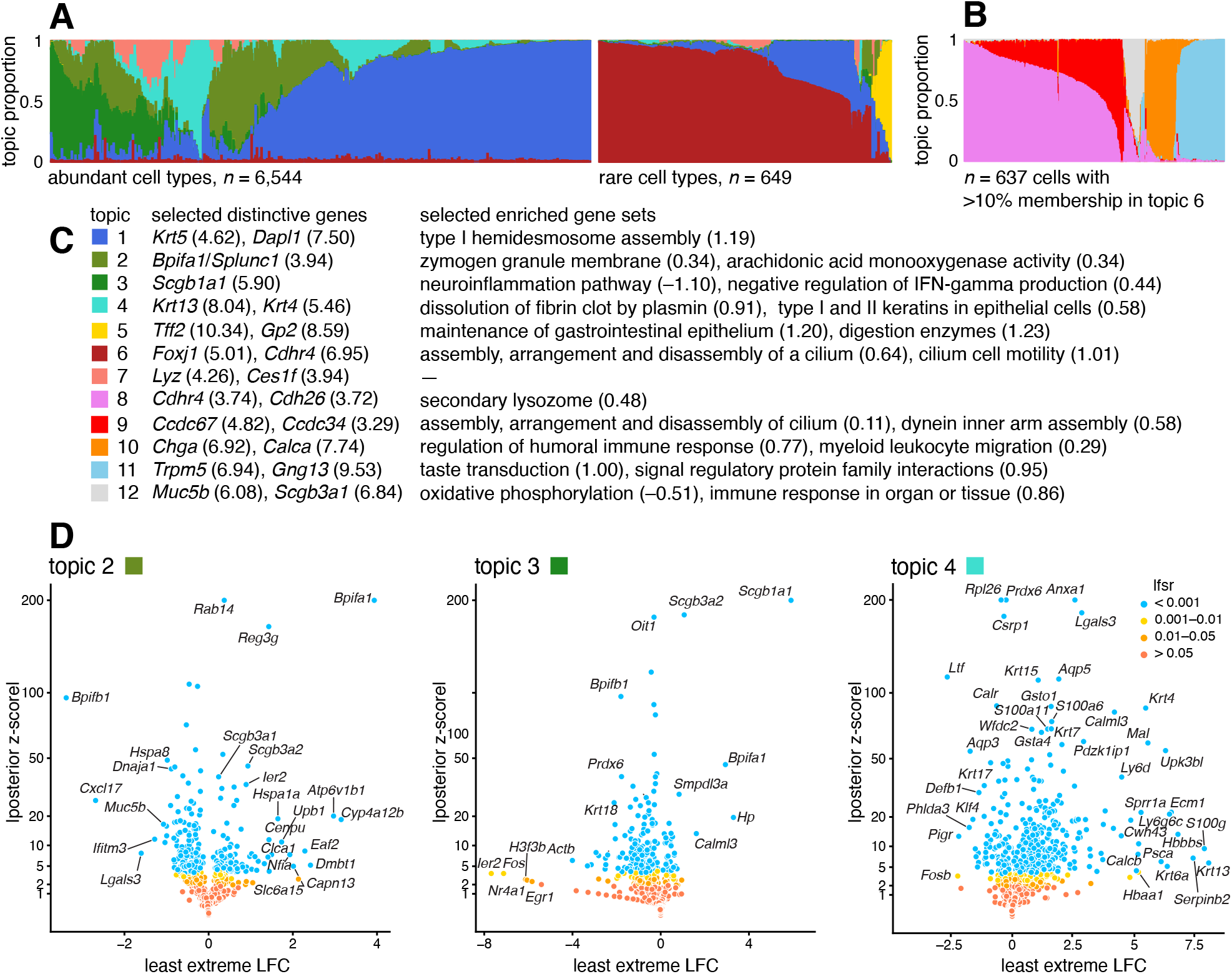
Structure in mouse epithelial airway data (*n* = 7,193 cells [86]) inferred from topic modeling (A, B), and GoM DE analysis (D) of selected topics using the membership proportions matrix **L** shown in A. In C, the topics are annotated by selected distinctive genes (numbers in parentheses are posterior mean l.e. LFCs) and selected enriched gene sets (numbers in parentheses are posterior mean estimates of the enrichment coefficients). In A, to better visualize the rare cell types, the cells were divided into two groups, “abundant” and “rare”, based on the estimated membership proportions, then the “abundant” cells were subsampled. The Structure plot in B was obtained by fitting another topic model, with *K* = 5 topics, to rare epithelial cell types (defined as the subset of 637 cells *i* with at least 10% membership to topic 6). The volcano plots show posterior estimates of l.e. LFC vs. posterior *z* -scores for 18,388 genes. A small number of genes with extreme posterior *z* -scores are shown with smaller posterior *z* -scores so that they fit within the y-axis range. See also the interactive volcano plots (Additional file 5: S6), GoM DE results (Additional file 2: Table S2, Additional file 1: Figures S6; S7), and GSEA results (Additional file 3: Tables S5, S6).

The “hillock” transitional cells, which were originally identified via a diffusion maps analysis [86], emerge as a single topic (topic 4, cyan), with *Krt13* (l.e. LFC = 8.04) and *Krt4* (l.e. LFC = 5.46) being among the most distinctive genes. The transitional nature of these cells is evoked by their mixed membership; only 237 out of the 7,193 cells have *>*90% membership to this topic.

Other less abundant epithelial cell types emerge as separate topics once a topic model is fit separately to the subpopulation of these rare cell types (Figure 7B). These topics recover ciliated cells (topics 8, 9; *Ccdc153*, posterior l.e. LFC = 5.39), neuroendocrine cells (topic 10; *Chga*, l.e. LFC = 6.92) and tuft cells (topic 11; *Trpm5*, l.e. LFC = 6.94). Note that *Foxi1+* ionocytes were previously identified as a novel cell type from a small cluster of 26 cells [86], but our analysis failed to distinguish this very rare cell type from the neuroendocrine cells (Additional file 1: Figures S4, S5).

The topics also capture biologically relevant *continuous substructure* in club cells (topics 2 and 3) and ciliated cells (topics 8 and 9) that was not discovered in the original analysis [86]. This continuous substructure may be reflective of finer scale cell differentiation or specialization of function. In particular, we interpret topic 3 as capturing “canonical” or “mature” (*Scgb1a1+*, l.e. LFC = 5.90) club cells [90], with negative regulation of inflammation, whereas cells with greater membership to topic 2 are “club-like” (*Bpifa1/Splunc1+*, l.e. LFC = 3.94) [89, 91]. Topic 9, similarly, appears to represent “canonical” ciliated cells, featuring upregulated genes such as such as *Ccdc67/Deup1* (l.e. LFC = 4.82) and *Ccdc34* (3.29) [89, 92, 93], and enrichment of Gene Ontology terms [94] such as cilium organization (GO:0044782) and axonemal dynein inner arm assembly (GO:0036159).

In summary, by taking a topic-model-based approach we identified and annotated well-characterized cell types such as basal cells, as well less distinct but potentially interesting substructures such as “Hillock” cells and club cell subtypes.

Case study: Mouse sci-ATAC-seq Atlas data from Cusanovich et al, 2018

We reanalyzed data from the Mouse sci-ATAC-seq Atlas [97], comprising 81,173 single cells in 13 tissues. First, to provide an overview of the primary structure in the whole data set, we fit a topic model with *K* = 13 topics to these data. The topics correspond closely to the clusters identified in [97] (Additional file 1: Figure S8), and several different tissues are distinguished by different topics (Figure 8A). For the 4 tissues that have replicates, the replicates show a similar composition of the topics (Figure 8A).

**Figure 8.**
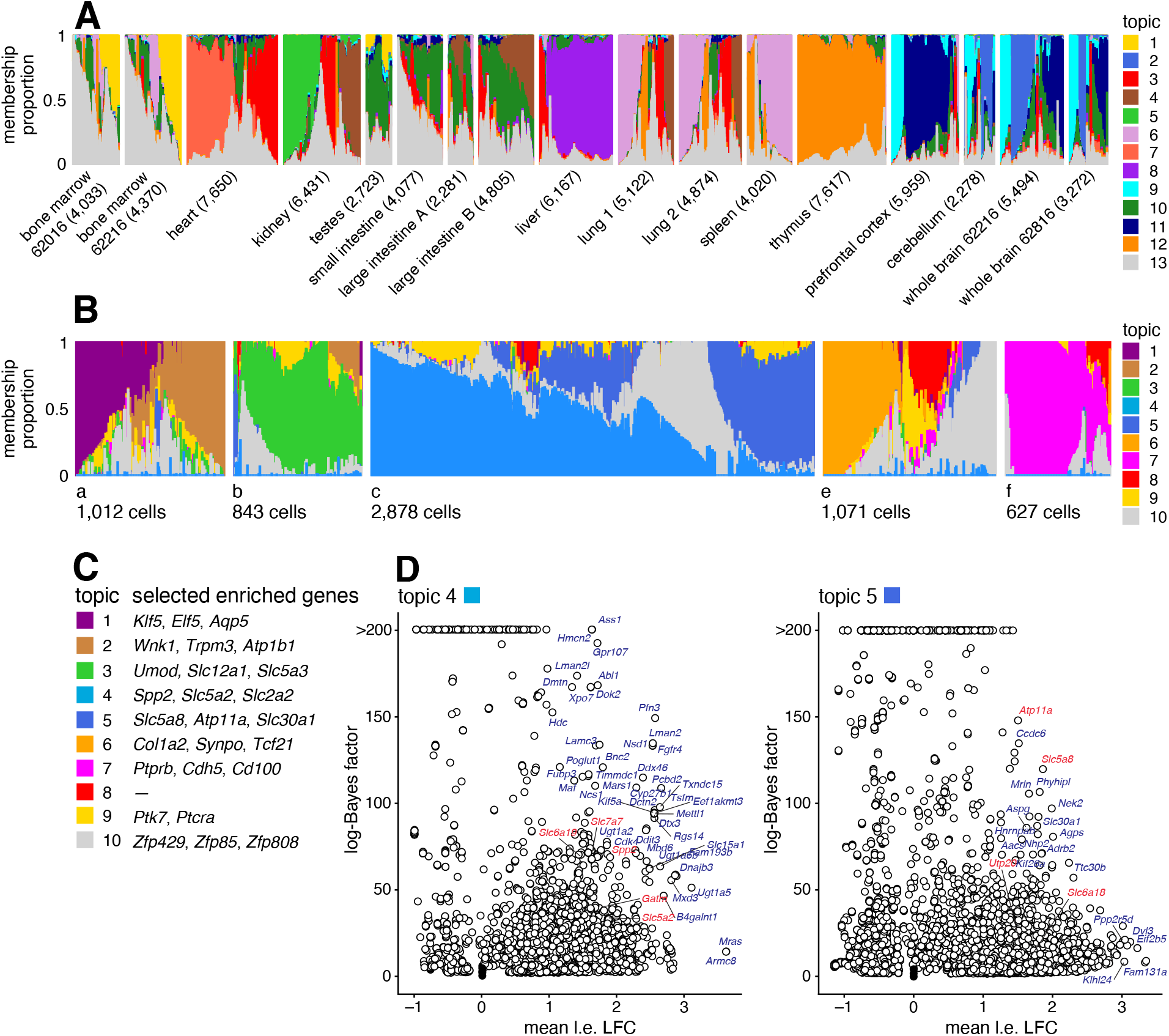
(A) Structure in Mouse Atlas sci-ATAC-seq data (*n* = 81,173) inferred from topic modeling, with *K* = 13 topics; (B) topic model fit to kidney cells (*n* = 6,431) with *K* = 10 topics; (C, D) gene-based enrichment analysis of differentially accessible peaks for the kidney cell topics shown in B, in which peaks are linked to genes using Cicero [95]. In A, the cells are grouped by tissue, and replicates (for bone marrow, large intestine, lung and whole brain) are shown as separate tissues. Numbers in parentheses next to each tissue give the number of cells in that tissue. In D, marker genes for S1 (topic 4) and S3 (topic 5) proximal epithelial tubular cells are highlighted in red (see Table 1 of [96]). “Mean l.e. LFC” is the average l.e. LFC among all peaks connected to the gene, restricted to l.e. LFCs with *lfsr* 0.05. Log-Bayes factors greater than 200 are shown as 200 in the volcano plots. See Additional file 1: Figure S9 and Additional file 6: Table S7 for more gene enrichment results. In B, the cells are subdivided into 5 groups (a–f) only to improve visualization. See also Additional file 1: Figure S10 which compares the topics in B to cell-type predictions based on clustering [97].

Next we performed a more detailed analysis of just the kidney (6,431 cells), fitting a topic model with *K* = 10 to just these cells. We focussed on the kidney cells because, as noted previously [97, 98], both expression and chromatin accessibility vary in relation to the spatial organization of the renal tubular cells, and we predicted that this spatial structure could be better captured by topics rather than by traditional clustering methods. To interpret these topics obtained from chromatin accessibility data, we first used the GoM DE analysis to identify differentially accessible peaks for each topic, then we used “co-accessibility” as predicted by Cicero [95, 97] to connect genes to peaks representing distal regulated sites. Finally, we performed a simple enrichment analysis to identify the “distinctive genes” for each topic, which we defined as the genes with many distal regulatory sites that were differentially accessible.

**Table 1.** GoM DE simulation running times for the single-cell data sets with *n*_*s*_ = 10, 000 simulation states; *n* is the number of cells, *m* is the number of genes or accessibility peaks analyzed, and *K* is the number of topics. See “Computing environment” for more details.

| data set | $n$ | $m$ | $K$ | runtime |
| --- | --- | --- | --- | --- |
| PBMC [29] | 94,655 | 21,952 | 6 | 5.1 h |
| Epithelial airway [86] | 7,193 | 18,388 | 7 | 1.0 h |
| Mouse Atlas, kidney only [97] | 6,431 | 270,864 | 10 | 3.2 h |
| Hematopoietic system [104] | 2,034 | 126,719 | 10 | 1.1 h |

The results of these analyses are shown in Figure 8. Many of the distinctive genes (Figure 8, Additional file 1: Figure S9, Additional file 6: Table S7) clearly relate topics to known kidney cell types. For example, topic 1 is enriched for genes *Klf5* and *Elf5* which relate to the collecting duct [98, 99]; topic 3 is enriched for genes *Umod, Slc12a1* associated with the loop of Henle [98, 100]); and topics 2, 6, 7 are respectively enriched for genes related to the distal convoluted tubule (*Wnk1*), podocytes (*Col1a2*) and glomerular endothelial cells (*Ptprb*).

Most interestingly, spatial organization of the proximal tubule is captured by two topics; topic 4 is enriched for *Slc5a2* (also known as *Sglt2*) and *Slc2a2* (also known as *Glut2*), associated with the S1 segment of the proximal tube [96, 101, 102], and topic 5 is enriched for *Slc5a8* (*Smct1*) and *Atp11a*, related to the S3 segment [96, 103]. This result illustrates the ability of the topic model to capture continuous variation in membership of two somewhat complementary processes, which traditional clustering methods are not designed for.

### Case study: chromatin accessibility profiles of the hematopoietic system from Buenrostro et al, 2018

Buenrostro et al [104] studied 2,034 single-cell ATACseq profiles of 10 cell populations isolated by FACS to characterize regulation of the human hematopoietic system. Both PCA and *t*-SNE showed, visually, the expected structure into the main developmental branches (Figure 2 in [104]). However, neither PCA nor *t*-SNE isolated these branches as *individual dimensions* of the embedding. Identifying these branches may allow for more precise characterization of the underlying regulatory patterns. Here, by fitting a topic model to the data, the main developmental branches are identified as individual topics (Figure 9A): topic 3, pDC; topic 4, erythroid (MEP); topic 5, lymphoid (CLP); and topic 6, myeloid (GMP and monocytes). Another topic captures the cells at the top of the developmental path (topic 1; HSC and MPP). Other cells at intermediate points in the developmental trajectory, such as CMP, GMP and LMPP cells, are more heterogeneous, and this is reflected by their high variation in topic membership.

**Figure 9.**
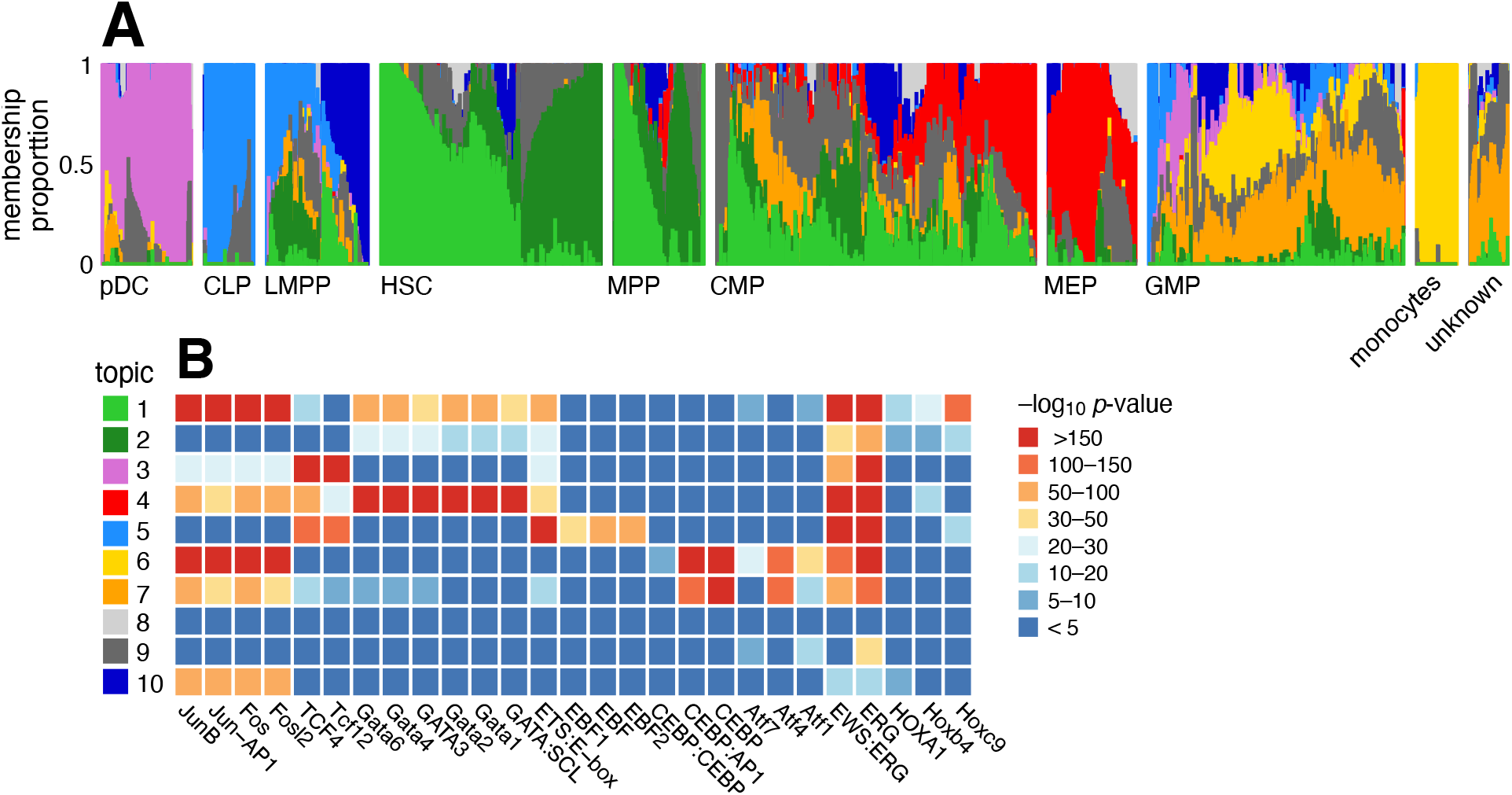
Structure in human hematopoietic system data [104] (*n* = 2,034 cells) inferred from the topic model with *K* = 10 topics (A), and HOMER motif enrichment analysis [105] applied to the results of the GoM DE analysis (B). In the Structure plot, the cells are grouped by FACS, as well as an unknown population from human bone marrow [104]. Panel B shows HOMER enrichment results for selected motifs (for the full results, see Additional file 7: Table S8). Acronyms used: common lymphoid progenitor (CLP); common myeloid progenitor (CMP); granulocyte-macrophage progenitor (GMP); hematopoietic stem cell (HSC); lymphoid-primed multipotent progenitor (LMPP); megakaryocytic-erythroid progenitor (MEP); multi-potent progenitor (MPP); plasmacytoid dendritic cells (pDC).

To better interpret the regulatory patterns behind each topic, we identified transcription factor (TF) motifs that were enriched for differentially accessible regions in each topic (Figure 9B, Additional file 7: Table S8). Many of the top TF motifs (as ranked by HOMER *p*-values [105]) point toward regulation of the main developmental trajectories, such as EBF motifs in topic 5 (lymphoid), CEBP motifs in topics 6 and 7 (myeloid), and Hox motifs in topic 1 (HSC and MPP cells). A few topics (topics 8–10) are much less abundant and do not align well with the FACS cell types, and their motif enrichment results were correspondingly more difficult to interpret.

A complication that arose in analyzing these data, which was also noted in [104], is that the cells were obtained from different sources, and this shows up as systematic variation in the chromatin accessibility. This donor effect is captured by topics 1 and 2 in HSC and MPP cells, and, to a lesser extent, in CMP and LMPP cells (Additional file 1: Figure S11). Topic 1 is enriched for Jun and Fos TF motifs, similar to what was found in [104].

## Discussion

The GoM DE analysis is part of a *topic-model-based pipeline* for analysis of single-cell RNA-seq [47] or ATAC-seq data [49]. This pipeline includes the following steps: (1) fit a topic model to the data; (2) visualize the structure inferred by the topic model; (3) run the GoM DE analysis with the estimated topics; and, optionally, (4) perform other downstream analyses using the results of the GoM DE analysis, e.g., gene set enrichment analysis (for RNA-seq data) or motif enrichment analysis (for ATAC-seq data). Unlike most analysis pipelines for clustering and dimensionality reduction (e.g., [4, 19, 23, 26, 27]), the topicmodel-based pipeline is directly applied to the “raw” count data, and therefore does not require an initial step to transform and normalize the data which can lead to downstream issues in the statistical analysis [8, 106–108]. We presented several case studies illustrating the use of the topic-model-based pipeline to analyze single-cell RNA-seq and ATAC-seq data sets. From these case studies, we have drawn a few lessons on the practical challenges that may arise in applying topic modeling approaches to single-cell data, and we share these lessons here. (See also [47, 49] for related discussion.)

One practical question is how to choose *K*, the number of topics. Many papers have suggested different criteria for determining *K*. Our view, following [47], is that there is no single “best” *K*, and we recognize the advantages of learning topics at multiple settings of *K*; in some data sets, different *K*’s can reveal structure at different levels of granularity (for example, increasing the number of topics in the Mouse sci-ATACseq Atlas data revealed more structure within tissues; see https://tinyurl.com/2p99swdk). We have found that it is often helpful to start with a smaller *K* to elucidate the less granular structure, which is often easier to interpret, then rerun the topic modeling with larger *K* to identify finer structure.

We proposed annotating topics by distinctive genes identified using the l.e. LFC. One drawback is that this does not reveal the commonalities that may exist among multiple topics, for example, topics corresponding to subpopulations within a common class of cells. A simple alternative to the l.e. LFC, which is also implemented in the `fastTopics` R package, is to compare against expression under the “null model” (see Methods). We view this as a complementary LFC metric that may reveal additional insights into the topics.

Donor, batch or other technical effects in the singlecell RNA-seq or ATAC-seq data can complicate the analysis and interpretation of the topics if these effects are not small. Since these effects are usually not known, usually we must assess their impact indirectly [109]. For example, the Mouse sci-ATAC-seq Atlas data included several replicates, but the replicate effects appeared to be small judging by the fact that the replicates showed a similar composition of topics. By contrast, the donor effects in the human hematopoietic system data were much larger, and in the topic model these donor effects were at least partially captured by individual topics. The broader question of how to deal with non-ignorable donor or batch effects—in particular, how to separate technical effects from biological effects of interest—remains a question of considerable debate and continued investigation [25, 39, 109–116]. In particular, it has been noted that attempting to “correct” for effects can sometimes remove differences that we would like to learn about such as differences in cell type proportions among the batches.

For modeling UMI counts, an open question is whether the Poisson or multinomial model (1) is sufficient, or whether more flexible models are needed. (This question was investigated in [69] for single-gene models, but not for multi-gene models.) Alternative models such as the negative binomial [117] or Poisson log-normal [80, 118], which can capture additional random variation (“overdispersion”) in underlying expression or measurement error, may result in more robust estimation of the topics.

In single-cell ATAC-seq data, the GoM DE analysis identifies differentially accessible peaks or regions. Usually these peak-level results need to be translated into biological units that are more useful for annotating the topics (e.g., genes, gene sets, transcription factors). In the analysis of the hematopoietic system single-cell ATAC-seq data, we used HOMER [105] to identify TF motifs enriched for differentially accessible peaks. In the analysis of the Mouse sci-ATAC-seq Atlas data, we identified genes enriched for differentially accessible distal regularity sites. Clearly, the quality of the gene enrichment results will depend on our ability to accurately associate peaks with genes. For this, we used the scores computed in [97] using Cicero [95]. However, there are now several alternatives to Cicero that may be preferred [19, 27, 28, 119–122], and in principle any of these approaches could be combined with the peak-level GoM DE results to identify relevant genes.

Recently developed technologies profile both transcription and chromatin accessibility in single cells [123, 124]. For such data, one could fit two topic models, one to the RNA-seq data and another to the ATAC-seq data. With a careful initialization of the topic model fitting algorithm, the topics may be more consistent across the two modalities. But it would be preferrable to analyze the multimodal data jointly for improved accuracy [125–130]. Potentially, the strategy used in MOFA [131, 132] could be adapted for topic modeling—that is, the transcripts and accessibility profiles would share the same membership proportions, **L**, but each modality would have a different **F**. However, it remains to be seen how well this strategy works in practice.

## Conclusions

To summarize, we have described a new method that aids in annotating and interpretating the “parts” of cells learned by fitting a topic model to scRNA-seq data or single-cell ATAC-seq data. Our method, GoM DE (differential expression analysis allowing for grades of membership), can be viewed as an extension of existing differential expression methods that allows for mixed membership to multiple groups or topics.

## Methods

### Models for single-cell ATAC-seq data

In single-cell ATAC-seq data, *x*_*ij*_ is the number of unique reads mapping to peak or region *j* in cell *i*. Although *x*_*ij*_ can take non-negative integer values, it is common to “binarize” the accessibility data [e.g., 19, 74, 133–135], meaning that *x*_*ij*_ = 1 when at least one read in cell *i* maps to region *j* and *x*_*ij*_ = 0 otherwise. For this reason, one might prefer to model the binarized accessibility values as binomial (Bernoulli) random variables. A multinomial model, on the other hand, should better capture the sampling process for reads mapping to regions, but does not account for the truncation of read counts above 1. Therefore, we view both the binomial and multinomial models as approximations. As we explain next, under reasonable assumptions the binomial and multinomial models are similar to each other so it may not matter which model one chooses.

The multinomial topic model for analyzing single-cell ATAC-seq data was suggested by [49]. They assumed the multinomial model (1) in which the *x*_*ij*_’s are binarized accessibility values instead of UMI counts.

A binomial model was proposed in [74],

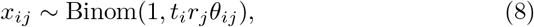

where *t*_*i*_ *>* 0 is a cell-specific factor that depends on sequencing coverage and other properties (e.g., amplification, read post-processing [136]), *r*_*j*_ *>* 0 is a regionspecific factor (say, proportional to the size of the region), and the *θ*_*ij*_’s capture additional variation in accessibility across cells and regions. Moving forward, we make the simplifying assumption that the regions are all approximately the same size; that is, *r*_*j*_ = 1 for all *j* = 1, …, *m*.

The binomial model (8) is closely related to a multinomial model. To make the connection, we first note that the binomial model with *r*_*j*_ = 1 for all *j* can be approximated by a Poisson model,

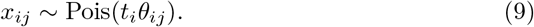

This will be a good approximation when the *θ*_*ij*_’s are small and the cell-specific factors *t*_*i*_ are large, which is usually the case in single-cell ATAC-seq data. Next, we note that the Poisson model (9) and multinomial model (1) are closely related if we choose the size factors to be *t*_*i*_ = *s*_*i*_ [69, 137]; this implies **Θ** *≈* **Π**, where **Θ** is the *n× m* matrix with entries *θ*_*ij*_. By these arguments, the binomial model (8) (also the model used in [74]) and the multinomial model (1) (also the model used in [49]) are similar, and connecting the two models clarifies the assumptions made by each of the models. In particular, the multinomial topic model (1–2) used here and in [49] assumes a low-rank structure in the *θ*_*ij*_’s across cells and regions; *i*.*e*., **Θ** *≈* **LF**^*T*^ .

### Differential expression analysis allowing for grades of membership

#### Derivation of GoM DE model

In the “Methods overview,” we motivated the GoM DE model (3) as extending a basic Poisson model expression to allow for partial membership to *K* groups or topics. The GoM DE model can also be motivated from an approximation to the topic model. Recall, the topic model is a multinomial model (1) in which the multinomial probabilities *π*_*ij*_ are given by affine combinations of the expression levels *f*_*jk*_ in the *K* topics, 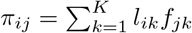. The non-negativity constraints *l*_*ik*_ *≥* 0, *f*_*jk*_ *≥* 0 and sum-to-one constraints 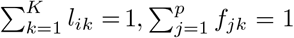 ensure that the *π*_*ij*_’s are multinomial probabilities. From a basic identity relating the multinomial and Poisson distributions [138, 139], the multinomial likelihood for the topic model can be replaced with a likelihood formed by a simple product of independent Poissons; that is,

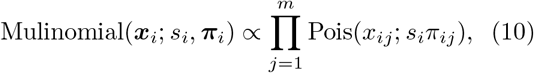

where ***x***_*i*_ = (*x*_*i*1_, …, *x*_*im*_) and ***π***_*i*_ = (*π*_*i*1_, …, *π*_*im*_). The approximation then comes from no longer requiring the *π*_*ij*_’s to be multinomial probabilities by removing the constraint that *f*_1*k*_ + *· · ·* + *f*_*mk*_ = 1. This allows us to analyze the genes *j* = 1, …, *m* independently. This is a good approximation so long as *s*_*i*_ is large and the *f*_*jk*_’s are small. (A similar approximation was used for GLM-PCA [8].) To be explicit about this approximation, we say *π*_*ij*_ *≈ θ*_*ij*_ (which are no longer guaranteed to be multinomial probabilities) and *f*_*jk*_ *≈ p*_*jk*_ (which are no longer guaranteed to sum to one), resulting in the GoM DE model, which for convenience we restate here:

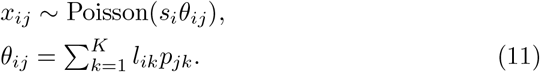

*“Null” model*

The simplest Poisson model of the form (3) is one in which *θ*_*ij*_ is the same across all cells *i*; that is, *θ*_*ij*_ = *p*_*j*0_ for all *i* = 1, …, *n*. We treat this a “null” model, which can be used to make certain comparisons, e.g., to estimate changes in expression in relative to expression in all cells. The maximum-likelihood estimate (MLE) of *p*_*j*0_ under the null model is

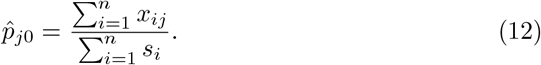

### Estimation of log-fold change

In practice, we have found the l.e. LFC to work well, so in our results we use the l.e. LFC. But the l.e. LFC may not be appropriate in all circumstances, and for this reason we note that the GoM DE analysis framework is quite general and accommodates alternatives to the l.e. LFC. Two alternatives are implemented in the software. One alternative is to compare with the “null” model,

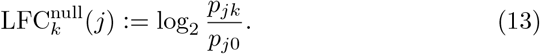

Another treats one topic *l* as a “reference topic”, and compares all other topics *k /*= *l* to *l* using (4).

#### Maximum-likelihood estimation

A convenience of the Poisson model allowing for grades of membership is that we can reuse Poisson NMF computations (described below and in more detail in [48]) to compute MLEs of the unknowns *p*_*jk*_: if we consider all genes *j* = 1, …, *m* simultaneously, we recover a Poisson NMF model, *x*_*ij*_ *∼* Poisson(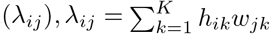, by setting *h*_*ik*_ = *s*_*i*_*l*_*ik*_, *w*_*jk*_ = *p*_*jk*_. Therefore, we can reuse the Poisson NMF algorithms to compute MLEs of the unknowns *p*_*jk*_.

#### *Maximum* a posteriori *estimation*

To improve numerical stability in the parameter estimation, we compute *maximum a posteriori* (MAP) estimates of *p*_*j*1_, …, *p*_*jK*_ in which each *p*_*jk*_ is assigned a gamma prior, *p*_*jk*_ *∼* Gamma(*α, β*), with *α* = 1 + *ε, β* = 1, and *ε >* 0. Typically *ε* will be some small, positive number, e.g., *ε* = 0.1. Here we use the parameterization of the gamma distribution from [140] in which *α* is the shape parameter and *β* is the inverse scale parameter; under this parameterization, the mean is *α/β* and the variance is *α/β*^2^. The maximum-likelihood computations can be reused for MAP estimation with this gamma prior by adding “pseudocounts” to the data; specifically, MAP estimation of *p*_*j*1_, …, *p*_*jK*_ given counts *x*_1*j*_, …, *x*_*nj*_ and membership proportions **L** and is equivalent to maximum-likelihood estimation of *p*_*j*1_, …, *p*_*jK*_ given counts *x*_1*j*_, …, *x*_*nj*_, *ε*, …, *ε* and membership proportions matrix 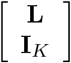, where **I**_*K*_ is the *K × K* identity matrix. Unless otherwise stated, we added *ε* = 0.1 pseudocounts to the data.

#### Quantifying uncertainty and stabilizing LFC estimates

We implemented a simple Markov chain Monte Carlo (MCMC) algorithm [141, 142] to quantify uncertainty in the LFC estimates. Although normal approximations to likelihoods are typically used by DE methods to quickly obtain analytical measures of uncertainty (e.g., standard errors, confidence intervals) for LFCs, we found that normal approximations to the likelihoods from (5) were sometimes poorly behaved, particularly for lowly expressed genes. Another consideration was that the analytical solutions provide confidence intervals for the unknowns *p*_*jk*_, but ultimately we are interested in quantifying uncertainty in the l.e. LFCs (6) which do not have a simple linear relationship to the *p*_*jk*_’s. Therefore, it is unclear whether the standard analytical solutions can be applied to the l.e. LFCs without making further approximations or simplifications.

MCMC is typically computationally intensive, but with careful implementation (e.g., use of sparse matrix operations and multithreaded computations) the MCMC algorithm is quite fast. Other benefits of using MCMC is that the algorithm can straightforwardly accommodate different choices of LFC statistics and no normality assumptions are needed.

The basic idea behind the MCMC algorithm is as follows: for a given gene *j*, simulate the posterior distribution of the LFC statistic by performing a “random walk” on ***g***_*j*_ = (*g*_*j*1_, …, *g*_*jK*_), where *g*_*jk*_ := log *p*_*jk*_, *k* = 1, …, *K*. The random walk generates a sequence of states 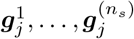, in which *n*_*s*_ denotes the pre-specified length of the simulated Markov chain. After choosing an initial state 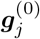, each new state 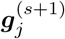 is generated from the current state 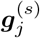 by the following procedure: first, a topic *k* 1, …, *K* is chosen uniformly at random; next, a proposed state 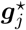 is generated as 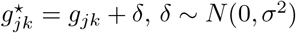, with 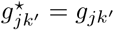 for all *k*^*l*^ */*= *k*. Assuming an (improper) uniform prior for the unknowns, Pr(*p*_*jk*_) ∝ 1, the proposed state is accepted into the Markov chain with probability

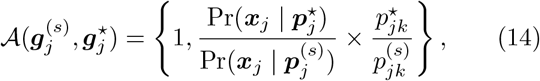

in which ***x***_*j*_ is the *j*th column of the counts matrix **X, *x***_*j*_ = (*x*_1*j*_, …, *x*_*nj*_il), and Pr(***x***_*j*_ | ***p***_*j*_) is the likelihood at 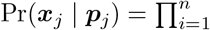 Poisson(*x*_*i*_; *s*_*i*_*θ*_*i*_). (Note that ***x***_*j*_ may include pseudocounts.) The standard deviation of the Gaussian proposal distribution, *σ*, is a tuning parameter. (Unless otherwise stated, we used *σ* = 0.3) The additional 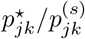 term in the acceptance probability is needed to account for the fact that we are simulating the log-transformed parameters ***g***_*j*_, not ***p***_*j*_ [143, p. 11]. When the proposal is not accepted, the new state is simply copied from the previous state,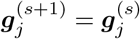

Most of the effort in running the MCMC goes into computing the acceptance probabilities (14), so we have carefully optimized these computations. For example, we have taken advantage of the fact that the count vectors ***x***_*j*_ are typically very sparse. Additionally, these computations can be performed in parallel since the Markov chains are simulated independently for each gene *j*.

Once Monte Carlo samples ***g***^(*s*)^, for *s* = 1, …, *n*_*s*_, have been simulated by this random-walk MCMC, we compute posterior mean LFC estimates, and quantify uncertainty in the LFC estimates. For example, expressing the l.e. LFC for gene *j* and topic *k* as a function of the unknowns, 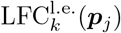, the posterior mean l.e. LFCs are calculated as 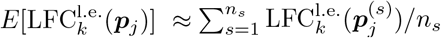.

The final step in the GoM DE analysis is to perform adaptive shrinkage [81] to stabilize the posterior mean estimates. To implement this step, we used the `ash` function from the `ashr` R package [144]. We used the same settings as DESeq2 to replicate as closely as possible the performance of DESeq2 with adaptive shrinkage. DESeq2 calls `ash` with `method = “shrink”`, which sets the prior to be a mixture of uniforms without a point-mass at zero.

The adaptive shrinkage method takes as input a collection of effect estimates 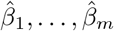 and associated standard errors ŝ_1_, …, ŝ_*m*_. In this setting, it is not immediately obvious what are the standard errors, in part because the posterior distribution of the unknowns is not always symmetric about the mean or median. To provide a reasonable substitute summarizing uncertainty in the estimates, we computed Monte Carlo estimates of highest posterior density (HPD) intervals. A (1 − *α*) HPD interval is the smallest interval that contains 100(1 − *α*)% of the probability mass [145, 146]. Specifically, let [*a*_*jk*_, *b*_*jk*_] denote the (1 − *α*)

HPD interval for the LFC estimate of gene *j* in topic *k*, and let 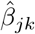denote the posterior mean. We defined the standard error as 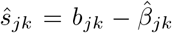 when 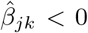 otherwise, 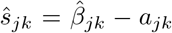. Defining the standard errors in this way prevented overshrinking of estimates that were uncertain but had little overlap with zero.

We set the size of the HPD intervals to 1 − *α* = 0.68 so that the ŝ_*jk*_ would recover conventional standard error calculations when the posterior distribubtion is well approximated by the normal distribution. The revised posterior means and standard errors returned by the adaptive shrinkage method were then used by `ashr` to calculate test statistics including posterior *z*-scores (defined as the posterior mean divided by the posterior standard error [147]), local false sign rates (*lfsr*) and *s*-values.

An important question is the choice of *n*_*s*_. One heuristic way to assess whether *n*_*s*_ is large enough is to perform two independent MCMC runs initialized with different pseudorandom number generator states (“seeds”) and check consistency of the posterior estimates from the two runs. (We checked consistency of the posterior estimates after stabilizing the estimates using adaptive shrinkage, as described above.) In simulated data sets (below), comparison of two independent MCMC runs suggested that *n*_*s*_ = 10,000 was sufficient to obtain reasonably accurate estimates of posterior means and posterior *z*-scores for all genes (Additional file 1: Figure S2). Therefore, we performed initial MCMC simulations for all single-cell data sets using *n*_*s*_ = 10,000. The runtimes for performing these MCMC simulations on the single-cell data sets (described below), with *n*_*s*_ = 10,000, are given in Table 1. Although this consistency check suggested that running a simulation with *n*_*s*_ = 10,000 would be “good enough”, to provide additional assurance we performed another consistency assessment in the PBMC data set. We found that even better consistency was achieved with *n*_*s*_ = 100,000 (Additional file 1: Figure S12). Therefore, to provide more reliable results, the final GoM DE results were generated with *n*_*s*_ = 100,000.

The GoM DE analysis methods are implemented in the `de_analysis` function in the `fastTopics` package [184].

### Single-cell data sets

All data sets analyzed were stored as sparse *n× m* matrices **X**, where *n* was the number of cells and *m* was the number of genes or regions. The data sets are summarized in Table 1.

#### Preparation of scRNA-seq data

Since the topic model is a multinomial model of count data, no log-normalization or other transformation of the scRNA-seq molecule counts was needed. Further, we kept all genes other than those with no variation in the data set. (This is done in part to demonstrate that our methods are robust to including genes with little variation.) Also note that due to the use of sparse matrix techniques in our software implementations, including genes with low variation did not greatly increase computational effort.

#### Preparation of single-cell ATAC-seq data

As previously suggested [19, 133–135]), we “binarized” the single-cell ATAC-seq data; that is, we assigned *x*_*ij*_ = 1 (“accessible”) when least one fragment in cell *i* mapped to peak or region *j*, otherwise *x*_*ij*_ = 0 (“inaccessible”). There are at least a couple reasons for doing this. For small peaks (say, *<* 5 kb), read counts do not provide a reliable quantitative measure of accessibility in single cells. This is because the (random) first insertion restricts the space for subsequent insertions. Additionally, insertions could occur within the same site on the same allele or on each of the two alleles, complicating interpretation of the read counts.

Like the RNA molecule count data (see above), we kept all regions except those that showed no variation.

#### PBMC data from Zheng et al, 2017

We combined reference transcriptome profiles generated from 10 bead-enriched subpopulations of PBMCs (Donor A) processed using Cell Ranger 1.1.0 [29, 148]. We downloaded the “Gene/cell matrix (filtered)” tar.gz file from the 10x Genomics website for each of the following 10 FACS-purified data sets: CD14+ monocytes, CD19+ B cells, CD34+ cells, CD4+ helper T Cells, CD4+/CD25+ regulatory T Cells, CD4+/CD45RA+/CD25naive T cells, CD4+/CD45RO+ memory T Cells, CD56+ natural killer cells, CD8+ cytotoxic T cells and CD8+/CD45RA+ naive cytotoxic T cells. After combining these 10 data sets, then filtering out unexpressed genes, the combined data set contained molecule counts for 94,655 cells and 21,952 genes; 97.1% of the molecule counts were zero.

In Figure 1, the 54,132 cells from these data sets were labeled as “T cells”: CD4+ helper T Cells, CD4+/CD25+ regulatory T Cells, CD4+/CD45RA+/CD25naive T cells, CD4+/CD45RO+ memory T Cells and CD8+/CD45RA+ naive cytotoxic T cells.

#### Epithelial airway data from Montoro et al, 2018

We analyzed a mouse epithelial airway data set from [86, 149]. These were gene expression profiles of trachea epithelial cells in C57BL/6 mice obtained using droplet-based 3’ scRNA-seq, processed using the GemCode Single Cell Platform. We downloaded file `GSE103354_Trachea_droplet_UMIcounts.txt.gz`. This file also contained the cluster assignments that we compared with. (In [86], the samples were subdivided into 7 clusters using a community detection algorithm.) After removing genes that were not expressed in any of the cells, the data set contained molecule counts for 7,193 cells and 18,388 genes (90.7% of counts were zero).

#### Mouse Atlas data from Cusanovich et al, 2018

Cusanovich et al [97] profiled chromatin accessibility by single-cell combinatorial indexing ATAC-seq (sciATAC-seq) [150, 151] in nuclei from 13 distinct tissues of a 8-week-old male C57BL/6J mouse. Replicates for 4 of the 13 tissues were obtained by profiling chromatin accessibility in a second mouse. We downloaded the (sparse) binarized peak × cell matrix in RDS format, `atac_matrix.binary.qc_filtered.rds`, from the Mouse sci-ATAC-seq Atlas website [152]. We also downloaded `cell_metadata.txt` which included cell types estimated by a clustering of the cells (see Table S1 in [97]). The full data set used in our analysis (13 tissues, including 4 replicated tissues) consisted of the binary accessibility values for 81,173 cells and 436,206 peaks (1.2% overall rate of accessibility). Note that all peaks had fragments mapping to at least 40 cells, so no extra step was taken to filter out peaks.

Separately, we analyzed the sci-ATAC-seq data from kidney only, in which peaks with fragments mapping to fewer than 20 kidney cells were removed, resulting in data set containing binary accessibility values for 6,431 cells and 270,864 peaks. Base-pair positions of the peaks were based on Mouse Genome Assembly mm9 (NCBI and Mouse Genome Sequencing Consortium, Build 37, July 2007).

From the Mouse sci-ATAC-seq website, we also downloaded the file `master_cicero_conns.rds` containing the Cicero co-accessibility predictions [95, 152], which we used to link chromatin accessibility peaks to genes. For the kidney data, we connected a peak given in the “Peak2” column of the Cicero co-accessibility data table to a gene given in the “peak1.tss.gene id” column if the “cluster” column was 11, 18, 22 or 25. (These four clusters were the main kidney-related clusters identified in [97].) This extracted, for each gene, the distal and proximal sites connected to the gene associated with Peak1 (specifically, a gene in which the transcription start site overlaps with Peak1). Among the 22,194 genes associated with at least one peak, the median number of peaks connected to a gene was 19, and the largest number of peaks was 179 (for *Bahcc1* on chromosome 11). Among the 270,864 peaks included in the topic modeling analysis, 113,489 (42%) were connected to at least one gene, 95% of peaks were connected to 10 genes or fewer, and the largest number of connected genes was 60.

#### Human hematopoietic system data from Buenrostro et al, 2018

Buenrostro et al [104] used FACS to isolate 10 hematopoietic cell populations from human bone marrow and blood, then the cells were assayed using single-cell ATAC-seq. The processed single-cell ATACseq data were downloaded from [153]; specifically, file `GSE96769_scATACseq_counts.txt.gz` containing the fragment counts, and file `GSE96769_PeakFile 20160207.bed.gz` containing peaks obtained from bulk ATAC-seq data [104]. Although there may be benefits to calling peaks using aggregated single-cell data instead [154], we used the original accessibility data based on the bulk ATAC-seq peaks so that our analysis was more directly comparable to the analysis of [104].

Following [104, 154], we extracted the 2,034 samples passing quality control filters, then we “binarized” the counts. The list of 2,034 cells considered “high quality” was obtained from file `metadata.tsv` included in the online benchmarking repository [154]. After removing peaks with fragments mapping to fewer than 20 cells, the final data set used in our analysis consisted of binary accessibility values for 2,034 cells and 126,719 peaks (4.6% overall rate of accessibility). Base-pair positions of the peaks were based on human genome assembly 19 (Genome Reference Consortium Human Build 37, February 2009).

In [104], a large, patient-specific batch effect was identified in the accessibility profiles for the HSC cells, and therefore steps were taken in [104] to normalize the accessibility data before performing PCA. We instead fit the topic model to the unnormalized binary accessibility values, in part to find out how well the topic model can cope with the complication of a batch effect. In agreement with [104], this batch effect is at least partly captured by the topics, although in our analysis the batch effect also appeared in MPP cells and, to a lesser extent, in CMP cells (Additional file 1: Figure S11).

### Fitting the topic models

In brief, we took the following steps to fit a topic model. All these steps are implemented in the R package `fastTopics`.

First, we fit a Poisson NMF model [37, 155],

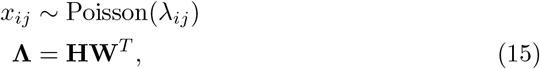

where **Λ** ∈ **R**^*n×m*^ is a matrix of the same dimension as **X** with entries *λ*_*ij*_ *≥* 0 giving the Poisson rates for the counts *x*_*ij*_. The parameters of the Poisson NMF model are stored as two matrices, **H** ∈ **R**^*n×K*^, **W** ∈ **R**^*m×K*^, with non-negative entries *h*_*ik*_, *w*_*jk*_. `fastTopics` has efficient implementations of algorithms for computing maximum-likelihood estimates (MLEs) of **W, H** [48]. Second, we recovered MLEs of **F, L** from MLEs of **W, H** by a simple reparameterization [48].

In an empirical comparison of Poisson NMF algorithms with count data sets, including scRNA-seq data sets [48], we found that a simple co-ordinate descent (CD) algorithm [156, 157], when accelerated with the extrapolation method of Ang and Gillis [158], almost always produced the best Poisson NMF (and topic model) fits, and in the least amount of time. To confirm this, we compared topic model fits obtained by running the same four algorithms that were compared in [48]—EM and CD, with and without extrapolation— on the PBMC data set, and assessed the quality of the fits. We evaluated the model fits in two ways: using the likelihood, and using the residuals of the KarushKuhn-Tucker (KKT) first-order conditions. (The residuals of the KKT system should vanish as the algorithm approaches maximum-likelihood estimates of **W, H**.) Following [48], to reduce the possibility that multiple optimizations converge to different local maxima of the likelihood, which could complicate these comparisons, we first ran 1,000 EM updates, then we examined the performance of the algorithms after this initialization phase (Additional file 1: Figures S13, S14). Consistent with [48], the extrapolated CD updates always produced the best fit, or at the very least a fit that was no worse than the other algorithms, and almost always converged on a solution more quickly than the other algorithms. Therefore, subsequently we used the extrapolated CD updates to fit the topic models.

In more detail, the pipeline for fitting topic models consisted of the following steps: (1) Initialize **W** using Topic-SCORE [159]; (2) perform 10 CD updates of **H**, with **W** fixed; (3) perform 1,000 EM updates (without extrapolation) to get close to a solution (“prefitting phase”); (4) run an additional 1,000 extrapolated CD updates to improve the fit (“refinement phase”); and (5) recover **F, L** from **W, H** by a simple transformation. The prefitting phase was implemented by calling `fit_poisson_nmf` from `fastTopics` with these settings: `numiter = 1000`, `method = “em”`, `control = list(numiter = 4)`. The refinement phase was implemented with a second call to `fit poisson nmf`, with `numiter = 1000`, `method = “scd”`, `control = list(numiter = 4,extrapolate = TRUE)`, in which the model fit was initialized using the fit from the prefitting phase. The topic model fit was recovered by calling `poisson2multinom` in `fastTopics`. Note that only the estimates of **L** were used in the GoM DE analysis.

For each data set, we fit topic models with different choices of *K* and compared the fits for each *K* by comparing their likelihoods (Additional file 1: Figure S15).

### Visualizing the membership proportions

The membership proportions matrix **L** can be viewed as an embedding of the cells *i* = 1, …, *n* in a continuous space with *K*− 1 dimensions [50]. (It is *K*−1 dimensions because of the constraint that the membership proportions for each cell must add up to 1.) A simple way to visualize this embedding in 2-d is to apply a nonlinear dimensionality reduction technique such as *t*-SNE [11, 160] or UMAP [12] to **L** ([49] used *t*-SNE). We have also found that plotting principal components (PCs) of the membership proportions can be an effective way to explore the structure inferred by the topic model (Additional file 1: Figures S1, S4). However, we view these visualization techniques as primarily for exploration, and a more powerful approach is to visualize all *K*− 1 dimensions simultaneously using a Structure plot [70, 71]. Here we describe some improvements to the Structure plot for better visualization. These improvements are implemented in the `structure plot` function in `fastTopics`.

When cells were labeled, we compared topics against labels by grouping the cells by these labels in the Structure plot. We then applied *t*-SNE to the **L** matrix, separately for each group, to arrange the cells on a line within each group. For this, we used the R package `Rtsne` [161]. (In `fastTopics`, we also implemented options to arrange the cells in each group using UMAP or PCA, but in our experience we found that UMAP and PCA produced “noisier” visualizations.)

Arranging the cells by 1-d *t*-SNE worked best for smaller groups of cells with less complex structure. For large groups of cells, or for unlabeled single-cell data sets, we randomly subsampled the cells to reduce *t*SNE runtime. (When cells number in the thousands, it is nearly impossible to distinguish individual cells in the Structure plot anyhow.) Even with this subsampling, the Structure plot sometimes did not show fine-scale substructures or rare cell types. Therefore, in more complex cases, we first subdivided the cells into smaller groups based on the membership proportions, then ran *t*-SNE on these smaller groups. These groups were either identified visually from PCs of **L**, or in a more automated way by running *k*-means on PCs of **L** (see [162]).

### Gene enrichment analysis based on differential accessibility of peaks connected to genes

Here we describe a simple approach to obtain genelevel statistics from the results of a differential accessibility analysis. This approach was applied in the topic modeling analysis of the Mouse Atlas kidney cells.

Cusanovich et al [97] used the Cicero co-accessibility predictions and the binarized single-cell ATAC-seq data to compute a “gene activity score” *R*_*ki*_ for each gene *k* and cell *i*. Here we have a related but different goal: we would like to use the results of the differential accessibility analysis, which generates differential accessibility estimates and related statistics for each peak and each topic, to rank genes according to their importance to a given topic. A difficulty, however, with ranking the genes is that the Cicero co-accessibility predictions are uncertain, and they are only partially informative about which peaks are relevant to a gene. In aggregate, however, the expectation is that the “most interesting” genes will be genes that are (a) predicted by Cicero to be connected many peaks that are differentially accessible and (b) the differences in accessibility are mainly in the same direction. This suggests an enrichment analysis in which, for each gene, we test for enrichment of differential acccessibility among the peaks connected to that gene. Here we describe a simple enrichment analysis for (a) and (b).

For (a), we computed a Bayes factor [163] measuring the support for the hypothesis that at least one of the peaks is differentially accessible (the LFC is not zero) against the null hypothesis that none of the peaks are differentially accessible. For (b), we computed the *average LFC* among all differentially accessible peaks (that is, peaks with nonzero LFC according to some significance criterion).

We implemented this gene enrichment analysis by running adaptive shrinkage [81] separately for each gene and topic. This had the benefit of adapting the shrinkage separately to each gene in each topic. In particular, in comparison to the usual adaptive shrinkage step for a GoM DE analysis (see above), it avoided overshrinking differences for genes exhibiting strong patterns of differential accessibility. We took the following steps to implement this adaptive shrinkage analysis. First, we ran function `ash` from the `ashr` package [144] once on the posterior mean l.e. LFC estimates 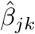 and their standard errors ŝ_*jk*_ for all topics *k* and all peaks *j*, with settings `mixcompdist = “normal”`, `method = “shrink”`. This was done only to determine the variances in the mixture prior and to get a “default” model fit to be used in the subsequent adaptive shrinkage analyses.

Next, we ran `ash` separately for gene and each topic *k* using the l.e. LFC estimates 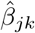 and standard errors ŝ_*jk*_ from the peaks *j* connected to that gene. We set the variances in the mixture prior to the variances determined from all the l.e. LFC estimates, and used `ash` settings `mixcompdist = “normal”` and `pointmass = FALSE`. One issue with running adaptive shrinkage using only the l.e. LFC estimates for the peaks connected to a gene is that some genes have few Cicero connections, leading to potentially unstable fits and unreliable posterior estimates. We addressed this issue by encouraging the fits toward the “default model” that was fitted to all genes and all topics; specifically, we set the Dirichlet prior on the mixture proportions to be Dirichlet(*α*_1_, …, *α*_*K*_) with prior sample sizes 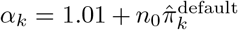, where here *K* denotes the number of components of the prior mixture (not the number of topics), 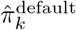 denotes the *k*th mixture proportion in the adaptive shrinkage prior for the fitted “default” model, and *n*_0_ = 20. This stabilized the fits for genes with few Cicero connections while still allowing some ability to adapt to genes with many connections.

Finally, we used the `logLR` output from `ash` as a measure of support for enrichment (this is the Bayes factor on the log-scale), and we computed the mean l.e. LFC as the average of the posterior mean estimates of the l.e. LFCs taken over all peaks *j* connected to the gene and with posterior *lfsr <* 0.05.

### Motif enrichment analysis for differentially accessible regions

We used HOMER [105] to identify transcription factor (TF) motifs enriched for differentially accessible regions, separately for each topic estimated from the single-cell ATAC-seq data. For each topic *k* = 1, …, *K*, we applied the HOMER Motif Analysis tool `findMotifsGenome.pl` to estimate motif enrichment in differentially accessible regions; specifically, we took “differentially accessible regions” to be those with *p*value less than 0.05 in the GoM DE analysis (Additional file 1: Figure S16). These differentially accessible regions were stored in a BED file `positions.bed`.

The exact call from the command-line shell was `findMotifsGenome.pl positions.bed hg19 homer -len 8,10,12 -size 200 -mis 2 -S 25 -p 4 -h`.

Note that the adaptive shrinkage step was skipped in the GoM DE analysis, so these are the *p*-values for the unmoderated l.e. LFC estimates. The reason for skipping the adaptive shrinkage step is that the shrinkage is performed uniformly for the LFC estimates for all regions, and since the vast majority of regions have l.e. LFC estimates are that are indistinguishable from zero, the result is that very few differentially accessible regions remain shrinkage.

### Gene sets

Human and mouse gene sets for the gene set enrichment analyses (GSEA) were compiled from the following gene set databases: NCBI BioSystems [164]; Pathway Commons [165, 166]; and MSigDB [167– 169], which includes Gene Ontology (GO) gene sets [94, 170]. Specifically, we downloaded `bsid2info.gz` and `biosystems gene.gz` from the NCBI FTP site (https://ftp.ncbi.nih.gov/gene) on March 22, 2020; `PathwayCommons12.All.hgnc.gmt.gz` from the Pathway Commons website (https://www.pathwaycommons.org) on March 20, 2020; and `msigdb_v7.2.xml.gz` from the MSigDB website (https://www.gsea-msigdb.org) on October 15, 2020. For the gene set enrichment analyses we also downloaded human and mouse gene information (“gene info”) files `Homo sapiens.gene info.gz` and `Mus musculus.gene info.gz` from the NCBI FTP site on October 15, 2020. Put together, we obtained 37,856 human gene sets and 33,380 mouse gene sets. In practice, we filtered gene sets based on certain criteria before running the GSEA. To facilitate integration of these gene sets into our analyses, we have compiled these gene sets into an R package [171].

### Gene set enrichment analysis

We took a simple multiple linear regression approach to the gene set enrichment analysis (GSEA), in which we modeled the l.e. LFC estimate for gene *i* in a given topic, here denoted by *y*_*i*_, as 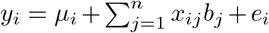, *e*_*i*_ *∼ N* (0, *σ*^2^), in which *x*_*ij*_ ∈ *{*0, 1*}* indicates gene set membership; *x*_*ij*_ = 1 if gene *i* belongs to gene set *j*, otherwise *x*_*ij*_ = 0. (We represented the geneset membership as a sparse matrix since most *x*_*ij*_’s are zero.) Here, *n* denotes the number of candidate gene sets, and *σ*^2^ is the residual variance to be estimated. The idea behind this simple approach was that the most relevant gene sets are those that best explain the log-fold changes *y*_*i*_, and therefore in the multiple regression we sought to identify these gene sets by finding coefficients *b*_*j*_ that were nonzero with high probability. See [172, 173] for similar ideas using logistic regression. Additionally, since many genes were typically differentially expressed in a given topic, modeling LFCs helped distinguish among DE genes that showed only a slight increase in expression versus those that were highly overexpressed [174, 175]. Of course, this simple multiple linear approach ignores uncertainty in the LFC estimates *y*_*i*_, which is accounted for in most gene set enrichment analyses. We addressed this issue by shrinking the l.e. LFC estimates *prior to running the GSEA*; that is, we took *y*_*i*_ to be the the posterior mean LFC estimate after applying adaptive shrinkage, as described above (see “Quantifying uncertainty and stabilizing LFC estimates”). The result was that genes that we were more uncertain about had have an l.e. LFC estimate *y*_*i*_ that was zero or near zero.

We implemented this multiple linear regression approach using SuSiE (`susieR` version 0.12.10) [176]. A benefit to using SuSiE is that it automatically organized similar or redundant gene sets into “credible sets” (CSs), making it easier to quickly recognize complementary gene sets; see [177–182] for related ideas.

In detail, the GSEA was performed as follows. We performed a separate GSEA for each topic, *k* = 1, …, *K*. Specifically, for each topic *k*, we ran the `susieR` function `susie` with the following options: `L = 10`, `intercept = TRUE, standardize = FALSE, estimate_residual_variance = TRUE, refine = FALSE compute_univariate_zscore = FALSE` and `min_abs_corr = 0`. We set `L = 10` so that SuSiE returned at most 10 credible sets. For a given topic *k*, we reported a gene set as being enriched if it was included in at least one CS. We organized the enriched gene sets by (95%) credible sets. We also recorded the Bayes factor for each CS, which gives a measure of the level of support for that CS. For each gene set included in a CS, we reported the posterior inclusion probability (PIP), and the posterior mean estimate of the regression coefficient *b*_*j*_. In the results, we refer to *b*_*j*_ as the “enrichment coefficient” for gene set *j* since it is an estimate of the expected increase in the l.e. LFC for genes that belong to gene set *j* relative to genes that do not belong to the gene set.

Often, a CS contained only one gene set, in which case the PIP for that gene set was close to 1. In several other cases, the CS contained multiple similar gene sets; in these cases, the smaller PIPs indicated that it was difficult to choose among the gene sets because they are similar to each other. (Note that the sum of the PIPs in a 95% CS should always be above 0.95 and less than 1.) Occasionally, SuSiE returned a CS with a small Bayes factor containing a very large number of gene sets. We excluded such CSs from the results.

When repeated these gene set enrichment analyses with two collections of gene sets: (1) all gene sets other than the MSigDB collections C1, C3, C4 and C6, and “archived” MSigDB gene sets; and (2) only gene sets from curated pathway databases, specifically Pathway Commons, NCBI BioSystems and “canonical pathways” (CP) in the MSigDB C2 collection, and Gene Ontology (GO) gene sets in the MSigDB C5 collection. In all cases, we removed gene sets with fewer than 10 genes and with more than 400 genes. Table 2 gives the exact number of gene sets included in each GSEA.

**Table 2.** Number of gene sets included in each GSEA.

| data set | all gene sets | curated only |
| --- | --- | --- |
| PBMC | 23,193 | 12,225 |
| mouse epithelial airway | 20,917 | 9,946 |
| —rare epithelial cell types only | 20,288 | 9,450 |

### Simulations

For evaluating the DE analysis methods, we generated matrices of UMI counts **X** ∈ **R**^*n×m*^ for *m* = 10,000 genes and *n* = 200 or *n* = 1,000 cells. We simulated the UMI counts *x*_*ij*_ from a Poisson NMF model (15) in which **W** and **H** were chosen to emulate UMI counts from scRNA-seq experiments.

The matrices **W** and **H** were generated as follows. First, for each cell *i*, we generated membership proportions *l*_*i*1_, …, *l*_*iK*_ then set *h*_*ik*_ = *s*_*i*_*l*_*ik*_, for *k* = 1, …, *K*, where *s*_*i*_ is the total UMI count. To simulate the wide range of total UMI counts often seen in scRNA-seq data sets, total UMI counts *s*_*i*_ were normally distributed on the log-scale, *s*_*i*_ = 10^*u*^*i, u*_*i*_ *∼ N* (0, 1*/*5), where *N* (*μ, σ*) denotes the univariate normal distribution with mean *μ* and standard deviation *σ*.

Membership proportions *l*_*ik*_ for each cell *i* were generated so as to obtain a wide range of mixed memberships, according to the following procedure: the number of nonzero proportions was set to *K*^*t*^ ∈ *{*1, …, *K}* with probability 2^− *I K*^ ; the *K*^*t*^ selected topics *t*_1_, …, *t*_*K I*_ ⊆ { 1, …, *K* } were drawn uniformly at random (without replacement) from 1, …, *K*; then the membership proportions for the selected topics were set to 1 when *K*^*t*^ = 1, or, when *K*^*t*^ *>* 1, they were drawn from the Dirichlet distribution with shape parameters 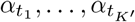 .

Expression rates *w*_*jk*_ were generated so as to emulate the wide distribution of gene expression levels observed in single-cell data sets, and to allow for differences in expression rates among topics. The procedure for generating the expression rates for each gene *j* was as follows: with probability 0.5, the expression rates were the same across all topics, and were generated as 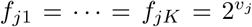,*v*_*j*_ *∼ N* ( − 4, 2). Otherwise, with probability 0.5, the expression rates were the same in all topics except for one topic. The differing topic *k*^*t*^ was chosen uniformly at random from 1, …, *K*, then the expression rate for topic *k*^*t*^ was set to 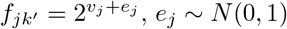 . As a result, the expression rates were roughly normally distributed on the log-scale, and the expression differences were also normally distributed on the log-scale. About half of genes had an expression difference among the topics.

Using this simulation procedure, we generated three collections of data sets. The simulation settings were altered slightly for each collection. In the first, data sets were simulated with *K* = 2, *α* = (1*/*100, 1*/*100), *n* = 200 so that most membership proportions were equal or very close to 0 or 1. In the second, we used *K* = 2, *α* = (1, 1), *n* = 200 to allow for a range of mixed memberships. In the third, we generated data sets with *K* = 6, *α* = (1, …, 1), *n* = 1,000.

For the data sets simulated with *K* = 2, *α* = (1*/*100, 1*/*100), the cells could essentially be subdivided into two groups. Therefore, we ran `MAST` [84, 183] and `DESeq2` [78, 83] to test for genes that were differentially expressed between the two groups. `MAST` (R package version 1.20.0) was called via the `FindMarkers` interface in `Seurat` [25] (`Seurat` 4.0.3, `SeuratObject` 4.0.2) with the following settings: `ident.1 = “2”`,`ident.2 = NULL, test.use = “MAST”, logfc.threshold = 0, min.pct = 0`. `DESeq` was called from the `DESeq2` R package (version 1.34.0) using settings recommended in the package vignette: `test = “LRT”, reduced = ∼1, useT = TRUE, minmu = 1e-6, minReplicatesForReplace = Inf`. Size factors were calculated using the `calculateSumFactors` method from `scran` version 1.22.1 [23]. The LFC estimates returned by `DESeq` were subsequently revised using adaptive shrinkage [81] by calling `lfcShrink` in DESeq2 with `type = “ashr”, svalue = TRUE`. (As in the GoM DE analysis, the DESeq2 posterior *z*-scores were defined as the posterior means divided by the posterior standard errors returned by the adaptive shrinkage.)

To perform the GoM DE analysis in each of the simulations, we first fit a Poisson NMF model to the simulated counts **X** using `fit_poisson_nmf` from the `fastTopics` R package [48, 184] (version 0.6-97). The loadings matrix **H** was fixed to the matrix used to simulate the data, and **W** was estimated by running 40 co-ordinate ascent updates on **W** alone (`update.loadings = NULL, method = “scd”, numiter = 40`). The equivalent topic model fit was then recovered. Three GoM DE analyses were performed using the `de_analysis` function from the `fastTopics` R package, with the topic model fit provided as input: one analysis without adaptive shrinkage (`shrink.method = “none”`), and two analyses with adaptive shrinkage (`shrink.method = “ash”, ashr` version 2.2-51 [144]) in which the MCMC was initialized with different pseudorandom number generator states. In all three runs, posterior calculations were performed with *n*_*s*_ = 10,000, *ε* = 0.01. Comparison of the two MCMC runs (with adaptive shrinkage) suggested that *n*_*s*_ = 10, 000 was sufficient to obtain reasonably accurate posterior estimates in these simulations (Additional file 1: Figure S2).

### Computing environment

Most computations on real data sets were run in R 3.5.1 [185], linked to the OpenBLAS 0.2.19 optimized numerical libraries, on Linux machines (Scientific Linux 7.4) with Intel Xeon E5-2680v4 (“Broadwell”) processors. For performing the Poisson NMF optimization, which included some multithreaded computations, as many as 8 CPUs and 16 GB of memory were used. The DESeq2 analysis of the PBMC data was performed in R 4.1.0, using 4 CPUs and 264 GB of memory. The evaluation of the DE analysis methods in simulated data sets was performed in R 4.1.0, using as many and 8 CPUs as 24 GB of memory. More details about the computing environment, including the R packages used, are recorded in the workflowr pages in the companion code repositories [186, 187].

## Supporting information

Additional file 4: Interactive volcano plots for PBMC data

Additional file 5: Interactive volcano plots for epithelial airway data

## Acknowledgements

We thank Mihai Anitescu, Nicolas Chevrier, Michihiro Takahama, Kushal Dey, Adam Gruenbaum, Youngseok Kim, John Novembre, Alan Selewa, Katie Rhodes, Yusha Liu, Dongyue Xie, Zepeng Mu and Jason Willwerscheid for their valuable input. We also thank the staff at the Research Computing Center for providing the high-performance computing resources used to implement the numerical experiments.

## Availability of data and materials

The fastTopics R package is available on GitHub (https://github.com/stephenslab/fastTopics) and CRAN (https://cran.r-project.org/package=fastTopics). A Seurat wrapper for fastTopics is available from https://github.com/stephenslab/seurat-wrappers. The data sets supporting the conclusions of this article are available in Zenodo repositories [186, 187]. These Zenodo repositories also include the source code implementing the analyses and workflowr websites [188] for browsing the code and results. Permission to use the source code in these repositories is granted under the MIT license. Numerical implementations of the contributed statistical methods, including tools for visualizing the results generated by these methods, are available from the fastTopics R package [48, 184] under the MIT license. The gene sets used in the GSEA were compiled into an R package [171], also distributed under the MIT license. All data sets used in the study were obtained from public sources [148, 149, 152, 153]. A description of how these data sets were used is provided in Methods.

## Ethics approval and consent to participate

Not applicable.

## Consent for publication

Not applicable.

## Authors’ contributions

PC implemented the methods based on mathematical derivations by PC and MS, with contributions from KL, AS and SP. PC performed empirical evaluations of the methods using simulated data sets. PC and KL analyzed the single-cell data sets. PC wrote the draft manuscript, with contributions, revisions and discussion by all authors. All authors approved the final manuscript.

## References

1. Kharchenko, P.V.: The triumphs and limitations of computational methods for scRNA-seq. Nature Methods 18(7), 723–732 (2021)

2. Stegle, O., Teichmann, S.A., Marioni, J.C.: Computational and analytical challenges in single-cell transcriptomics. Nature Reviews Genetics 16(3), 133–145 (2015)

3. Wagner, A., Regev, A., Yosef, N.: Revealing the vectors of cellular identity with single-cell genomics. Nature Biotechnology 34(11), 1145–1160 (2016)

4. Duo, A., Robinson, M.D., Soneson, C.: A systematic performance evaluation of clustering methods for single-cell RNA-seq data. F1000Research 7, 1141 (2018)

5. Freytag, S., Tian, L., Lönnstedt, I., Ng, M., Bahlo, M.: Comparison of clustering tools in R for medium-sized 10x Genomics single-cell RNA-sequencing data. F1000Research 7, 1297 (2018)

6. Kiselev, V.Y., Andrews, T.S., Hemberg, M.: Challenges in unsupervised clustering of single-cell RNA-seq data. Nature Reviews Genetics 20(5), 273–282 (2019)

7. Sun, S., Zhu, J., Ma, Y., Zhou, X.: Accuracy, robustness and scalability of dimensionality reduction methods for single-cell RNA-seq analysis. Genome Biology 20, 269 (2019)

8. Townes, F.W., Hicks, S.C., Aryee, M.J., Irizarry, R.A.: Feature selection and dimension reduction for single-cell RNA-Seq based on a multinomial model. Genome Biology 20, 295 (2019)

9. Hotelling, H.: Analysis of a complex of statistical variables into principal components. Journal of Educational Psychology 24(6), 417–441 (1933)

10. Risso, D., Perraudeau, F., Gribkova, S., Dudoit, S., Vert, J.-P.: A general and flexible method for signal extraction from single-cell RNA-seq data. Nature Communications 9, 284 (2018)

11. van der Maaten, L., Hinton, G.: Visualizing data using t-SNE. Journal of Machine Learning Research 9(86), 2579–2605 (2008)

12. McInnes, L., Healy, J., Saul, N., Großberger, L.: UMAP: uniform manifold approximation and projection. Journal of Open Source Software 3(29), 861 (2018)

13. Becht, E., McInnes, L., Healy, J., Dutertre, C.-A., Kwok, I.W.H., Ng, L.G., Ginhoux, F., Newell, E.W.: Dimensionality reduction for visualizing single-cell data using UMAP. Nature Biotechnology 37, 38–44 (2019)

14. Cooley, S.M., Hamilton, T., Aragones, S.D., Ray, J.C.J., Deeds, E.J.: A novel metric reveals previously unrecognized distortion in dimensionality reduction of scRNA-seq data. bioRxiv (2022). doi:10.1101/689851

15. Chari, T., Pachter, L.: The specious art of single-cell genomics. PLoS Computational Biology 19(8), 1011288 (2023)

16. Kobak, D., Berens, P.: The art of using t-SNE for single-cell transcriptomics. Nature Communications 10(1), 5416 (2019)

17. Kobak, D., Linderman, G.C.: Initialization is critical for preserving global data structure in both t-SNE and UMAP. Nature Biotechnology 39(2), 156–157 (2021)

18. Wattenberg, M., Viégas, F., Johnson, I.: How to use t-SNE effectively. Distill (2016). doi:10.23915/distill.00002

19. Granja, J.M., Corces, M.R., Pierce, S.E., Bagdatli, S.T., Choudhry, H., Chang, H.Y., Greenleaf, W.J.: ArchR is a scalable software package for integrative single-cell chromatin accessibility analysis. Nature Genetics 53(3), 403–411 (2021)

20. Heiser, C.N., Lau, K.S.: A quantitative framework for evaluating single-cell data structure preservation by dimensionality reduction techniques. Cell Reports 31(5), 107576 (2020)

21. Linderman, G.C., Rachh, M., Hoskins, J.G., Steinerberger, S., Kluger, Y.: Fast interpolation-based t-SNE for improved visualization of single-cell RNA-seq data. Nature Methods 16(3), 243–245 (2019)

22. Linderman, G.C., Steinerberger, S.: Clustering with t-SNE, provably. SIAM Journal on Mathematics of Data Science 1(2), 313–332 (2019)

23. Lun, A.T.L., McCarthy, D.J., Marioni, J.C.: A step-by-step workflow for low-level analysis of single-cell RNA-seq data with Bioconductor. F1000Research 5, 2122 (2016)

24. McCarthy, D.J., Campbell, K.R., Lun, A.T.L., Wills, Q.F.: Scater: Pre-processing, quality control, normalization and visualization of single-cell RNA-seq data in R. Bioinformatics 33(8), 1179–1186 (2017)

25. Stuart, T., Butler, A., Hoffman, P., Hafemeister, C., Papalexi, E., Mauck, W.M., Hao, Y., Stoeckius, M., Smibert, P., Satija, R.: Comprehensive integration of single-cell data. Cell 177(7), 1888–1902 (2019)

26. Butler, A., Hoffman, P., Smibert, P., Papalexi, E., Satija, R.: Integrating single-cell transcriptomic data across different conditions, technologies, and species. Nature Biotechnology 36(5), 411–420 (2018)

27. Stuart, T., Srivastava, A., Madad, S., Lareau, C.A., Satija, R.: Single-cell chromatin state analysis with Signac. Nature Methods 18(11), 1333–1341 (2021)

28. Fang, R., Preissl, S., Li, Y., Hou, X., Lucero, J., Wang, X., Motamedi, A., Shiau, A.K., Zhou, X., Xie, F., Mukamel, E.A., Zhang, K., Zhang, Y., Behrens, M.M., Ecker, J.R., Ren, B.: Comprehensive analysis of single cell ATAC-seq data with SnapATAC. Nature Communications 12, 1337 (2021)

29. Zheng, G.X.Y., Terry, J.M., Belgrader, P., Ryvkin, P., Bent, Z.W., Wilson, R., Ziraldo, S.B., Wheeler, T.D., McDermott, G.P., Zhu, J., Gregory, M.T., Shuga, J., Montesclaros, L., Underwood, J.G., Masquelier, D.A., Nishimura, S.Y., Schnall-Levin, M., Wyatt, P.W., Hindson, C.M., Bharadwaj, R., Wong, A., Ness, K.D., Beppu, L.W., Deeg, H.J., McFarland, C., Loeb, K.R., Valente, W.J., Ericson, N.G., Stevens, E.A., Radich, J.P., Mikkelsen, T.S., Hindson, B.J., Bielas, J.H.: Massively parallel digital transcriptional profiling of single cells. Nature Communications 8, 14049 (2017)

30. Brunet, J.-P., Tamayo, P., Golub, T.R., Mesirov, J.P.: Metagenes and molecular pattern discovery using matrix factorization. Proceedings of the National Academy of Sciences 101(12), 4164–4169 (2004)

31. Donoho, D., Stodden, V.: When does non-negative matrix factorization give a correct decomposition into parts? In: Proceedings of the 16th International Conference on Neural Information Processing Systems, pp. 1141–1148. MIT Press, Cambridge, MA, USA (2003)

32. Durif, G., Modolo, L., Mold, J.E., Lambert-Lacroix, S., Picard, F.: Probabilistic count matrix factorization for single cell expression data analysis. Bioinformatics 35(20), 4011–4019 (2019)

33. Gong, W., Rasmussen, T.L., Singh, B.N., Koyano-Nakagawa, N., Pan, W., Garry, D.J.: Dpath software reveals hierarchical haemato-endothelial lineages of Etv2 progenitors based on single-cell transcriptome analysis. Nature Communications 8, 14362 (2017)

34. Elyanow, R., Dumitrascu, B., Engelhardt, B.E., Raphael, B.J.: netNMF-sc: leveraging gene–gene interactions for imputation and dimensionality reduction in single-cell expression analysis. Genome Research 30(2), 195–204 (2020)

35. Ho, Y.-J., Anaparthy, N., Molik, D., Mathew, G., Aicher, T., Patel, A., Hicks, J., Hammell, M.G.: Single-cell RNA-seq analysis identifies markers of resistance to targeted BRAF inhibitors in melanoma cell populations. Genome Research 28(9), 1353–1363 (2018)

36. Kotliar, D., Veres, A., Nagy, M.A., Tabrizi, S., Hodis, E., Melton, D.A., Sabeti, P.C.: Identifying gene expression programs of cell-type identity and cellular activity with single-cell RNA-seq. eLife 8, 43803 (2019)

37. Lee, D.D., Seung, H.S.: Learning the parts of objects by non-negative matrix factorization. Nature 401(6755), 788–791 (1999)

38. Levitin, H.M., Yuan, J., Cheng, Y.L., Ruiz, F.J., Bush, E.C., Bruce, J.N., Canoll, P., Iavarone, A., Lasorella, A., Blei, D.M., Sims, P.A.: De novo gene signature identification from single-cell RNA-seq with hierarchical poisson factorization. Molecular Systems Biology 15, 8557 (2019)

39. Welch, J.D., Kozareva, V., Ferreira, A., Vanderburg, C., Martin, C., Macosko, E.Z.: Single-cell multi-omic integration compares and contrasts features of brain cell identity. Cell 177(7), 1873–188717 (2019)

40. Shao, C., Höfer, T.: Robust classification of single-cell transcriptome data by nonnegative matrix factorization. Bioinformatics 33(2), 235–242 (2016)

41. Sun, S., Chen, Y., Liu, Y., Shang, X.: A fast and efficient count-based matrix factorization method for detecting cell types from single-cell RNAseq data. BMC Systems Biology 13, 28 (2019)

42. Venkatasubramanian, M., Chetal, K., Schnell, D.J., Atluri, G., Salomonis, N.: Resolving single-cell heterogeneity from hundreds of thousands of cells through sequential hybrid clustering and NMF. Bioinformatics 36(12), 3773–3780 (2020)

43. Zhang, S., Yang, L., Yang, J., Lin, Z., Ng, M.K.: Dimensionality reduction for single cell RNA sequencing data using constrained robust non-negative matrix factorization. NAR Genomics and Bioinformatics 2(3) (2020)

44. Pierson, E., Yau, C.: ZIFA: Dimensionality reduction for zero-inflated single-cell gene expression analysis. Genome Biology 16, 241 (2015)

45. Blei, D.M., Ng, A.Y., Jordan, M.I.: Latent Dirichlet allocation. Journal of Machine Learning Research 3, 993–1022 (2003)

46. DuVerle, D.A., Yotsukura, S., Nomura, S., Aburatani, H., Tsuda, K.: CellTree: an R/bioconductor package to infer the hierarchical structure of cell populations from single-cell RNA-seq data. BMC Bioinformatics 17, 363 (2016)

47. Dey, K.K., Hsiao, C.J., Stephens, M.: Visualizing the structure of RNA-seq expression data using grade of membership models. PLoS Genetics 13(3), 1006599 (2017)

48. Carbonetto, P., Sarkar, A., Wang, Z., Stephens, M.: Non-negative matrix factorization algorithms greatly improve topic model fits. arXiv 2105.13440 (2021)

49. González-Blas, C., Minnoye, L., Papasokrati, D., Aibar, S., Hulselmans, G., Christiaens, V., Davie, K., Wouters, J., Aerts, S.: cisTopic: cis-regulatory topic modeling on single-cell ATAC-seq data. Nature Methods 16(5), 397–400 (2019)

50. Hofmann, T.: Probabilistic latent semantic indexing. In: Proceedings of the 22nd Annual International ACM SIGIR Conference, pp. 50–57 (1999)

51. Bielecki, P., Riesenfeld, S.J., Hütter, J.-C., Torlai Triglia, E., Kowalczyk, M.S., Ricardo-Gonzalez, R.R., Lian, M., Amezcua Vesely, M.C., Kroehling, L., Xu, H., Slyper, M., Muus, C., Ludwig, L.S., Christian, E., Tao, L., Kedaigle, A.J., Steach, H.R., York, A.G., Skadow, M.H., Yaghoubi, P., Dionne, D., Jarret, A., McGee, H.M., Porter, C.B.M., Licona-Limón, P., Bailis, W., Jackson, R., Gagliani, N., Gasteiger, G., Locksley, R.M., Regev, A., Flavell, R.A.: Skin-resident innate lymphoid cells converge on a pathogenic effector state. Nature 592, 128–132 (2021)

52. Housman, G., Briscoe, E., Gilad, Y.: Evolutionary insights into primate skeletal gene regulation using a comparative cell culture model. PLoS Genetics 18(3), 1010073 (2022)

53. Hung, A., Housman, G., Briscoe, E.A., Cuevas, C., Gilad, Y.: Characterizing gene expression in an in vitro biomechanical strain model of joint health. F1000Research 11, 296 (2022)

54. Rhodes, K., Barr, K.A., Popp, J.M., Strober, B.J., Battle, A., Gilad, Y.: Human embryoid bodies as a novel system for genomic studies of functionally diverse cell types. eLife 11, 71361 (2022)

55. Schenkel, J.M., Herbst, R.H., Canner, D., Li, A., Hillman, M., Shanahan, S.-L., Gibbons, G., Smith, O.C., Kim, J.Y., Westcott, P., Hwang, W.L., Freed-Pastor, W.A., Eng, G., Cuoco, M.S., Rogers, P., Park, J.K., Burger, M.L., Rozenblatt-Rosen, O., Cong, L., Pauken, K.E., Regev, A., Jacks, T.: Conventional type I dendritic cells maintain a reservoir of proliferative tumor-antigen specific TCF-1+ CD8+ T cells in tumor-draining lymph nodes. Immunity 54(10), 2338–23536 (2021)

56. Xu, H., Ding, J., Porter, C.B.M., Wallrapp, A., Tabaka, M., Ma, S., Fu, S., Guo, X., Riesenfeld, S.J., Su, C., Dionne, D., Nguyen, L.T., Lefkovith, A., Ashenberg, O., Burkett, P.R., Shi, H.N., Rozenblatt-Rosen, O., Graham, D.B., Kuchroo, V.K., Regev, A., Xavier, R.J.: Transcriptional atlas of intestinal immune cells reveals that neuropeptide α-CGRP modulates group 2 innate lymphoid cell responses. Immunity 51(4), 696–708 (2019)

57. Ding, C., Li, T., Peng, W.: On the equivalence between non-negative matrix factorization and probabilistic latent semantic indexing. Computational Statistics and Data Analysis 52(8), 3913–3927 (2008)

58. Gaussier, E., Goutte, C.: Relation between PLSA and NMF and implications. In: Proceedings of the 28th Annual International ACM SIGIR Conference, pp. 601–602 (2005)

59. Gillis, N.: Nonnegative Matrix Factorization. Society for Industrial and Applied Mathematics, Philadelphia, PA (2021)

60. Kim, J., Park, H.: Sparse nonnegative matrix factorization for clustering. Technical report, Georgia Institute of Technology (2008)

61. Soneson, C., Robinson, M.D.: Bias, robustness and scalability in single-cell differential expression analysis. Nature Methods 15(4), 255–261 (2018)

62. Wang, T., Li, B., Nelson, C.E., Nabavi, S.: Comparative analysis of differential gene expression analysis tools for single-cell RNA sequencing data. BMC Bioinformatics 20, 40 (2019)

63. Erosheva, E., Fienberg, S., Lafferty, J.: Mixed-membership models of scientific publications. Proceedings of the National Academy of Sciences 101(Supplement 1), 5220–5227 (2004)

64. Abdelaal, T., Michielsen, L., Cats, D., Hoogduin, D., Mei, H., Reinders, M.J.T., Mahfouz, A.: A comparison of automatic cell identification methods for single-cell RNA sequencing data. Genome Biology 20, 194 (2019)

65. Diaz-Mejia, J.J., Meng, E.C., Pico, A.R., MacParland, S.A., Ketela, T., Pugh, T.J., Bader, G.D., Morris, J.H.: Evaluation of methods to assign cell type labels to cell clusters from single-cell RNA-sequencing data. F1000Research 8, 296 (2019)

66. Blei, D.M., Lafferty, J.D.: Topic models. In: Srivastava, A.N., Sahami, M. (eds.) Text Mining: Classification, Clustering, and Applications, pp. 71–94. Chapman and Hall/CRC, Boca Raton, FL (2009)

67. Griffiths, T.L., Steyvers, M.: Finding scientific topics. Proceedings of the National Academy of Sciences 101(Supplement 1), 5228–5235 (2004)

68. Hofmann, T.: Unsupervised learning by probabilistic latent semantic analysis. Machine Learning 42(1), 177–196 (2001)

69. Sarkar, A., Stephens, M.: Separating measurement and expression models clarifies confusion in single-cell RNA sequencing analysis. Nature Genetics 53(6), 770–777 (2021)

70. Rosenberg, N.A.: Genetic structure of human populations. Science 298(5602), 2381–2385 (2002)

71. Rosenberg, N.A.: distruct: a program for the graphical display of population structure. Molecular Ecology Notes 4(1), 137–138 (2004)

72. Haak, W., Lazaridis, I., Patterson, N., Rohland, N., Mallick, S., Llamas, B., Brandt, G., Nordenfelt, S., Harney, E., Stewardson, K., Fu, Q., Mittnik, A., Bánffy, E., Economou, C., Francken, M., Friederich, S., Pena, R.G., Hallgren, F., Khartanovich, V., Khokhlov, A., Kunst, M., Kuznetsov, P., Meller, H., Mochalov, O., Moiseyev, V., Nicklisch, N., Pichler, S.L., Risch, R., Rojo Guerra, M.A., Roth, C., Szécsényi-Nagy, A., Wahl, J., Meyer, M., Krause, J., Brown, D., Anthony, D., Cooper, A., Alt, K.W., Reich, D.: Massive migration from the steppe was a source for Indo-European languages in europe. Nature 522(7555), 207–211 (2015)

73. Pereira, B.I., De Maeyer, R.P.H., Covre, L.P., Nehar-Belaid, D., Lanna, A., Ward, S., Marches, R., Chambers, E.S., Gomes, D.C.O., Riddell, N.E., Maini, M.K., Teixeira, V.H., Janes, S.M., Gilroy, D.W., Larbi, A., Mabbott, N.A., Ucar, D., Kuchel, G.A., Henson, S.M., Strid, J., Lee, J.H., Banchereau, J., Akbar, A.N.: Sestrins induce natural killer function in senescent-like CD8+ T cells. Nature Immunology 21(6), 684–694 (2020)

74. Ashuach, T., Reidenbach, D.A., Gayoso, A., Yosef, N.: PeakVI: a deep generative model for single-cell chromatin accessibility analysis. Cell Reports Methods 2(3), 100182 (2022)

75. Marioni, J.C., Mason, C.E., Mane, S.M., Stephens, M., Gilad, Y.: RNA-seq: an assessment of technical reproducibility and comparison with gene expression arrays. Genome Research 18(9), 1509–1517 (2008)

76. Robinson, M.D., Smyth, G.K.: Moderated statistical tests for assessing differences in tag abundance. Bioinformatics 23(21), 2881–2887 (2007)

77. Robinson, M.D., McCarthy, D.J., Smyth, G.K.: edgeR: a Bioconductor package for differential expression analysis of digital gene expression data. Bioinformatics 26(1), 139–140 (2009)

78. Anders, S., Huber, W.: Differential expression analysis for sequence count data. Genome Biology 11, 106 (2010)

79. Love, M.I., Huber, W., Anders, S.: Moderated estimation of fold change and dispersion for RNA-seq data with DESeq2. Genome Biology 15(12), 550 (2014)

80. Cable, D.M., Murray, E., Shanmugam, V., Zhang, S., Zou, L.S., Diao, M., Chen, H., Macosko, E.Z., Irizarry, R.A., Chen, F.: Cell type-specific inference of differential expression in spatial transcriptomics. Nature Methods 19(9), 1076–1087 (2022)

81. Stephens, M.: False discovery rates: a new deal. Biostatistics 18(2), 275–294 (2016)

82. Becker-Herman, S., Lantner, F., Shachar, I.: Id2 negatively regulates B cell differentiation in the spleen. Journal of Immunology 168(11), 5507–5513 (2002)

83. Zhu, A., Ibrahim, J.G., Love, M.I.: Heavy-tailed prior distributions for sequence count data: removing the noise and preserving large differences. Bioinformatics 35(12), 2084–2092 (2019)

84. Finak, G., McDavid, A., Yajima, M., Deng, J., Gersuk, V., Shalek, A.K., Slichter, C.K., Miller, H.W., McElrath, M.J., Prlic, M., Linsley, P.S., Gottardo, R.: MAST: a flexible statistical framework for assessing transcriptional changes and characterizing heterogeneity in single-cell RNA sequencing data. Genome Biology 16, 278 (2015)

85. Squair, J.W., Gautier, M., Kathe, C., Anderson, M.A., James, N.D., Hutson, T.H., Hudelle, R., Qaiser, T., Matson, K.J.E., Barraud, Q., Levine, A.J., La Manno, G., Skinnider, M.A., Courtine, G.: Confronting false discoveries in single-cell differential expression. Nature Communications 12, 5692 (2021)

86. Montoro, D.T., Haber, A.L., Biton, M., Vinarsky, V., Lin, B., Birket, S.E., Yuan, F., Chen, S., Leung, H.M., Villoria, J., Rogel, N., Burgin, G., Tsankov, A.M., Waghray, A., Slyper, M., Waldman, J., Nguyen, L., Dionne, D., Rozenblatt-Rosen, O., Tata, P.R., Mou, H., Shivaraju, M., Bihler, H., Mense, M., Tearney, G.J., Rowe, S.M., Engelhardt, J.F., Regev, A., Rajagopal, J.: A revised airway epithelial hierarchy includes CFTR-expressing ionocytes. Nature 560(7718), 319–324 (2018)

87. Rosvall, M., Bergstrom, C.T.: Maps of random walks on complex networks reveal community structure. Proceedings of the National Academy of Sciences 105(4), 1118–1123 (2008)

88. Coifman, R.R., Lafon, S., Lee, A.B., Maggioni, M., Nadler, B., Warner, F., Zucker, S.W.: Geometric diffusions as a tool for harmonic analysis and structure definition of data: diffusion maps. Proceedings of the National Academy of Sciences 102(21), 7426–7431 (2005)

89. Ruiz García, S., Deprez, M., Lebrigand, K., Cavard, A., Paquet, A., Arguel, M.-J., Magnone, V., Truchi, M., Caballero, I., Leroy, S., Marquette, C.-H., Marcet, B., Barbry, P., Zaragosi, L.-E.: Novel dynamics of human mucociliary differentiation revealed by single-cell RNA sequencing of nasal epithelial cultures. Development 146(20) (2019)

90. Barkauskas, C.E., Chung, M.-I., Fioret, B., Gao, X., Katsura, H., Hogan, B.L.M.: Lung organoids: current uses and future promise. Development 144(6), 986–997 (2017)

91. Rawlins, E.L., Okubo, T., Xue, Y., Brass, D.M., Auten, R.L., Hasegawa, H., Wang, F., Hogan, B.L.M.: The role of Scgb1a1+ clara cells in the long-term maintenance and repair of lung airway, but not elveolar, epithelium. Cell Stem Cell 4(6), 525–534 (2009)

92. Spassky, N., Meunier, A.: The development and functions of multiciliated epithelia. Nature Reviews Molecular Cell Biology 18(7), 423–436 (2017)

93. Zhao, H., Zhu, L., Zhu, Y., Cao, J., Li, S., Huang, Q., Xu, T., Huang, X., Yan, X., Zhu, X.: The cep63 paralogue deup1 enables massive de novo centriole biogenesis for vertebrate multiciliogenesis. Nature Cell Biology 15(12), 1434–1444 (2013)

94. The Gene Ontology Consortium: The Gene Ontology resource: enriching a GOld mine. Nucleic Acids Research 49(D1), 325–334 (2020)

95. Pliner, H.A., Packer, J.S., McFaline-Figueroa, J.L., Cusanovich, D.A., Daza, R.M., Aghamirzaie, D., Srivatsan, S., Qiu, X., Jackson, D., Minkina, A., Adey, A.C., Steemers, F.J., Shendure, J., Trapnell, C.: Cicero predicts cis-regulatory DNA interactions from single-cell chromatin accessibility data. Molecular Cell 71(5), 858–8718 (2018)

96. Wu, C., Tao, Y., Li, N., Fei, J., Wang, Y., Wu, J., Gu, H.F.: Prediction of cellular targets in diabetic kidney diseases with single-cell transcriptomic analysis of db/db mouse kidneys. Journal of Cell Communication and Signaling (2022)

97. Cusanovich, D.A., Hill, A.J., Aghamirzaie, D., Daza, R.M., Pliner, H.A., Berletch, J.B., Filippova, G.N., Huang, X., Christiansen, L., DeWitt, W.S., Lee, C., Regalado, S.G., Read, D.F., Steemers, F.J., Disteche, C.M., Trapnell, C., Shendure, J.: A single-cell atlas of in vivo mammalian chromatin accessibility. Cell 174(5), 1309–1324 (2018)

98. Der, E., Ranabothu, S., Suryawanshi, H., Akat, K.M., Clancy, R., Morozov, P., Kustagi, M., Czuppa, M., Izmirly, P., Belmont, H.M., Wang, T., Jordan, N., Bornkamp, N., Nwaukoni, J., Martinez, J., Goilav, B., Buyon, J.P., Tuschl, T., Putterman, C.: Single cell RNA sequencing to dissect the molecular heterogeneity in lupus nephritis. JCI Insight 2(9), 93009 (2017)

99. Grassmeyer, J., Mukherjee, M., DeRiso, J., Hettinger, C., Bailey, M., Sinha, S., Visvader, J.E., Zhao, H., Fogarty, E., Surendran, K.: Elf5 is a principal cell lineage specific transcription factor in the kidney that contributes to Aqp 2 and Avpr 2 gene expression. Developmental Biology 424(1), 77–89 (2017)

100. Park, J., Shrestha, R., Qiu, C., Kondo, A., Huang, S., Werth, M., Li, M., Barasch, J., Suszták, K.: Single-cell transcriptomics of the mouse kidney reveals potential cellular targets of kidney disease. Science 360(6390), 758–763 (2018)

101. Ghezzi, C., Loo, D.D.F., Wright, E.M.: Physiology of renal glucose handling via SGLT1, SGLT2 and GLUT2. Diabetologia 61(10), 2087–2097 (2018)

102. Thiagarajan, R.D., Georgas, K.M., Rumballe, B.A., Lesieur, E., Chiu, H.S., Taylor, D., Tang, D.T.P., Grimmond, S.M., Little, M.H.: Identification of anchor genes during kidney development defines ontological relationships, molecular subcompartments and regulatory pathways. PLoS ONE 6(2), 17286 (2011)

103. Gopal, E., Umapathy, N.S., Martin, P.M., Ananth, S., Gnana-Prakasam, J.P., Becker, H., Wagner, C.A., Ganapathy, V., Prasad, P.D.: Cloning and functional characterization of human SMCT2 (SLC5A12) and expression pattern of the transporter in kidney. Biochimica et Biophysica Acta—Biomembranes 1768(11), 2690–2697 (2007)

104. Buenrostro, J.D., Corces, M.R., Lareau, C.A., Wu, B., Schep, A.N., Aryee, M.J., Majeti, R., Chang, H.Y., Greenleaf, W.J.: Integrated single-cell analysis maps the continuous regulatory landscape of human hematopoietic differentiation. Cell 173(6), 1535–1548 (2018)

105. Heinz, S., Benner, C., Spann, N., Bertolino, E., Lin, Y.C., Laslo, P., Cheng, J.X., Murre, C., Singh, H., Glass, C.K.: Simple combinations of lineage-determining transcription factors prime cis-regulatory elements required for macrophage and B cell identities. Molecular Cell 38(4), 576–589 (2010)

106. Booeshaghi, A.S., Pachter, L.: Normalization of single-cell RNA-seq counts by log(x + 1) or log(1 + x). Bioinformatics 37(15), 2223–2224 (2021)

107. Lun, A.: Overcoming systematic errors caused by log-transformation of normalized single-cell RNA sequencing data. bioRxiv (2018). doi:10.1101/404962

108. Warton, D.I.: Why you cannot transform your way out of trouble for small counts. Biometrics 74, 362–368 (2018)

109. Amezquita, R.A., Lun, A.T.L., Becht, E., Carey, V.J., Carpp, L.N., Geistlinger, L., Marini, F., Rue-Albrecht, K., Risso, D., Soneson, C., Waldron, L., Pages, H., Smith, M.L., Huber, W., Morgan, M., Gottardo, R., Hicks, S.C.: Orchestrating single-cell analysis with Bioconductor. Nature Methods 17(2), 137–145 (2020)

110. Barkas, N., Petukhov, V., Nikolaeva, D., Lozinsky, Y., Demharter, S., Khodosevich, K., Kharchenko, P.V.: Joint analysis of heterogeneous single-cell RNA-seq dataset collections. Nature Methods 16(8), 695–698 (2019)

111. Korsunsky, I., Millard, N., Fan, J., Slowikowski, K., Zhang, F., Wei, K., Baglaenko, Y., Brenner, M., Loh, P.-r., Raychaudhuri, S.: Fast, sensitive and accurate integration of single-cell data with Harmony. Nature Methods 16(12), 1289–1296 (2019)

112. Haghverdi, L., Lun, A.T., Morgan, M.D., Marioni, J.C.: Batch effects in single-cell RNA-sequencing data are corrected by matching mutual nearest neighbors. Nature Biotechnology 36(5), 421–427 (2018)

113. Parker, H.S., Leek, J.T., Favorov, A.V., Considine, M., Xia, X., Chavan, S., Chung, C.H., Fertig, E.J.: Preserving biological heterogeneity with a permuted surrogate variable analysis for genomics batch correction. Bioinformatics 30(19), 2757–2763 (2014)

114. Tran, H.T.N., Ang, K.S., Chevrier, M., Zhang, X., Lee, N.Y.S., Goh, M., Chen, J.: A benchmark of batch-effect correction methods for single-cell RNA sequencing data. Genome Biology 21, 12 (2020)

115. Richards, L.M., Riverin, M., Mohanraj, S., Ayyadhury, S., Croucher, D.C., Díaz-Mejía, J.J., Coutinho, F.J., Dirks, P.B., Pugh, T.J.: A comparison of data integration methods for single-cell RNA sequencing of cancer samples. bioRxiv (2021). doi:10.1101/2021.08.04.453579

116. Fan, J., Slowikowski, K., Zhang, F.: Single-cell transcriptomics in cancer: computational challenges and opportunities. Experimental and Molecular Medicine 52(9), 1452–1465 (2020)

117. Gouvert, O., Oberlin, T., Févotte, C.: Negative binomial matrix factorization for recommender systems. arXiv 1801.01708 (2018)

118. Gu, J., Wang, X., Halakivi-Clarke, L., Clarke, R., Xuan, J.: BADGE: A novel Bayesian model for accurate abundance quantification and differential analysis of RNA-Seq data. BMC Bioinformatics 15(S9), 6 (2014)

119. Wang, C., Sun, D., Huang, X., Wan, C., Li, Z., Han, Y., Qin, Q., Fan, J., Qiu, X., Xie, Y., Meyer, C.A., Brown, M., Tang, M., Long, H., Liu, T., Liu, X.S.: Integrative analyses of single-cell transcriptome and regulome using MAESTRO. Genome Biology 21, 198 (2020)

120. Lareau, C.A., Duarte, F.M., Chew, J.G., Kartha, V.K., Burkett, Z.D., Kohlway, A.S., Pokholok, D., Aryee, M.J., Steemers, F.J., Lebofsky, R., Buenrostro, J.D.: Droplet-based combinatorial indexing for massive-scale single-cell chromatin accessibility. Nature Biotechnology 37(8), 916–924 (2019)

121. Kartha, V.K., Duarte, F.M., Hu, Y., Ma, S., Chew, J.G., Lareau, C.A., Earl, A., Burkett, Z.D., Kohlway, A.S., Lebofsky, R., Buenrostro, J.D.: Functional inference of gene regulation using single-cell multi-omics. Cell Genomics 2(9), 100166 (2022)

122. Bravo González-Blas, C., De Winter, S., Hulselmans, G., Hecker, N., Matetovici, I., Christiaens, V., Poovathingal, S., Wouters, J., Aibar, S., Aerts, S.: SCENIC+: single-cell multiomic inference of enhancers and gene regulatory networks. Nature Methods 20(9), 1355–1367 (2023)

123. Ma, S., Zhang, B., LaFave, L.M., Earl, A.S., Chiang, Z., Hu, Y., Ding, J., Brack, A., Kartha, V.K., Tay, T., Law, T., Lareau, C., Hsu, Y.-C., Regev, A., Buenrostro, J.D.: Chromatin potential identified by shared single-cell profiling of RNA and chromatin. Cell 183(4), 1103–111620 (2020)

124. Cao, J., Cusanovich, D.A., Ramani, V., Aghamirzaie, D., Pliner, H.A., Hill, A.J., Daza, R.M., McFaline-Figueroa, J.L., Packer, J.S., Christiansen, L., Steemers, F.J., Adey, A.C., Trapnell, C., Shendure, J.: Joint profiling of chromatin accessibility and gene expression in thousands of single cells. Science 361(6409), 1380–1385 (2018)

125. Stuart, T., Satija, R.: Integrative single-cell analysis. Nature Reviews Genetics 20(5), 257–272 (2019)

126. Shiga, M., Seno, S., Onizuka, M., Matsuda, H.: Sc-jnmf: single-cell clustering integrating multiple quantification methods based on joint non-negative matrix factorization. PeerJ 9, 12087 (2021)

127. Zhang, S., Liu, C.-C., Li, W., Shen, H., Laird, P.W., Zhou, X.J.: Discovery of multi-dimensional modules by integrative analysis of cancer genomic data. Nucleic Acids Research 40(19), 9379–9391 (2012)

128. Yang, Z., Michailidis, G.: A non-negative matrix factorization method for detecting modules in heterogeneous omics multi-modal data. Bioinformatics 32(1), 1–8 (2015)

129. Jin, S., Zhang, L., Nie, Q.: scAI: an unsupervised approach for the integrative analysis of parallel single-cell transcriptomic and epigenomic profiles. Genome Biology 21, 25 (2020)

130. Hao, Y., Stuart, T., Kowalski, M.H., Choudhary, S., Hoffman, P., Hartman, A., Srivastava, A., Molla, G., Madad, S., Fernandez-Granda, C., Satija, R.: Dictionary learning for integrative, multimodal and scalable single-cell analysis. Nature Biotechnology (2023). doi:10.1038/s41587-023-01767-y

131. Argelaguet, R., Velten, B., Arnol, D., Dietrich, S., Zenz, T., Marioni, J.C., Buettner, F., Huber, W., Stegle, O.: Multi-omics factor analysis—a framework for unsupervised integration of multi-omics data sets. Molecular Systems Biology 14, 8124 (2018)

132. Argelaguet, R., Arnol, D., Bredikhin, D., Deloro, Y., Velten, B., Marioni, J.C., Stegle, O.: MOFA+: a statistical framework for comprehensive integration of multi-modal single-cell data. Genome Biology 21, 111 (2020)

133. Baker, S.M., Rogerson, C., Hayes, A., Sharrocks, A.D., Rattray, M.: Classifying cells with Scasat, a single-cell ATAC-seq analysis tool. Nucleic Acids Research 47(2), 10 (2019)

134. Nair, S., Kim, D.S., Perricone, J., Kundaje, A.: Integrating regulatory DNA sequence and gene expression to predict genome-wide chromatin accessibility across cellular contexts. Bioinformatics 35, 108–116 (2019)

135. Pott, S., Lieb, J.D.: Single-cell ATAC-seq: strength in numbers. Genome Biology 16, 172 (2015)

136. Yan, F., Powell, D.R., Curtis, D.J., Wong, N.C.: From reads to insight: a hitchhiker’s guide to ATAC-seq data analysis. Genome Biology 21, 22 (2020)

137. Taddy, M.: Distributed multinomial regression. Annals of Applied Statistics 9(3), 1394–1414 (2015)

138. Fisher, R.A.: On the interpretation of χ^2^ from contingency tables, and the calculation of P. Journal of the Royal Statistical Society 85(1), 87–94 (1922)

139. Good, I.J.: Some statistical applications of Poisson’s work. Statistical Science 1(2), 157–170 (1986)

140. Gelman, A., Carlin, J.B., Stern, H.S., Dunson, D.B., Vehtari, A., Rubin, D.B.: Bayesian Data Analysis, 3rd edn. CRC Press, Boca Raton, FL (2013)

141. Andrieu, C., de Freitas, N., Doucet, A., Jordan, M.I.: An introduction to MCMC for machine learning. Machine Learning 50, 5–43 (2003)

142. Robert, C.P.: Monte Carlo Statistical Methods, 2nd edn. Springer, New York, NY (2004)

143. Devroye, L.: Non-uniform Random Variate Generation. Springer, New York, NY (1986)

144. Stephens, M., Carbonetto, P., Gerard, D., Lu, M., Sun, L., Willwerscheid, J., Xiao, N.: ashr: methods for adaptive shrinkage, using empirical Bayes. R package version 2.2-51 (2020). https://github.com/stephens999/ashr

145. Chen, M.-H., Shao, Q.-M.: Monte Carlo estimation of Bayesian credible and HPD intervals. Journal of Computational and Graphical Statistics 8(1), 69–92 (1999)

146. Box, G.E.P., Tiao, G.C.: Bayesian Inference in Statistical Analysis. Addison-Wesley, Reading, MA (1992)

147. Gelman, A., Hill, J., Yajima, M.: Why we (usually) don’t have to worry about multiple comparisons. Journal of Research on Educational Effectiveness 5(2), 189–211 (2012)

148. Zheng, G.X.Y., Terry, J.M., Belgrader, P., Ryvkin, P., Bent, Z.W., Wilson, R., Ziraldo, S.B., Wheeler, T.D., McDermott, G.P., Zhu, J., Gregory, M.T., Shuga, J., Montesclaros, L., Underwood, J.G., Masquelier, D.A., Nishimura, S.Y., Schnall-Levin, M., Wyatt, P.W., Hindson, C.M., Bharadwaj, R., Wong, A., Ness, K.D., Beppu, L.W., Deeg, H.J., McFarland, C., Loeb, K.R., Valente, W.J., Ericson, N.G., Stevens, E.A., Radich, J.P., Mikkelsen, T.S., Hindson, B.J., Bielas, J.H.: Massively parallel digital transcriptional profiling of single cells. Data sets. 10x Genomics (2017). https://www.10xgenomics.com/support/single-cell-gene-expression

149. Montoro, D.T., Haber, A.L., Biton, M., Vinarsky, V., Chen, S., Villoria, J., Rogel, N., Tata, P.R., Rowe, S.M., Engelhardt, J.F., Regev, A., Rajagopal, J., Lin, B., Birket, S., Yuan, F., Leung, H., Burgin, G., Tsankov, A., Waghray, A., Slyper, M., Waldman, J., Nguyen, D. L. Dionne Rozenblatt-Rosen, O., Mou, H., Shivaraju, M., Bihler, H., Mense, M., Tearney, G.: A revised airway epithelial hierarchy includes CFTR-expressing ionocytes. Data sets. Gene Expression Omnibus (2018). https://www.ncbi.nlm.nih.gov/geo/query/acc.cgi?acc=GSE103354

150. Cusanovich, D.A., Daza, R., Adey, A., Pliner, H.A., Christiansen, L., Gunderson, K.L., Steemers, F.J., Trapnell, C., Shendure, J.: Multiplex single-cell profiling of chromatin accessibility by combinatorial cellular indexing. Science 348(6237), 910–914 (2015)

151. Cusanovich, D.A., Reddington, J.P., Garfield, D.A., Daza, R.M., Aghamirzaie, D., Marco-Ferreres, R., Pliner, H.A., Christiansen, L., Qiu, X., Steemers, F.J., Trapnell, C., Shendure, J., Furlong, E.E.M.: The cis-regulatory dynamics of embryonic development at single-cell resolution. Nature 555(7697), 538–542 (2018)

152. Cusanovich, D.A., Hill, A.J., Aghamirzaie, D., Daza, R.M., Pliner, H.A., Berletch, J.B., Filippova, G.N., Huang, X., Christiansen, L., DeWitt, W.S., Lee, C., Regalado, S.G., Read, D.F., Steemers, F.J., Disteche, C.M., Trapnell, C., Shendure, J.: A single-cell atlas of in vivo mammalian chromatin accessibility. Data sets. Mouse sci-ATAC-seq Atlas (2018). https://shendurelab.github.io/mouse-atac/

153. Buenrostro, J.D., Corces, M.R., Lareau, C.A., Wu, B., Schep, A.N., Aryee, M.J., Majeti, R., Chang, H.Y., Greenleaf, W.J.: Single-cell epigenomics maps the continuous regulatory landscape of human hematopoietic differentiation. Data sets. Gene Expression Omnibus (2018). https://www.ncbi.nlm.nih.gov/geo/query/acc.cgi?acc=GSE96772

154. Chen, H., Lareau, C., Andreani, T., Vinyard, M.E., Garcia, S.P., Clement, K., Andrade-Navarro, M.A., Buenrostro, J.D., Pinello, L.: Assessment of computational methods for the analysis of single-cell ATAC-seq data. Genome Biology 20, 241 (2019)

155. Hien, L.T.K., Gillis, N.: Algorithms for nonnegative matrix factorization with the Kullback–Leibler divergence. Journal of Scientific Computing 87(3), 93 (2021)

156. Hsieh, C.-J., Dhillon, I.S.: Fast coordinate descent methods with variable selection for non-negative matrix factorization. In: Proceedings of the 17th ACM SIGKDD International Conference, pp. 1064–1072 (2011)

157. Lin, X., Boutros, P.C.: Optimization and expansion of non-negative matrix factorization. BMC Bioinformatics 21, 7 (2020)

158. Ang, A.M.S., Gillis, N.: Accelerating nonnegative matrix factorization algorithms using extrapolation. Neural Computation 31(2), 417–439 (2019)

159. Ke, Z.T., Wang, M.: A new SVD approach to optimal topic estimation. arXiv 1704.07016 (2019)

160. van der Maaten, L.: Accelerating t-SNE using tree-based algorithms. Journal of Machine Learning Research 15(93), 3221–3245 (2014)

161. Krijthe, J.H.: Rtsne: t-distributed stochastic neighbor embedding using Barnes-Hut implementation. R package version 0.15 (2015). https://github.com/jkrijthe/Rtsne

162. Ding, C., He, X.: K-means clustering via principal component analysis. In: 21st International Conference on Machine Learning (2004)

163. Kass, R.E., Raftery, A.E.: Bayes factors. Journal of the American Statistical Association 90(430), 773–795 (1995)

164. Geer, L.Y., Marchler-Bauer, A., Geer, R.C., Han, L., He, J., He, S., Liu, C., Shi, W., Bryant, S.H.: The NCBI biosystems database. Nucleic Acids Research 38(upplement-1), 492–496 (2009)

165. Cerami, E.G., Gross, B.E., Demir, E., Rodchenkov, I., Babur, Ö ., Anwar, N., Schultz, N., Bader, G.D., Sander, C.: Pathway Commons, a web resource for biological pathway data. Nucleic Acids Research 39(upplement 1), 685–690 (2010)

166. Rodchenkov, I., Babur, Ö ., Luna, A., Aksoy, B.A., Wong, J.V., Fong, D., Franz, M., Siper, M.C., Cheung, M., Wrana, M., Mistry, H., Mosier, L., Dlin, J., Wen, Q., O’Callaghan, C., Li, W., Elder, G., Smith, P.T., Dallago, C., Cerami, E., Gross, B., Dogrusoz, U., Demir, E., Bader, G.D., Sander, C.: Pathway Commons 2019 update: integration, analysis and exploration of pathway data. Nucleic Acids Research 48(D1), 489–497 (2019)

167. Subramanian, A., Tamayo, P., Mootha, V.K., Mukherjee, S., Ebert, B.L., Gillette, M.A., Paulovich, A., Pomeroy, S.L., Golub, T.R., Lander, E.S., Mesirov, J.P.: Gene set enrichment analysis: a knowledge-based approach for interpreting genome-wide expression profiles. Proceedings of the National Academy of Sciences 102(43), 15545–15550 (2005)

168. Liberzon, A., Birger, C., Thorvaldsdóttir, H., Ghandi, M., Mesirov, J.P., Tamayo, P.: The molecular signatures database hallmark gene set collection. Cell Systems 1(6), 417–425 (2015)

169. Liberzon, A., Subramanian, A., Pinchback, R., Thorvaldsdottir, H., Tamayo, P., Mesirov, J.P.: Molecular signatures database (MSigDB) 3.0. Bioinformatics 27(12), 1739–1740 (2011)

170. Ashburner, M., Ball, C.A., Blake, J.A., Botstein, D., Butler, H., Cherry, J.M., Davis, A.P., Dolinski, K., Dwight, S.S., Eppig, J.T., Harris, M.A., Hill, D.P., Issel-Tarver, L., Kasarskis, A., Lewis, S., Matese, J.C., Richardson, J.E., Ringwald, M., Rubin, G.M., Sherlock, G.: Gene Ontology: tool for the unification of biology. Nature Genetics 25(1), 25–29 (2000)

171. Carbonetto, P., Stephens, M.: pathways: gene set enrichment analysis using human and mouse gene sets. R package version 0.1-20 (2021). https://github.com/stephenslab/pathways

172. Fang, T., Davydov, I., Marbach, D., Zhang, J.D.: Gene-set enrichment with regularized regression. bioRxiv (2019). doi:10.1101/659920

173. Sartor, M.A., Leikauf, G.D., Medvedovic, M.: LRpath: A logistic regression approach for identifying enriched biological groups in gene expression data. Bioinformatics 25(2), 211–217 (2009)

174. Harrison, P.F., Pattison, A.D., Powell, D.R., Beilharz, T.H.: Topconfects: a package for confident effect sizes in differential expression analysis provides a more biologically useful ranked gene list. Genome Biology 20, 67 (2019)

175. McCarthy, D.J., Smyth, G.K.: Testing significance relative to a fold-change threshold is a TREAT. Bioinformatics 25(6), 765–771 (2009)

176. Wang, G., Sarkar, A., Carbonetto, P., Stephens, M.: A simple new approach to variable selection in regression, with application to genetic fine mapping. Journal of the Royal Statistical Society, Series B 82(5), 1273–1300 (2020)

177. Bauer, S., Gagneur, J., Robinson, P.N.: GOing Bayesian: model-based gene set analysis of genome-scale data. Nucleic Acids Research 38(11), 3523–3532 (2020)

178. Ebrahimpoor, M., Spitali, P., Hettne, K., Tsonaka, R., Goeman, J.: Simultaneous enrichment analysis of all possible gene-sets: unifying self-contained and competitive methods. Briefings in Bioinformatics 21(4), 1302–1312 (2019)

179. Fontanillo, C., Nogales-Cadenas, R., Pascual-Montano, A., Rivas, J.D.L.: Functional analysis beyond enrichment: non-redundant reciprocal linkage of genes and biological terms. PLoS ONE 6(9), 24289 (2011)

180. Lu, Y., Rosenfeld, R., Simon, I., Nau, G.J., Bar-Joseph, Z.: A probabilistic generative model for GO enrichment analysis. Nucleic Acids Research 36(17), 109 (2008)

181. Simillion, C., Liechti, R., Lischer, H.E.L., Ioannidis, V., Bruggmann, R.: Avoiding the pitfalls of gene set enrichment analysis with SetRank. BMC Bioinformatics 18, 151 (2017)

182. Vivar, J.C., Pemu, P., McPherson, R., Ghosh, S.: Redundancy control in pathway databases (ReCiPa): an application for improving gene-set enrichment analysis in omics studies and “big data” biology. Omics 17(8), 414–422 (2013)

183. McDavid, A., Finak, G., Yajima, M.: MAST: model-based analysis of single cell transcriptomics. R package version 1.20.0 (2021). https://github.com/RGLab/MAST

184. Carbonetto, P., Luo, K., Dey, K., Hsiao, J., Sarkar, A., Hung, A., Stephens, M.: fastTopics: fast algorithms for fitting topic models and non-negative matrix factorizations to count data. R package version 0.6-142 (2022). https://cran.r-project.org/package=fastTopics

185. R Core Team: R: a language and environment for statistical computing, Vienna, Austria. R Foundation for Statistical Computing (2018). https://www.R-project.org

186. Carbonetto, P., Lao, K., Sarkar, A., Hung, A., Tayeb, K., Pott, S., Stephens, M.: Analysis of single-cell RNA-seq data sets for this manuscript. Zenodo (2023). doi:10.5281/zenodo.7962782

187. Carbonetto, P., Lao, K., Sarkar, A., Hung, A., Tayeb, K., Pott, S., Stephens, M.: Analysis of single-cell ATAC-seq data sets for this manuscript. Zenodo (2023). doi:10.5281/zenodo.7962831

188. Blischak, J.D., Carbonetto, P., Stephens, M.: Creating and sharing reproducible research code the workflowr way. F1000Research 8, 1749 (2019)

