## Additional file 1: Supplementary Figures S1-S16 for "GoM DE: interpreting structure in sequence count data with differential expression analysis allowing for grades of membership"

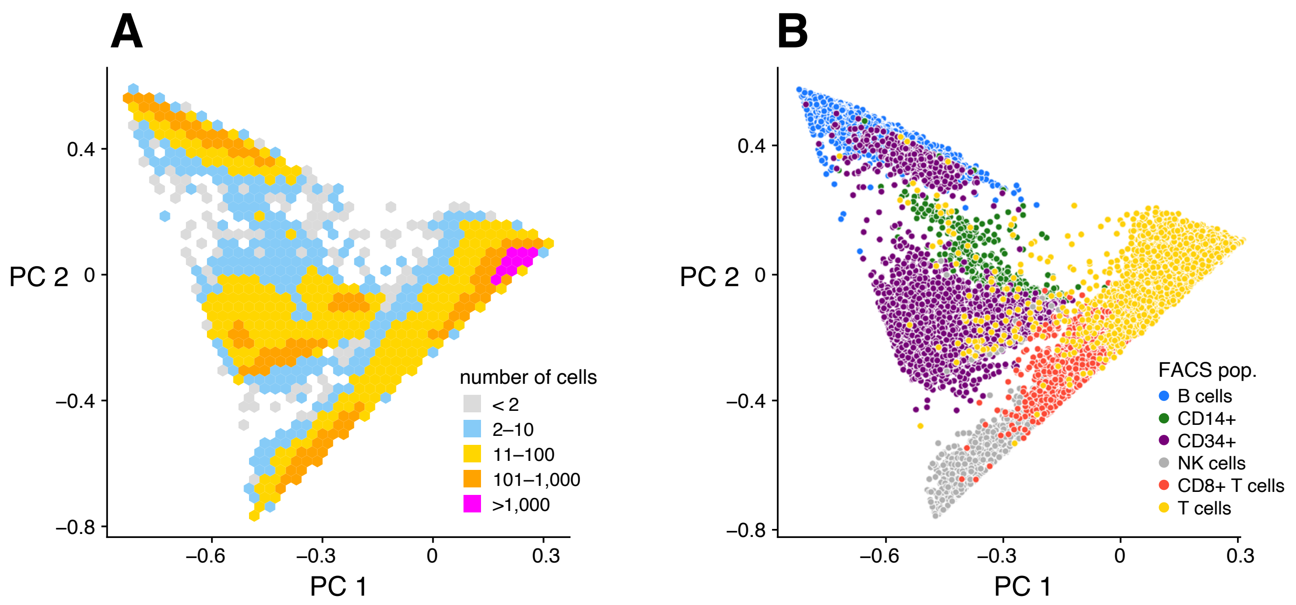


**Figure S1.** Cells plotted by the first two PCs of the estimated **L**. Panel A shows the density of the cells in this projection. In Panel B, cells are colored by FACS-labeled subpopulation. The “T cells” label includes all T cell FACS subpopulations except CD8+ cytotoxic T cells.

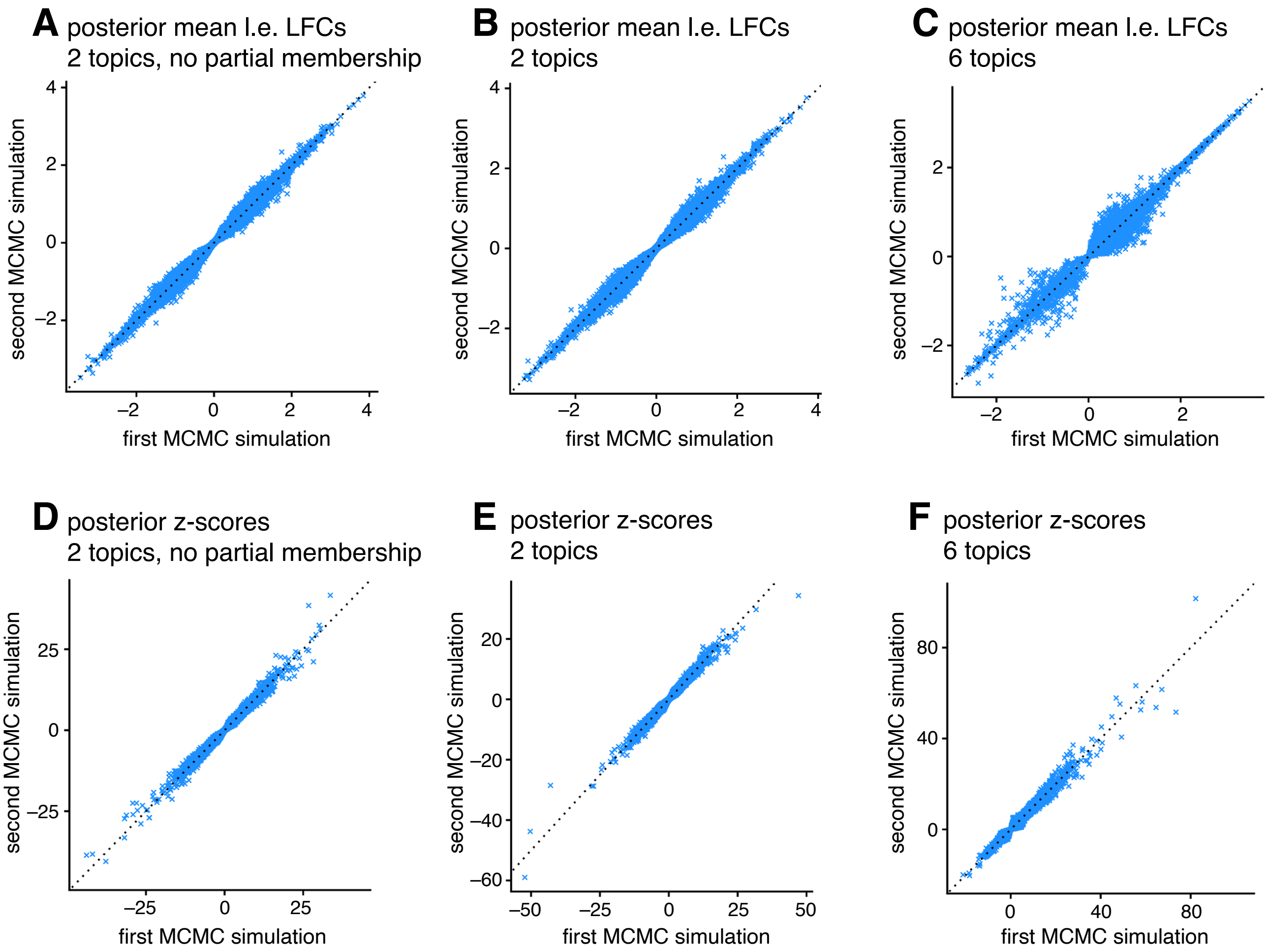


**Figure S2.** Assessment of posterior computations for GoM DE analysis. of the posterior calculations, we performed the MCMC twice, and compared the posterior means and posterior *z*-scores post-adaptive shrinkage. If the posterior mean LFCs and posterior *z*-scores are consistent between the two runs, this is a good indication that they are accurate. Panels A, B and C compare MCMC estimates of posterior mean l.e. LFCs. Panels D, E and F compare MCMC estimates of posterior *z*-scores. In each scatterplot, 200,000 points are plotted (10,000 genes $\times$ 20 simulated data sets).

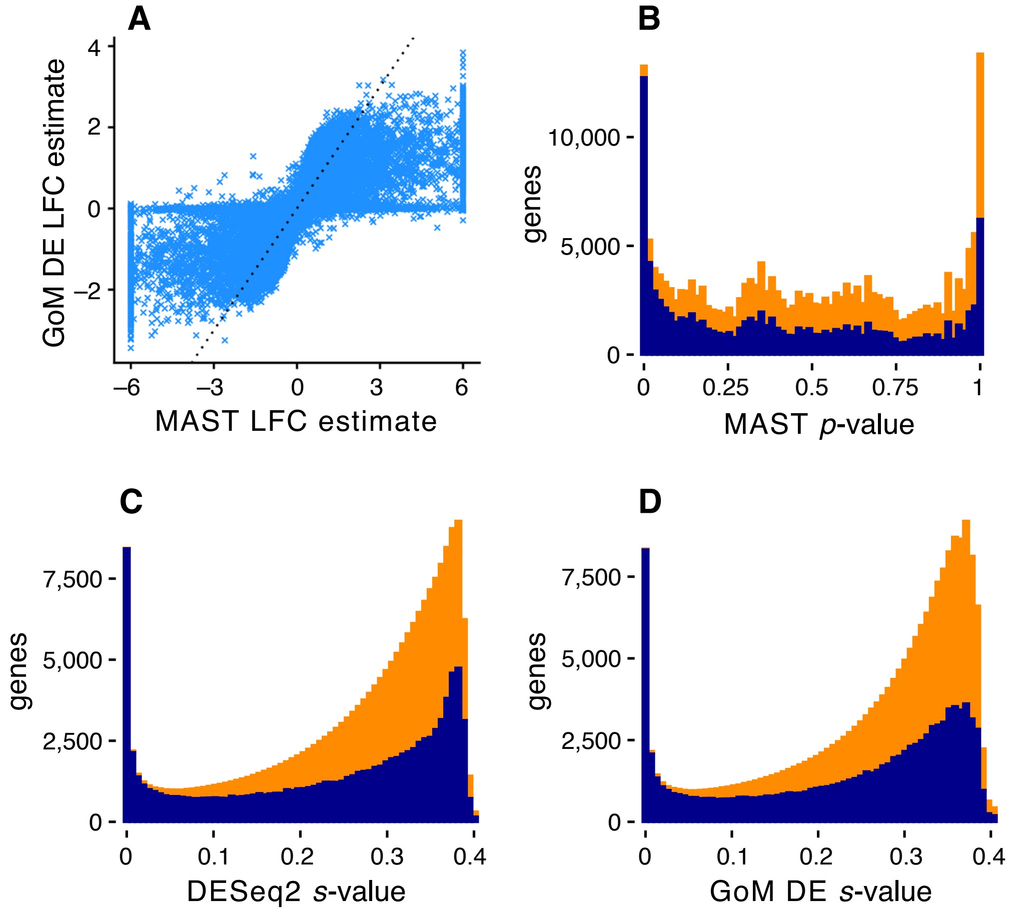


**Figure S3.** Additional evaluation of DE analysis methods in simulated data sets in which cells are simulated *without* partial membership to topics. Panel A compares LFC estimates returned by MAST and the GoM DE analysis. In this plot, 200,000 points are plotted (10,000 genes $\times$ 20 simulated data sets). Panels B, C and D compare the distribution of *p*-values (MAST) and *s*-values (DESeq2, GoM DE), separately for true differences (dark blue) and non-differences (orange).

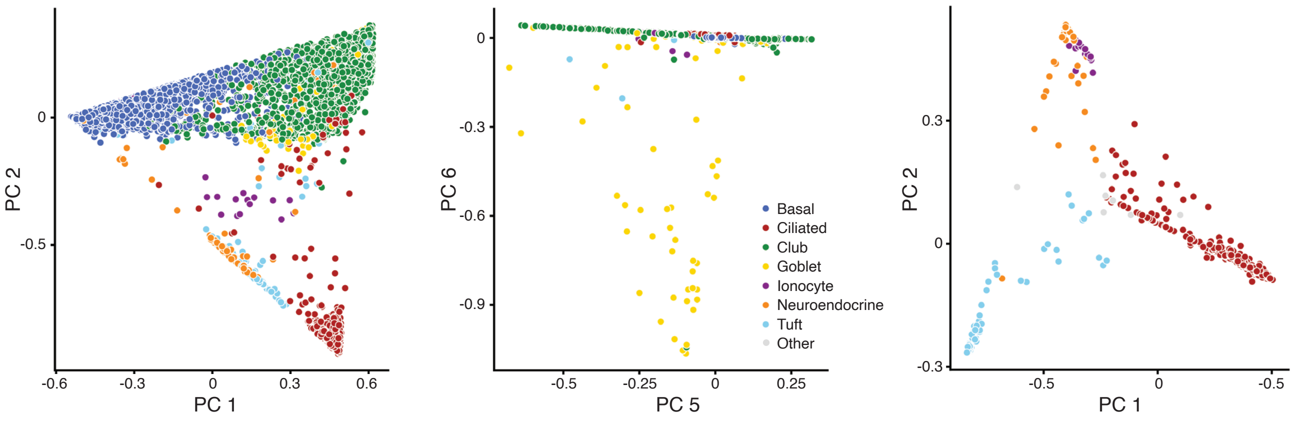


**Figure S4.** These plots show the 7,193 cells (left, middle) and 637 cells (right) projected onto PCs of the estimated **L**. The color of the points is varied according to the clustering obtained in Montoro et al (2018) using a community detection algorithm. *Left, middle:* Correspondence between the $K$ = 7 topic model fit to epithelial airway data and the clustering from Montoro et al (2018). *Right:* Correspondence between the $K$ = 5 topic model fit to the subset of rare epithelial cell types and the clustering from Montoro et al (2018). The “Other” label in the right-hand plot is for the small number cells included in the $n$ = 637 subset of rare epithelial cell types that were labeled by Montoro et al (2018) as belonging the basal, club and goblet clusters.

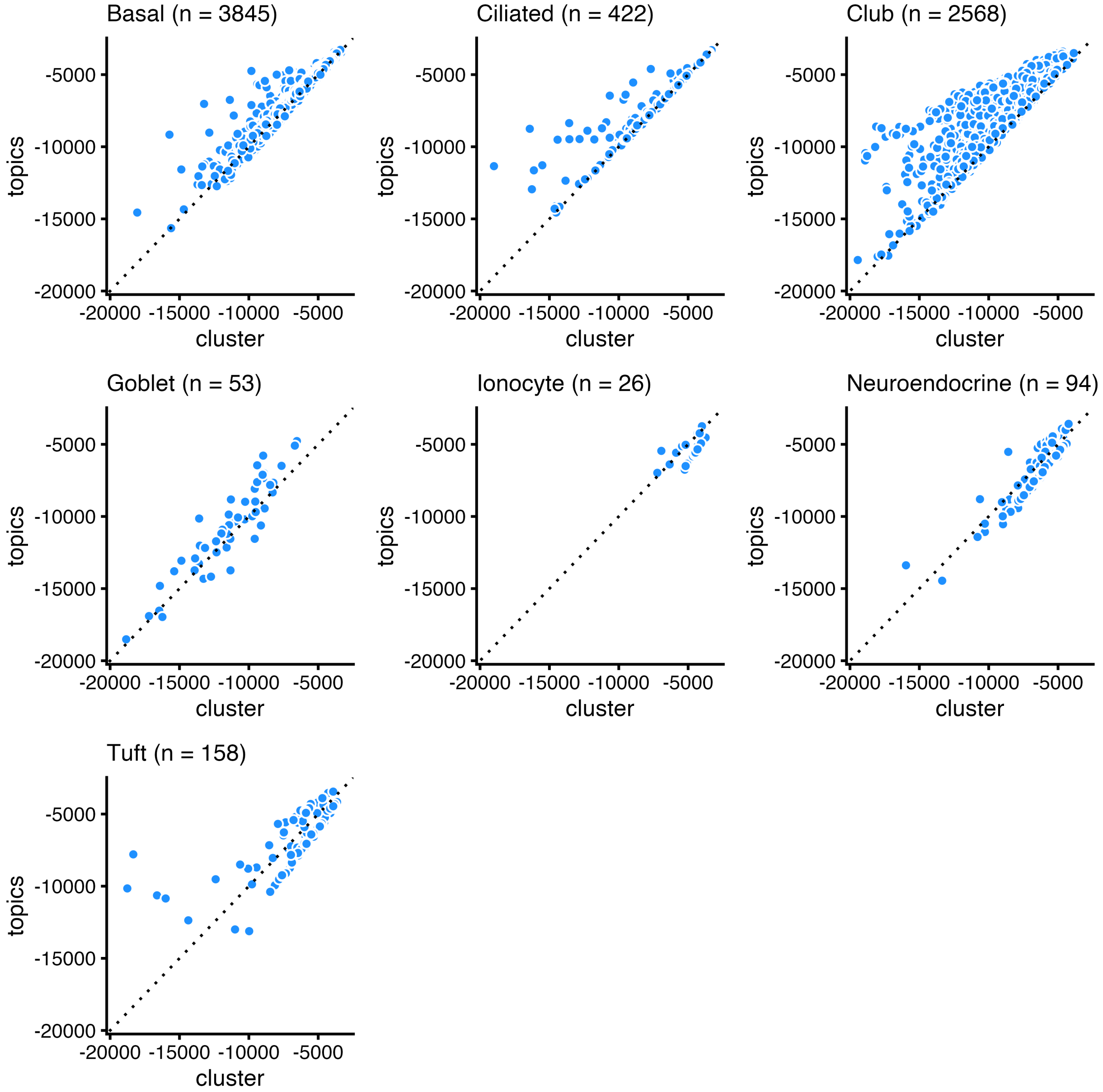


**Figure S5.** Assessment of single-cell likelihoods for $K$ = 7 topic model fit to epithelial airway data. In Montoro et al (2018), the 7,193 cells were subdivided into 7 clusters corresponding to 7 cell types (basal, ciliated, club, goblet, ionocyte, neuroendocrine, tuft). Here we assess how well the topic model captures expression in these cell types by comparing topic model likelihoods against likelihoods under a simple multinomial model (eq. 1 in main text) in which all cells share the same underlying pattern of expression; that is, $\pi_{ij} = \pi_j$ for all cells $i$ in the cluster. (The multinomial probabilities $\pi_j$ in this simple model are estimated by maximum-likelihood; $\hat{\pi}_j = \sum_i x_{ij} / \sum_i s_i$, in which the sums are over all cells $i$ assigned to the cluster.) The scatterplots compare, for each cell, the log-likelihood under this simple cluster-based multinomial model (x-axis) against the topic model log-likelihood (y-axis). The topic model provides a better fit for cells above the diagonal (dotted line) and a worse fit for cells below the diagonal.

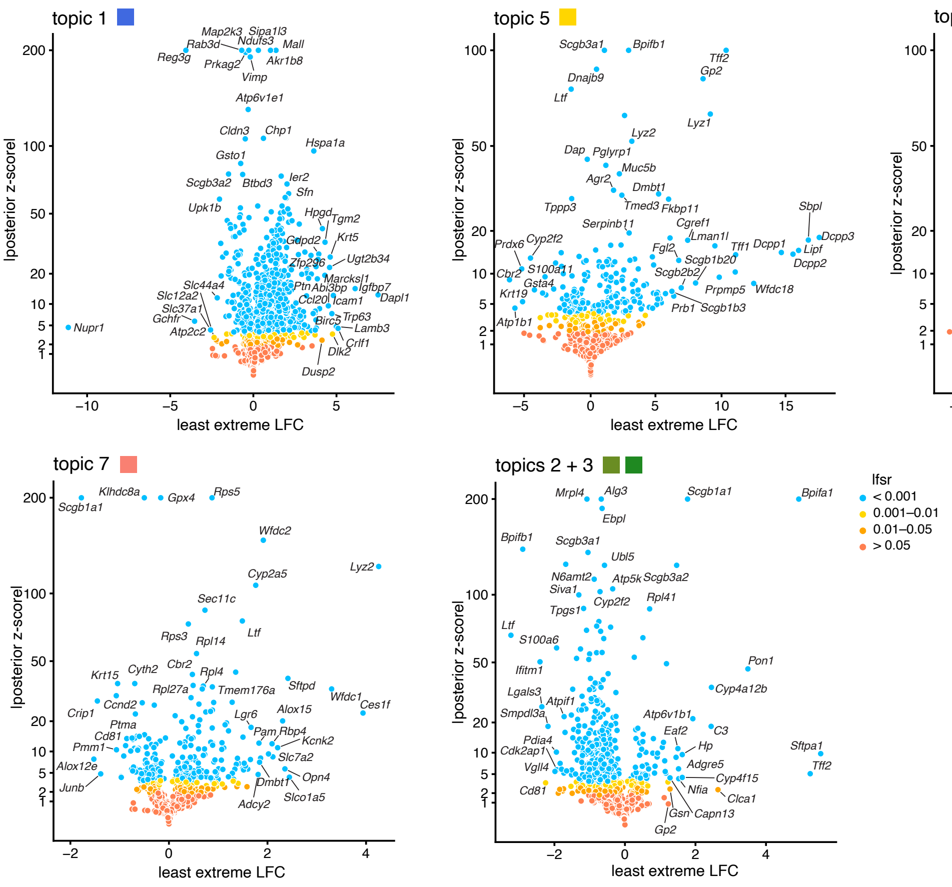


**Figure S6.** GoM DE analysis of additional topics in $K = 7$ topic model fitted to epithelial airway data. The topic colors correspond to colors used in the Structure plot in Figure 7A. The volcano plots show moderated posterior estimates of l.e. LFC vs. posterior $z$-scores for 18,388 genes. A small number of genes with extreme posterior $z$-scores are shown with smaller posterior $z$-scores so that they fit within the y-axis range; actual posterior $z$-score statistics and other detailed statistics can be accessed in the interactive volcano plots and supplementary tables. For the GoM DE analysis of the combined topic, the membership proportions for the combined topic were defined as $l'_{i,2+3} = l_{i2} + l_{i3}$, then the GoM DE analysis proceeded using a modified membership matrix $\mathbf{L}' \in \mathbf{R}^{18,388 \times 6}$, in which $l'_{ik} = l_{ik}$ for all $i, k = 1, 4$–$7$.

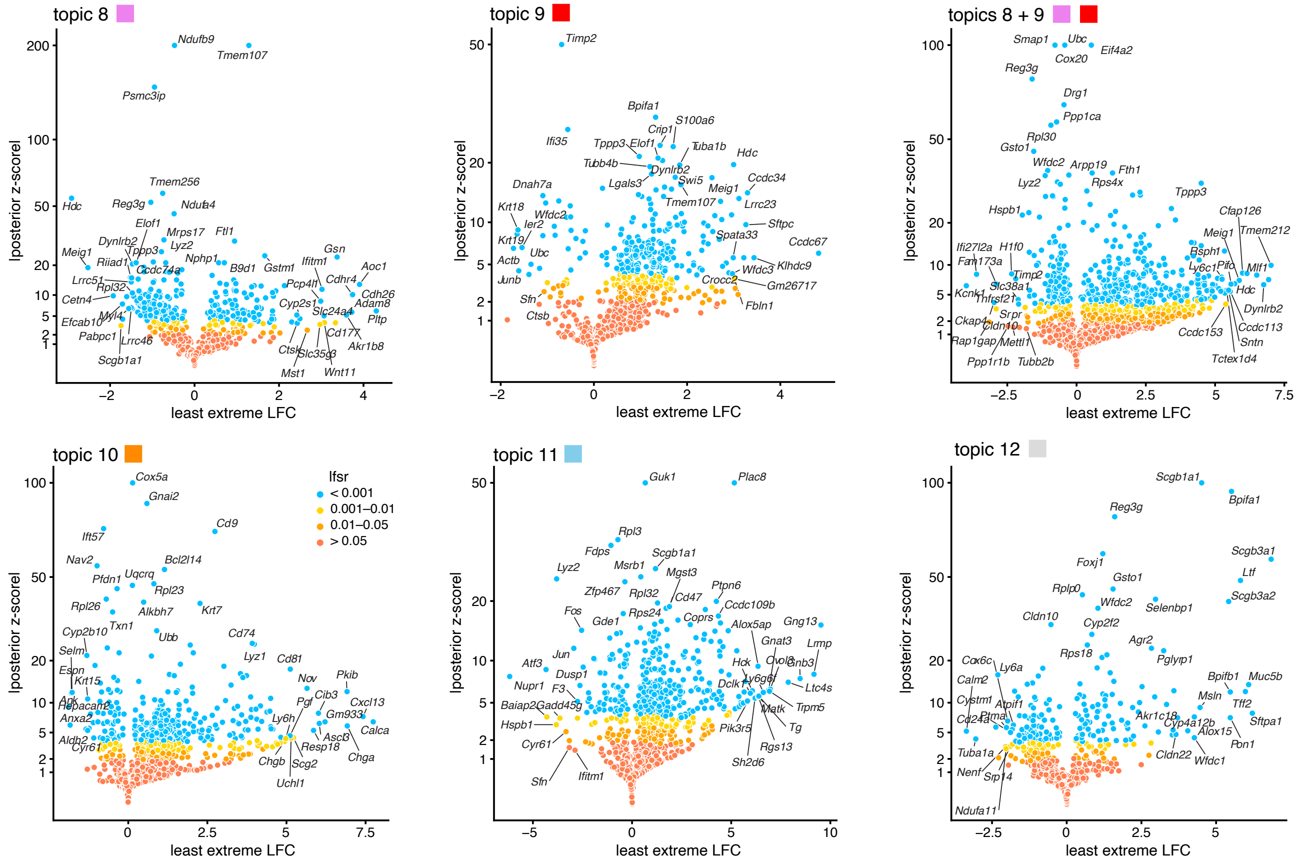


**Figure S7.** GoM DE analysis of topics in the $K = 5$ topic model fitted to the rare epithelial cell type subset. The topic colors correspond to colors used in the Structure plot in Figure 7A. The volcano plots show moderated posterior estimates of l.e. LFC vs. posterior $z$-scores for 15,594 genes. A small number of genes with extreme posterior $z$-scores are shown with smaller posterior $z$-scores so that they fit within the Y-axis range; actual posterior $z$-score statistics and other detailed statistics can be accessed in the interactive volcano plots and supplementary tables. For the GoM DE analysis of the combined topic, the membership proportions for the combined topic were defined as $l'_{i,8+9} = l_{i8} + l_{i9}$, then the GoM DE analysis proceeded using a modified membership matrix $\mathbf{L}' \in \mathbf{R}^{15{,}594 \times 5}$, in which $l'_{ik} = l_{ik}$ for all $i, k = 10, 11, 12$.

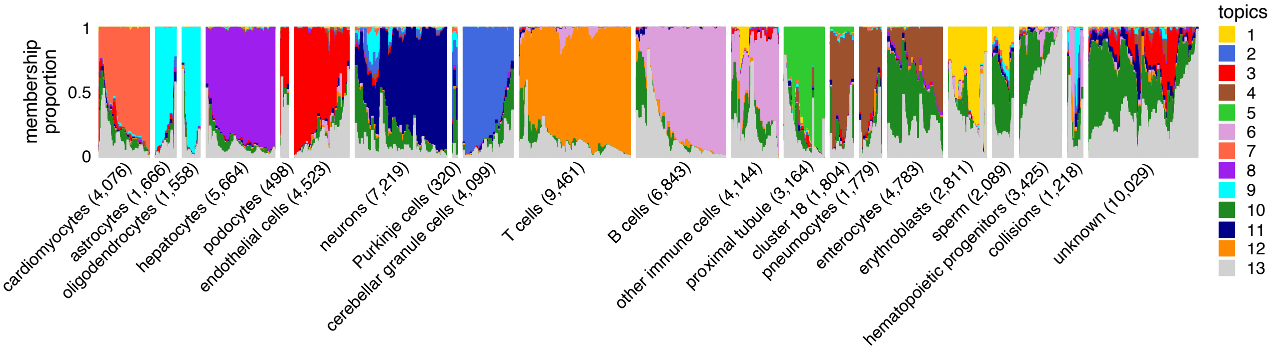


**Figure S8.** Structure plot visualizing topic model fit to the Mouse Atlas sci-ATAC-seq data with $K = 13$ topics. The cells are grouped by "cell type"—specifically the "cell_label" column of the metadata table which gives final cell-type assignments incorporating automated and manual assignments; see Table S1 in Cusanovich et al (2018) for details. "Other immune cells" includes natural killer cells, monocytes, macrophages, microglia and dendritic cells. "Cluster 18" includes the cells—mostly kidney cells—labeled as "collecting duct", "loop of henle" and "DCT/CD". The numbers of parentheses next to each cell type label give the number of cells assigned to that cell type. Compare to Figure 8A.

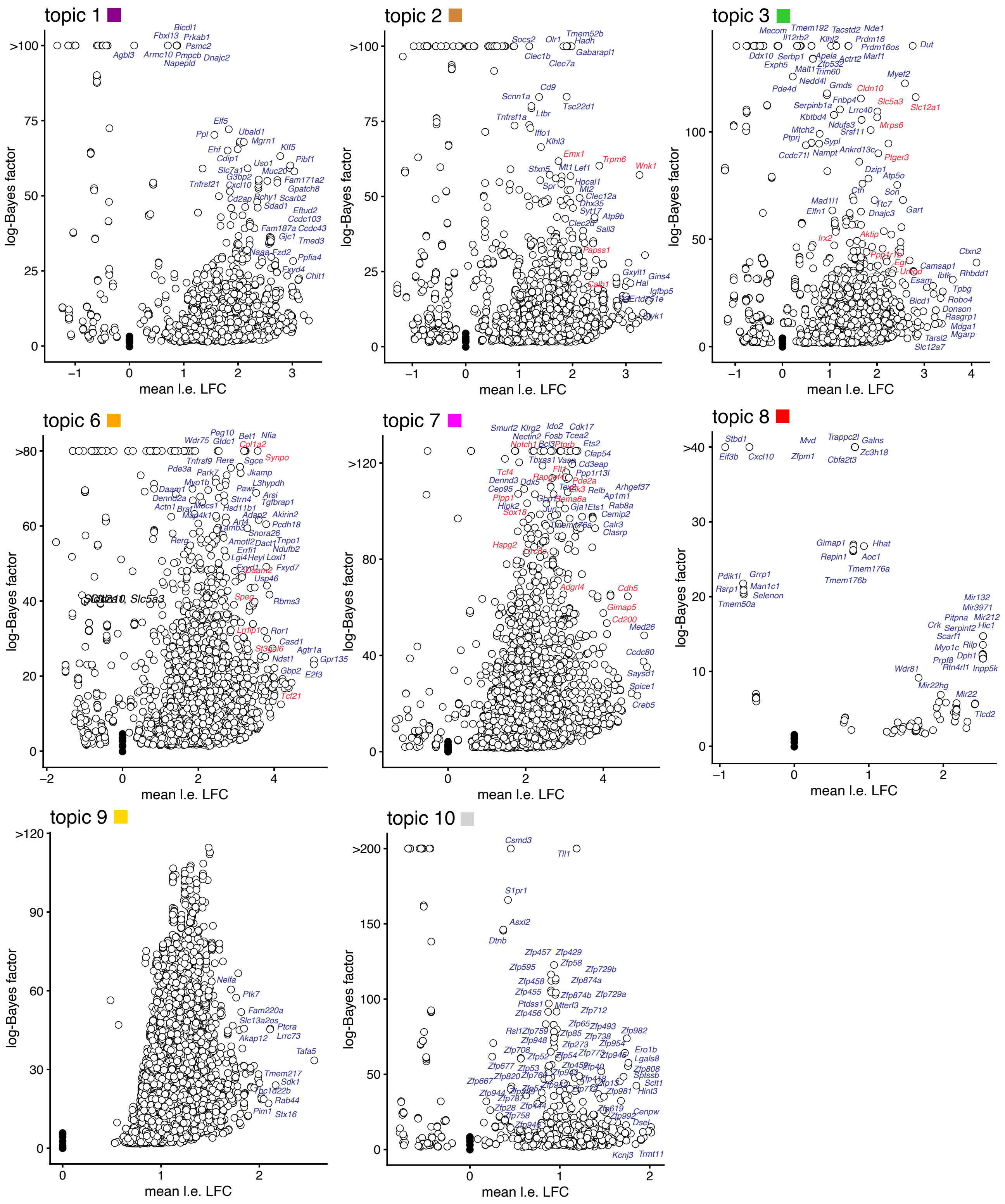


**Figure S9.** Additional gene enrichment results for topics in Mouse sci-ATAC-seq Atlas kidney data based on differences in accessibility of peaks linked to genes by Cicero. “Mean l.e. LFC” is the average l.e. LFC among all peaks connected to the gene, restricted to peaks that are differentially accessible (*lfsr* $< 0.05$). Note that some genes at the top of the volcano plot may actually have a larger log-Bayes factor than what is shown; the log-Bayes factors were truncated to fit within the range of the volcano plot. Marker genes with cell-type-specific expression in distal convoluted tubule (topic 2), loop of Henle (topic 3), podocytes (topic 6) and endothelial cells (topic 7) are highlighted in red. These marker genes were identified from a differential expression analysis of single-cell RNA-seq profiles in healthy kidney cells (see Table S1 of Park et al 2018 for the full list of identified marker genes).

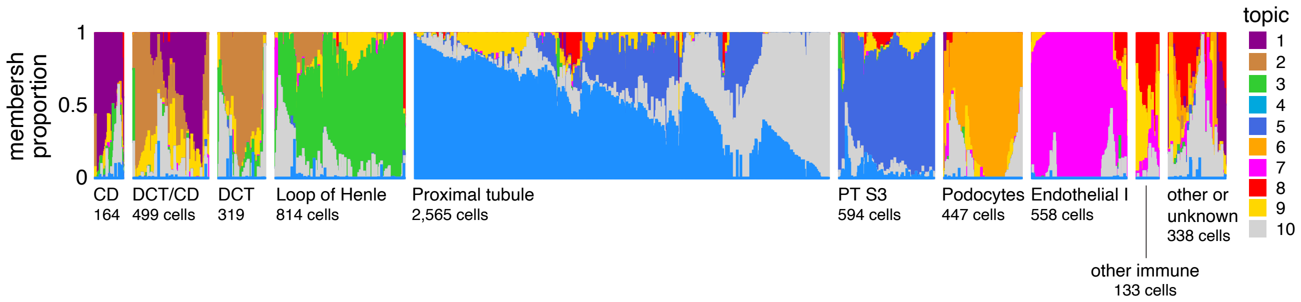


**Figure S10.** Structure plots visualizing the topic model fit to the Mouse Atlas kidney cells, with $K = 10$ topics, in which the cells are arranged by the cell-type assignments estimated in Cusanovich et al (2018). Specifically, these are the final cell-type assignments incorporating automated and manual assignment; see also Table S1 in Cusanovich et al (2018). These cell-type assignments were retrieved from the "cell_label" column in the metadata file. The "other immune" label includes cells labeled as B cells, NK cells, T cells, dendritic cells, macrophages and monocytes.

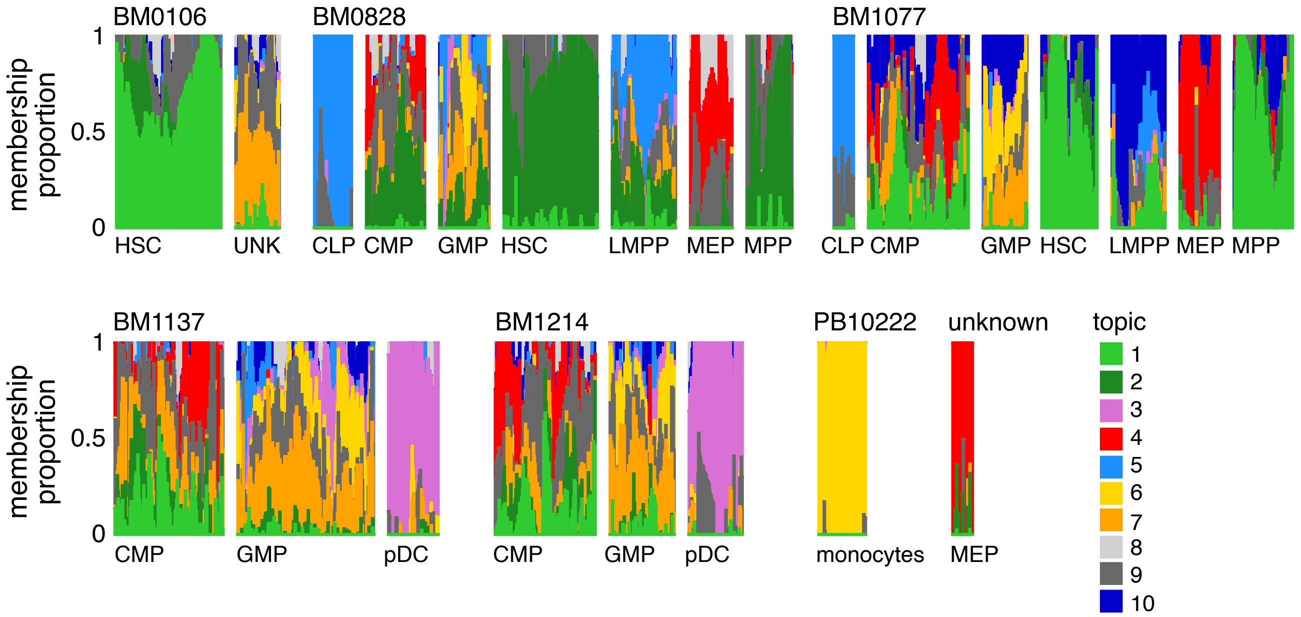


**Figure S11.** Topic model shown in Figure 9A in which the 2,034 cells are arranged by both donor ID and FACS cell-type classification. Topic 2 predominantly occurs in donor BM0828, and topic 10 predominantly occurs in donor BM1077.

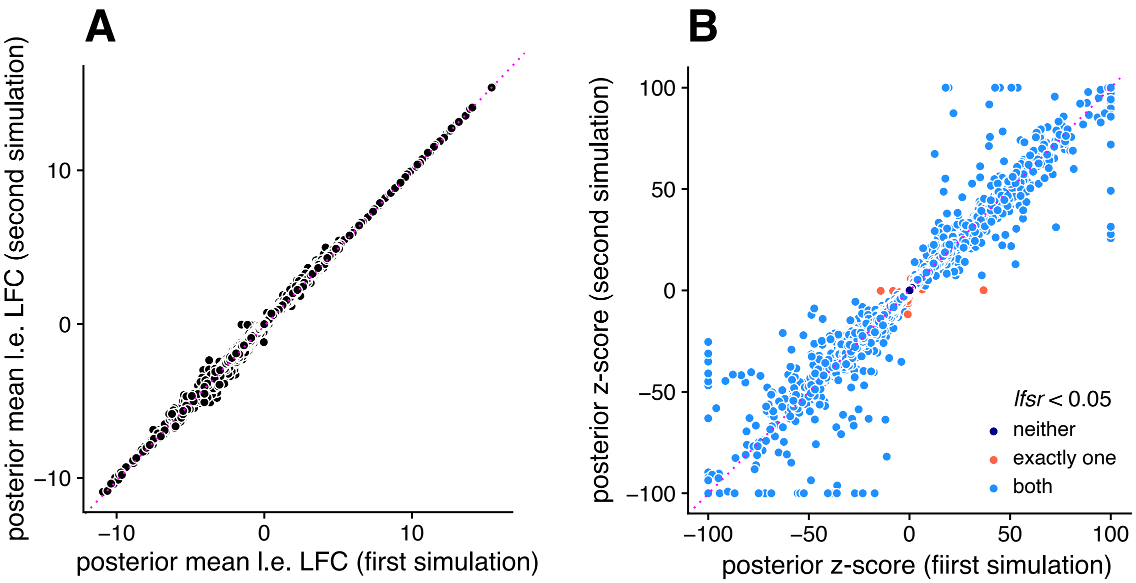


**Figure S12.** Accuracy assessment of MCMC computations in PBMC data. To assess accuracy of the MCMC computations for the DE analysis, we performed two MCMC simulations with different random initializations (100,000 states were simulated in both MCMC runs). Panel A compares the posterior mean l.e. LFC estimates obtained from each run after performing adaptive shrinkage. Panel B compares the estimated posterior *z*-scores (again after performing adaptive shrinkage). Although the posterior *z*-scores are estimated less consistently than the posterior means, the posterior *z*-scores are "accurate enough" in the sense that it is rare for an LFC to have an *lfsr* less than 0.05 in one MCMC simulation and not in the other (the red points depict the LFCs with *lfsr* estimated inconsistently between the two runs). For better visualization of the posterior *z*-scores, posterior *z*-scores larger than 100 (or smaller than -100) are shown as 100 (or -100).

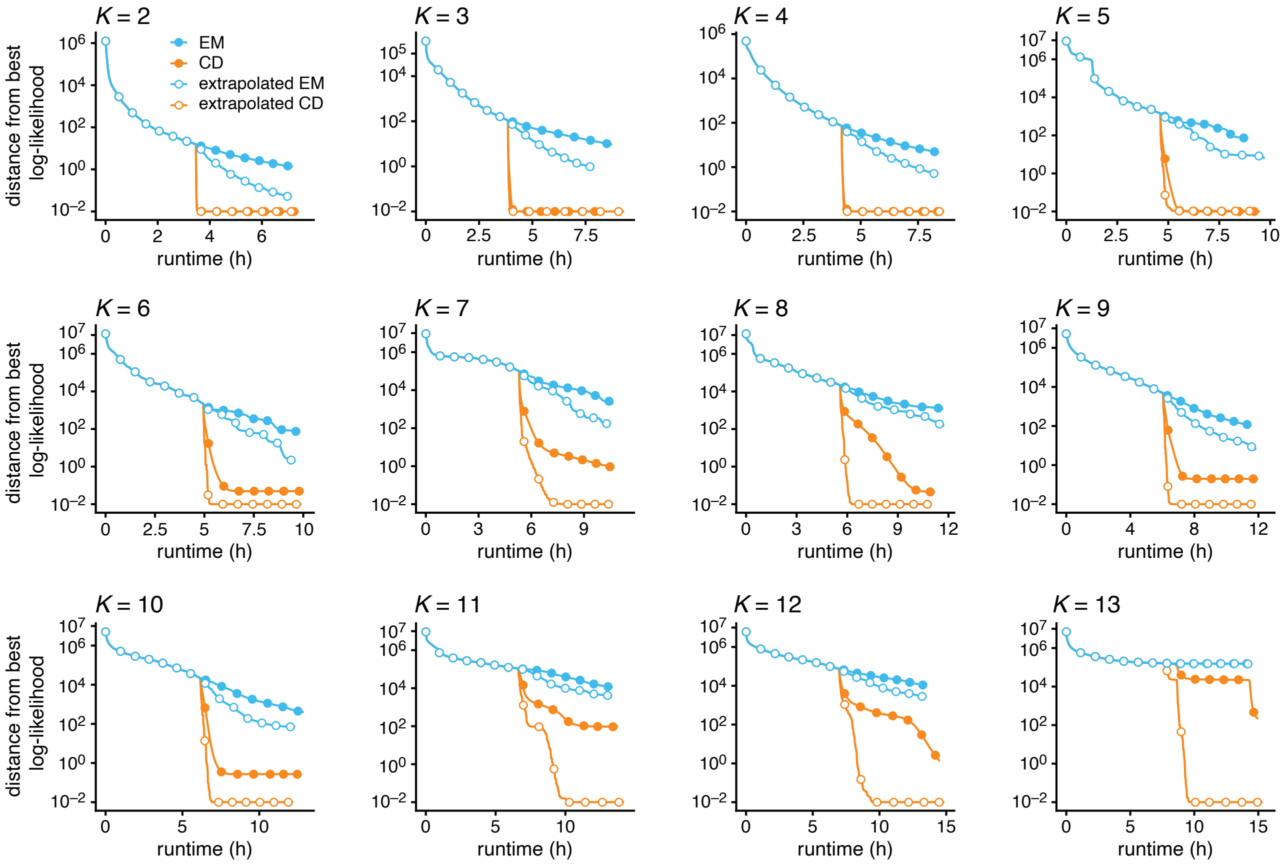


**Figure S13.** Improvement in model fit over time for the Poisson NMF optimization algorithms applied to the PBMC data, with $K = 2, \ldots, 13$. Poisson NMF log-likelihoods are shown relative to the best log-likelihood recovered among the four algorithms compared (EM and CD, with and without extrapolation). All runs began with a "pre-fitting" stage in which 1,000 EM updates were performed (with no extrapolation), followed by a "refinement" stage in which 1,000 updates were performed. Log-likelihood differences less than 0.01 are shown as 0.01. Circles are drawn at intervals of 150 iterations. Note on interpreting these plots: Poisson NMF log-likelihood differences shown in these plots are nearly identical to (multinomial) topic model log-likelihood differences. This is because the Poisson NMF likelihoods are equal to the topic model likelihoods up to a proportionality constant so long as the size factors $s_i$ are constant, and in practice the size factors do not change much over time because the optimization algorithms quickly converge to MLEs of the size factors.

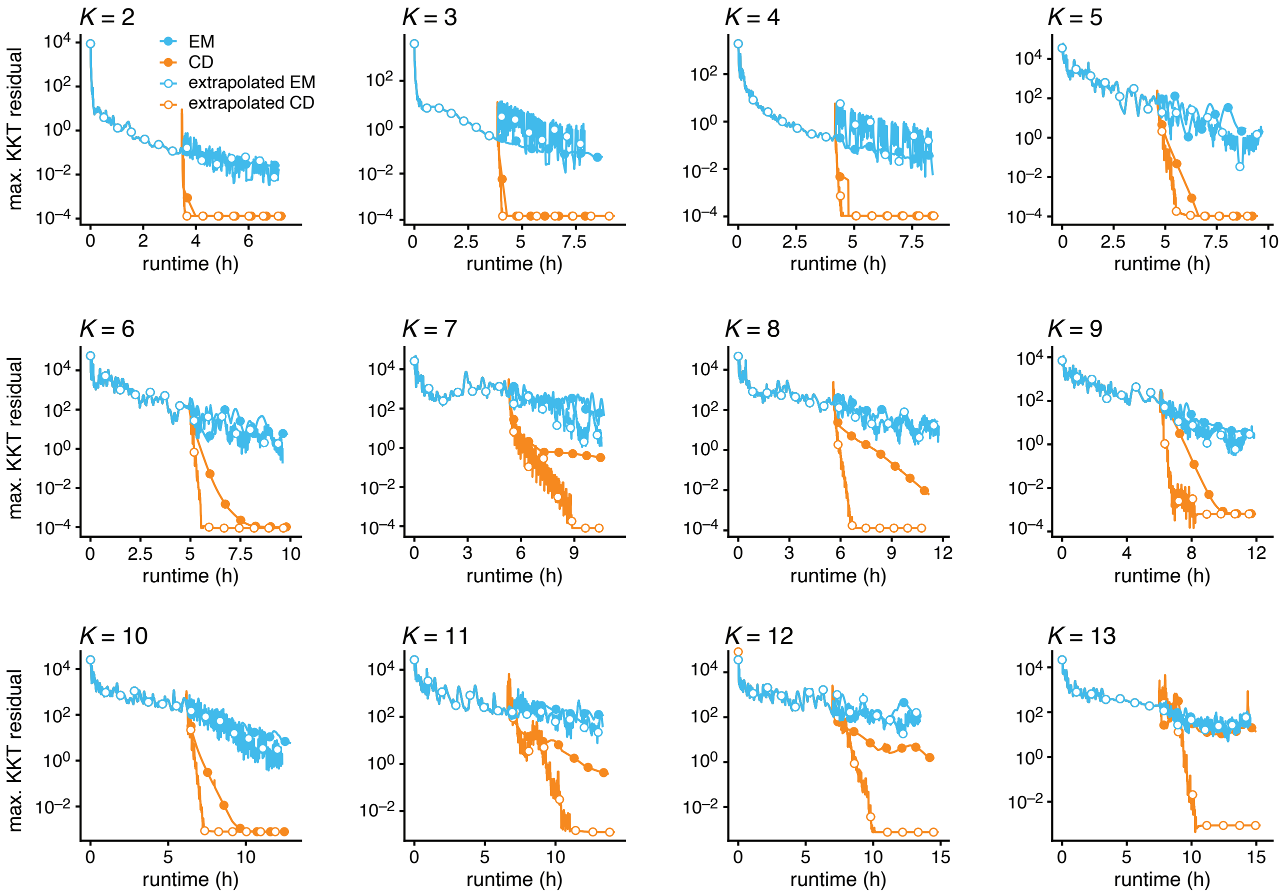


**Figure S14.** Evolution of the KKT residuals over time for the Poisson NMF optimization algorithms applied to the PBMC data, with $K = 2, \dots, 13$. All runs began with a "pre-fitting" stage in which 1,000 EM updates are performed (with no extrapolation), followed by a "refinement" stage in which 1,000 updates were performed. Circles are drawn at intervals of 150 iterations. The KKT residuals should vanish near a local maximum of the likelihood, so looking at the largest KKT residual can be used to assess how well the algorithm recovers an MLE for the Poisson NMF or (multinomial) topic model. Note that, unlike the likelihood, the KKT residuals are not expected to decrease monotonically over time.

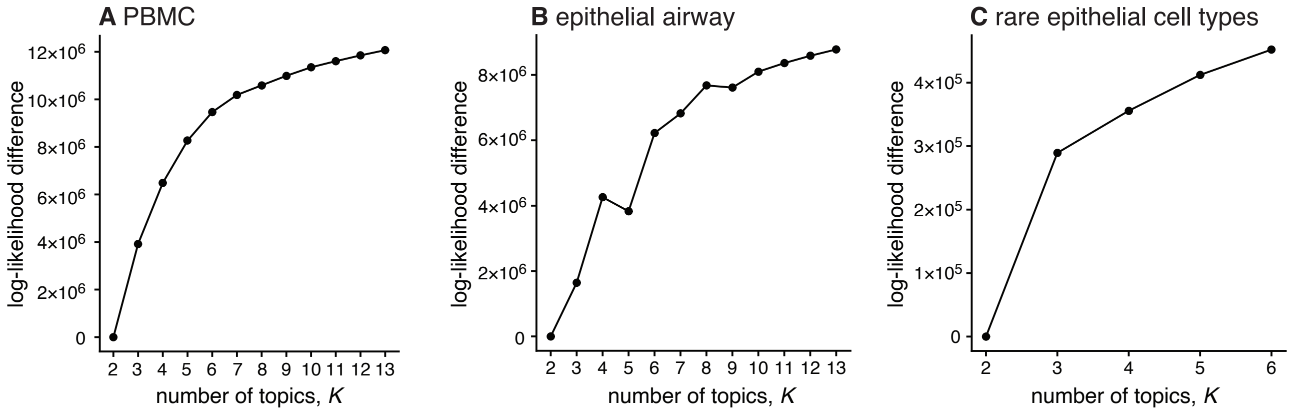


**Figure S15.** Likelihood vs. $K$ in (A) PBMC, (B) epithelial airway data, and (C) the rare cell type subset of epithelial airway cells. These plots shows topic model likelihoods for different choices of the number of topics, $K$. Log-likelihoods are shown relative to the log-likelihood at $K = 2$. Always the best likelihood is shown from the four optimization algorithms compared (EM and CD, with and without extrapolation). Note that these log-likelihood differences can either be interpreted as log-likelihood differences between two Poisson NMF model fits or between two topic model fits since the differences will be identical between two MLEs.

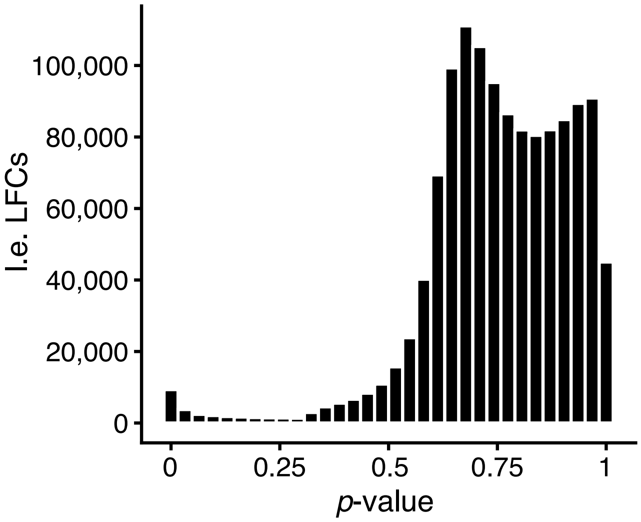


**Figure S16.** Distribution of *p*-values for all topics from the GoM DE analysis of chromatin accessibility using the $K$ = 10 topic model fitted to the hematopoietic system data. Note there is one LFC for each of the chromatin accessibility regions and for each topic. Also note that these *p*-values are not adjusted for multiple testing.
